# Polyclonal origins of human premalignant colorectal lesions

**DOI:** 10.1101/2025.09.08.674484

**Authors:** Debra Van Egeren, Ryan O. Schenck, Aziz Khan, Aaron M Horning, Shanlan Mo, Clemens L. Weiß, Edward D. Esplin, Winston R. Becker, Si Wu, Casey Hanson, Nasim Barapour, Lihua Jiang, Kévin Contrepois, Hayan Lee, Stephanie A. Nevins, Tuhin K. Guha, Hao Zhang, Zhen He, Zhicheng Ma, Emma Monte, Thomas V. Karathanos, Rozelle Laquindanum, Meredith A. Mills, Hassan Chaib, Roxanne Chiu, Ruiqi Jian, Joanne Chan, Mathew Ellenberger, Bahareh Bahmani, Basil Michael, Annika K. Weimer, D. Glen Esplin, Samuel Lancaster, Jeanne Shen, Uri Ladabaum, Teri A. Longacre, Anshul Kundaje, William J. Greenleaf, Zheng Hu, James M. Ford, Michael P. Snyder, Christina Curtis

## Abstract

Cancer is generally thought to be caused by expansion of a single mutant cell^1^. However, analyses of early colorectal cancer lesions suggest that tumors may instead originate from multiple, genetically distinct cell populations^2,3^. Detecting polyclonal tumor initiation is challenging in patients, as it requires profiling early-stage lesions before clonal sweeps obscure diversity. To investigate this, we analyzed normal colorectal mucosa, benign and dysplastic premalignant polyps, and malignant adenocarcinomas (123 samples) from six individuals with familial adenomatous polyposis (FAP). Individuals with FAP have a germline heterozygous *APC* mutation, predisposing them to colorectal cancer and numerous premalignant polyps by early adulthood^4^.

Whole-genome and/or whole-exome sequencing revealed that many premalignant polyps—40% with benign histology and 28% with dysplasia—were composed of multiple genetic lineages that diverged early, consistent with polyclonal origins. This conclusion was reinforced by whole-genome sequencing of single crypts from multiple polyps in additional patients which showed limited sharing of mutations among crypts within the same lesion. In some cases, multiple distinct *APC* mutations co-existed in different lineages of a single polyp, consistent with polyclonality. These findings reshape our understanding of early neoplastic events, demonstrating that tumor initiation can arise from the convergence of diverse mutant clones. They also suggest that cell-intrinsic growth advantages alone may not fully explain tumor initiation, highlighting the importance of microenvironmental and tissue-level factors in early cancer evolution.

## INTRODUCTION

Cancer begins when cells acquire novel malignant traits that increase cell growth, survival, and invasion^5^. Investigating the earliest steps in tumor initiation has important implications for understanding cancer biology, improving risk stratification, and prevention^6,7^. However, studying this process in human tissues is challenging, since phenotypic changes and evolutionary progression make it difficult to characterize the cells that originate the lesion when sampling only malignant tumors. Premalignant lesions are therefore a crucial window into these earlier initiating events, since they reflect an important, often long, period of molecular and evolutionary change where cells acquire the necessary traits for full-blown malignancy^8,9^.

Therefore, to study the initiation of colorectal tumors, we collected and profiled numerous premalignant colon polyps from multiple individuals with familial adenomatous polyposis (FAP), a hereditary cancer predisposition syndrome caused by a germline heterozygous loss-of-function mutation in the *APC* gene^4^. Starting in adolescence, individuals with FAP develop many colorectal polyps, some of which will inevitably progress to colorectal cancer (CRC) without prophylactic colectomy^4^. The abundant polyps in these individuals are an excellent model for studying early tumor progression, since they arise at different times and retain information about the initial state of the lesion that is easily obscured by later selective sweeps. We sampled the entire process of CRC development, in which normal mucosal epithelium can form premalignant polyps, some of which later may develop dysplastic features and then transform into malignant adenocarcinomas (AdCas).

Colorectal polyps are canonically assumed to develop after an epithelial cell acquires the required oncogenic mutation(s) to clonally expand and create a detectable lesion composed of the descendants of the original initiating mutant cell^10–12^. However, some studies of human premalignant polyps challenge this hypothesis. First, others have found that key early driver mutations (*APC* and *KRAS*) are sometimes not found in all cells in a polyp, as would be expected if they initiated a clonal expansion^13^. Instead, there can be multiple unique subclonal mutations in *APC* or *KRAS* in a single polyp^13^. Second, limited in situ hybridization and single-crypt sequencing data have suggested that polyps can be composed of epithelial cells from different genetic lineages that diverged early in the patient’s lifetime, before the polyp initiated^2,3^. Both observations suggest that polyp initiation is not always monoclonal, or the result of clonal expansion of a single cell. Instead, colorectal polyp initiation may often be polyclonal, or the result of expansion of a group of genetically diverse cells, possibly driven by cell-extrinsic effects.

To evaluate whether polyclonal initiation is evident in FAP patient colorectal premalignant colon polyps, to determine its prevalence, and to investigate the role of somatic driver mutations in CRC development, we performed an evolutionary analysis using whole genome sequencing (WGS) and/or whole exome sequencing (WES) data from 123total samples taken either from normal mucosa, benign polyps without evidence of dysplasia, or dysplastic premalignant polyps, or malignant AdCas from 6 patients with FAP. The profiled lesions were of varied age, size and physical location in the colon, and represent an important resource within the NIH Human Tumor Atlas Network (HTAN)^14^ for understanding premalignant progression in CRC. This bulk genomic profiling was used to detect spontaneously-occurring somatic mutations^15–18^, which are a record of past mutational processes and evolutionary dynamics^19–22^. We computationally analyzed these bulk sequencing data to infer clonal architecture and dynamics without longitudinal sampling.

We found that 40% of benign and 28% of dysplastic polyps had evidence of polyclonal initiation. Polyclonal samples were less likely to have clonal *APC* or *KRAS* driver mutations, suggesting that these oncogenic mutations do not always drive monoclonal expansions that lead to tumor initiation. Single-crypt WGS of 3 polyps and 1 AdCa from two additional individuals with FAP suggests the polyps are often polyclonal. Two of these polyps show early genetic divergence of individual crypts, including one polyp with multiple unique *APC* second hit mutations acquired after lesion initiation, consistent with polyclonal initiation. Taken together, our results suggest that polyclonal initiation of premalignant polyps is common, challenging the classic model of monoclonal initiation.

## RESULTS

### FAP polyps have canonical CRC drivers

To investigate early events in CRC lesion initiation, we performed WGS on fresh frozen histologically normal mucosal tissue and multiple benign and dysplastic polyps from 6 individuals with FAP (**Fig. 1a, Supplementary Tables 1-3, Methods, Supplementary Fig. 1**). Five of these individuals had a germline truncating heterozygous mutation in *APC* detected in all tissue samples and blood (**Supplementary Figs. 2-6**), while A014 had no detectable germline mutation in *APC* as assessed by WGS or clinical diagnostic genetic testing. Since this individual has no family history of FAP (**Supplementary Table 1**), this finding is consistent with reports that approximately half of index cases have no pathogenic variants detectable in the blood, and up to 20% of index cases with a detectable *APC* mutation have a somatic mosaic variant that might not be detected in all tissues^23,24^.

**Figure 1.**
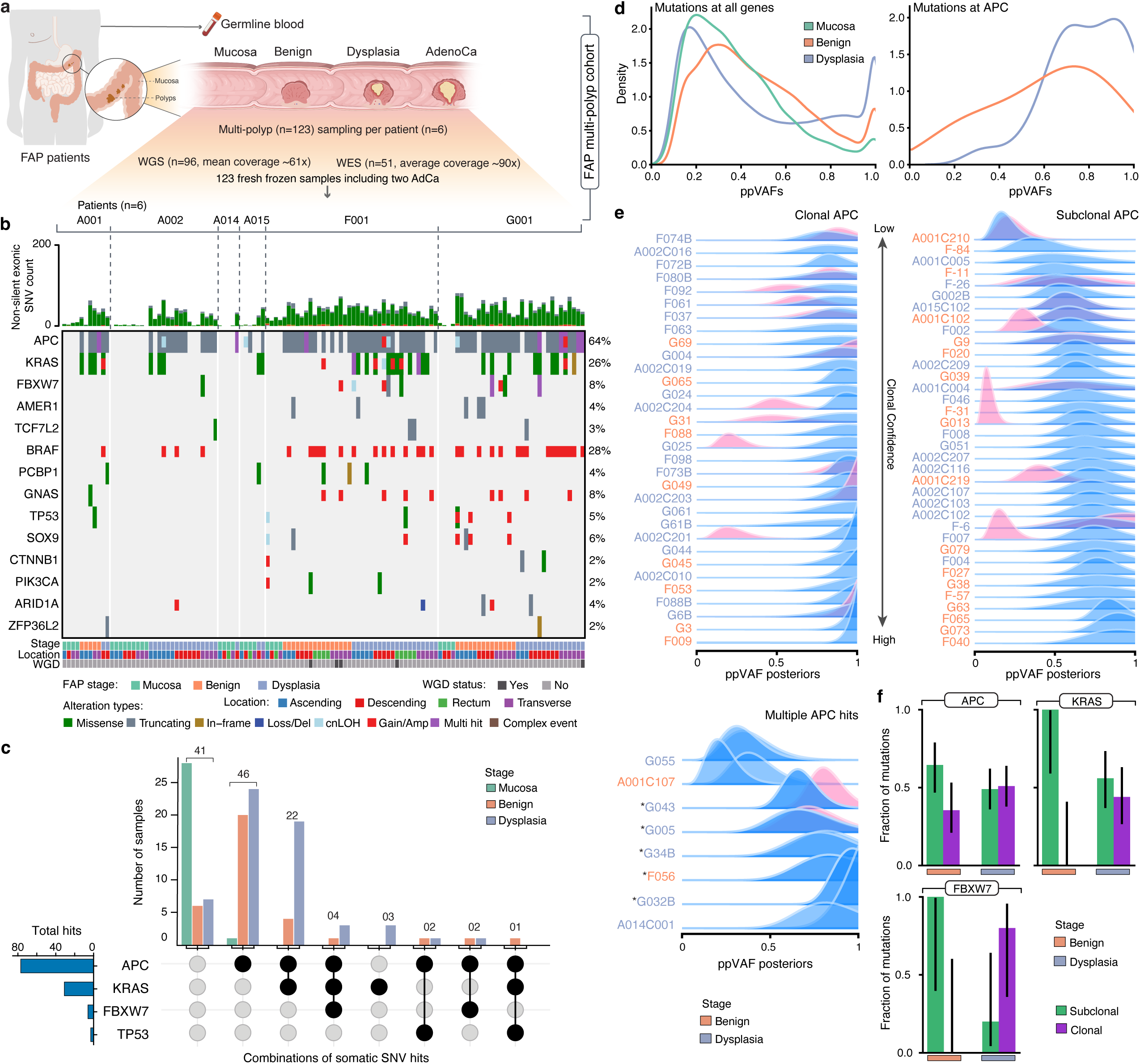
Colorectal polyps in FAP patients harbor common CRC driver mutations. **a.** Overview of the FAP cohort (6 patients, n=123 samples, WGS and/or WES). **b.** Oncoplot summarizing the landscape of non-silent single nucleotide variants (SNVs), small insertions/deletions, copy number gains/amplifications (Gain/Amp), deep deletions (Loss/Del), and copy neutral loss of heterozygosity (cnLOH) within CRC driver genes based on WGS when available or WES. Only somatic mutations are shown. **c.** UpSet plot showing the combination of somatic mutations in *APC*, *KRAS*, *FBXW7* and *TP53*. **d.** ppVAF distributions for all mutations (left) and *APC* second-hit driver mutations (right) in mucosa (green), benign polyps (orange), and dysplastic polyps (blue) from WGS data. **e.** ppVAF posterior distributions of *APC* mutations in samples with a single *APC* second-hit mutation (left, separated into clonal or subclonal *APC* mutations) or multiple *APC* 2nd hit driver mutations (right), ordered by ascending ppVAF point estimates. Asterisks denote clustered *APC* 2nd hit mutations (**Supplementary** Fig. 7). *APC* second-hit mutation ppVAF distributions are shown in blue and *KRAS* driver mutation ppVAF posterior distributions are shown in pink. WGS was used in samples for which it was available, otherwise WES data were used to estimate ppVAFs. **f.** Clonal driver mutation fractions (**Methods**) from WGS (when available) or WES data for benign and dysplastic polyps showing mutations in *APC* (benign: n = 31; dysplastic n = 53 mutations), *KRAS* (benign: n = 6; dysplastic n = 25 mutations), and *FBXW7* (benign: n = 3; dysplastic n = 5 mutations). Error bars are 95% Bayesian credible intervals (**Methods**). Graphics in **a** are adapted from BioRender.

Somatic alterations, including single nucleotide variants (SNVs) and copy number alterations (CNAs), in driver genes associated with CRC were detected in nearly all polyp samples (91% of benign and 95% of dysplastic polyps), but rarely in normal mucosa (10%) (**Fig. 1b**). Driver alterations in *APC* and *KRAS* were common in polyps; 83% of benign polyps, and 82% of dysplastic polyps (adenomas) had second hit *APC* driver mutations or *APC* loss of heterozygosity (LOH), while only 7% of normal mucosal samples had biallelic *APC* inactivation. Furthermore, 17% of benign and 35% of dysplastic polyps had both *APC* and *KRAS* somatic mutations (**Fig. 1c**). The high prevalence of *APC* and *KRAS* somatic mutations in polyp samples is consistent with previous reports in both FAP and sporadic adenomas^25,26^ (**Extended Data Figs. 1, 2**) and suggests that these mutations are associated with polyp initiation. Additional CRC driver mutations were found in *FBXW7, TP53*, and other proliferative signaling associated genes in some lesions (**Fig. 1b**). While recurrent gains of chromosomes 7, 13, and 20 were noted, as observed in other premalignant colorectal lesions^25,26^, polyps were generally less aneuploid than malignant CRC samples (**Extended Data Fig. 3**). The fraction of genome altered (FGA) was low in both benign (median 0.003) and dysplastic (median 0.03) polyps, and increased with disease stage in both our FAP cohort and a previously published FAP^25^ and sporadic cohort^26^ (**Extended Data Fig. 3**). Thus, FAP polyps harbor early oncogenic events, most frequently somatic SNVs, and more rarely CNAs.

### Clonal drivers are often absent in FAP polyps

To better assess the role of driver mutations in polyp initiation and growth, we computed the purity-and ploidy-adjusted variant allele frequencies (ppVAFs) of all somatic mutations detected (**Methods**). The ppVAF is the estimated fraction of epithelial cells that have a mutation and is analogous to the cancer cell fraction (CCF) in malignant samples^27^. While the CCF is usually calculated using tumor purity values estimated from CNA frequencies^28^, our normal and polyp samples have relatively few CNAs compared to CRCs (**Extended Data Fig. 3**), making copy number based purity estimation difficult. Instead, we used the epithelial cell fractions previously measured and reported in additional samples from this patient cohort using scATAC-seq^29^ to estimate the distribution of sample purities for normal mucosa, polyps, and AdCas (**Methods, Supplementary Notes 1-2, Extended Data Fig. 4**). The ppVAF point estimates and posterior probability distributions indicate that somatic *APC* mutations are found at higher allele frequencies than other somatic mutations, reinforcing the idea that they occur early in polyp development and/or experience positive selection (**Fig. 1d**).

However, many polyp samples in our cohort lacked clonal driver mutations (**Fig. 1e-f**). This observation is not consistent with the hypothesis that *APC* second hit mutations or other driver mutations are required for monoclonal expansions leading to polyp formation. Furthermore, 8 of the 69 polyps harbored more than one *APC* somatic driver mutation (**Fig. 1e**). While 5 represent clustered mutational events that occurred in the same subclone (**Supplementary Fig. 7**) or probable biallelic loss in a single subclone in the patient without a germline *APC* mutation (A014), two samples (A001C107 and G055) show evidence of two subclonal *APC* mutations. Since a strong selective advantage for multiple truncating *APC* mutations to appear in the same subclone with a germline heterozygous mutation is unlikely, this observation suggests that multiple unique *APC* mutations existed in the epithelial cell population that initiated the lesion and/or additional *APC* mutations were acquired by cells in the lesion that did not have an *APC* second hit at the time of lesion initiation. Together, these results suggest that polyps are not always caused by a monoclonal expansion of an epithelial cell with a driver mutation.

### Polyps have subclones that diverged early

Since we observed that polyp initiation is not always associated with monoclonal expansion of one or more driver mutations, we wondered how lesion initiation occurred in samples without clonal driver mutations. We hypothesized that multiple colonic crypts (clonal units of epithelial homeostasis derived from a small number of stem cells^30^) might collectively initiate each polyp, rather than a single mutant crypt. This would lead to a polyp composed of multiple distinct genetic lineages that diverged early. We defined initiation of a single polyp from multiple crypts as “polyclonal” and used the WGS data to determine whether polyp initiation was likely monoclonal or polyclonal (**Fig. 2a**). In monoclonal samples, clonal expansion of a single crypt leads to all somatic mutations in that founding crypt being present in all epithelial cells in the resulting lesion. This sweep leads to many detectable clonal mutations in the sequenced polyp. In contrast, polyclonal lesions will likely have fewer clonal mutations, since any clonal mutations in the lesion must be present in all founding crypts in a polyp (**Fig. 2b**).

**Figure 2.**
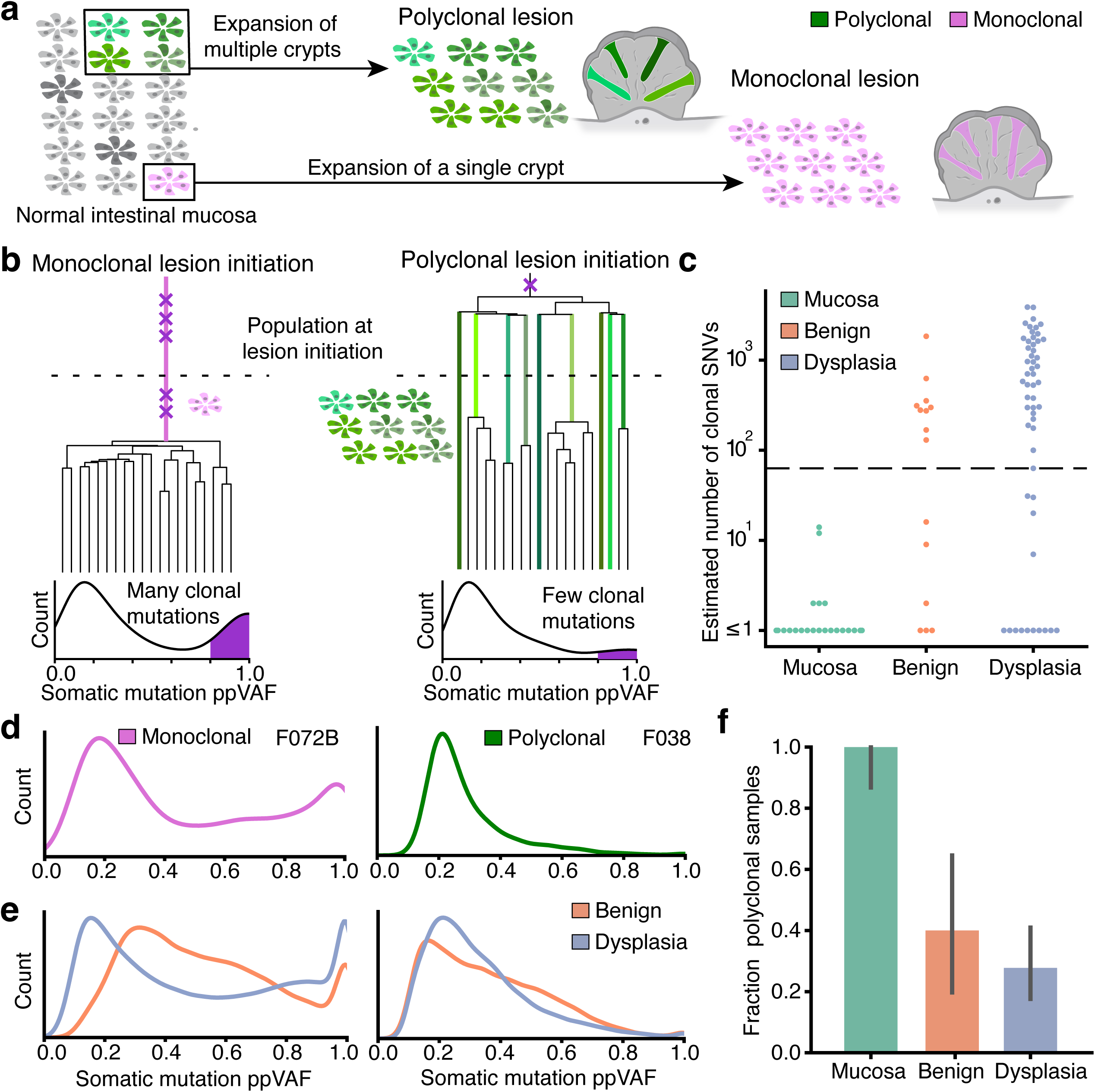
Some FAP premalignant polyps had an early most recent common evolutionary ancestor and are likely polyclonal. **a.** Schematic showing that polyclonal polyps are initiated from multiple colorectal crypts in the intestinal mucosa (green crypts), while monoclonal polyps are the result of a clonal expansion of a single crypt (pink). **b**. Schematic showing that monoclonal lesions have a most recent common ancestor occurring after lesion initiation (dotted line), since they initiate from a single crypt, leading to many detected clonal mutations (purple Xs on tree). Bottom panels show hypothetical ppVAF distribution shapes for monoclonal and polyclonal samples, with clonal mutations marked in purple. **c.** Number of expected clonal SNVs detected in WGS data from FAP patient samples. Samples with fewer than 63 clonal SNVs (dashed line, corresponding to an MRCA at < 1 year old) were classified as having an early MRCA and are likely polyclonal. **d.** ppVAF distributions for example monoclonal (left) and polyclonal (right) lesions. **e.** ppVAF distributions for all monoclonal (left) and polyclonal (right) lesions combined. Colors as indicated in panels **c, f. f.** Fraction of WGS samples classified as polyclonal based on estimated clonal SNV count. Fractions were estimated from n = 25 mucosal samples, n = 15 benign polyp samples, and n = 54 dysplastic polyp samples. Error bars are 95% Bayesian credible intervals (**Methods**). Graphics in **a** are adapted from BioRender.

We estimated the number of clonal single nucleotide variants (SNVs) in each sample using the ppVAF posterior probability distributions computed for each somatic mutation from the sequencing data. We classified each somatic SNV as clonal or subclonal based on its ppVAF posterior probability distribution and counted the number of clonal mutations in each sample (**Methods, Supplementary Note 1**). We used these estimated clonal SNV counts to determine whether lesions were likely monoclonal or polyclonal. Since most SNVs in these samples have clock-like mutational signatures^31^ (**Extended Data Fig. 5**), the number of clonal mutations can be used to estimate an upper bound on the age of the most recent common ancestor (MRCA) for the sample (**Methods**). Samples with an MRCA earlier than the time at which the lesion initiated are by definition polyclonal. In polyclonal samples, multiple genetic lineages must have existed during lesion initiation, so these genetic lineages must therefore have diverged from the MRCA before initiation. If the MRCA occurred at or after lesion initiation, the lesion is composed of cells that originated from a single genetic lineage at the time of initiation and is monoclonal. We observed that, while normal mucosal samples had few (<15) clonal SNVs, benign and dysplastic polyps had a much wider range of expected clonal SNV counts (0-3882 SNVs) (**Fig. 2c**). The low clonal mutation count in the normal samples is consistent with previous observations suggesting that normal intestinal mucosa is highly polyclonal, often having few mutations shared between nearby crypts^32^.

To investigate the prevalence of polyclonal polyps, we estimated the frequency of polyps with an MRCA that existed prior to one year of age (< 63 clonal SNVs, using WGS data) (**Methods**), which is far earlier than polyps appear in most individuals with FAP^33^ and is similar to the estimated clonal mutation counts of the polyclonal normal mucosa. We found that 6 out of 15 benign polyps (40%) had an early MRCA, suggesting they are polyclonal, and 15 out of 54 dysplastic polyps (28%) had an early MRCA (**Fig. 2d-f**). Samples called as polyclonal generally do not have a clonal mutation peak visible in the raw VAF or ppVAF distributions (**Fig. 2d-e**, **Supplementary Figs. 8-17**), consistent with their early divergence time. WES data from some samples with WGS data as well as 27 additional samples from our cohort were also subjected to clonal SNV count thresholding to estimate the prevalence of early-diverging samples and showed a similar fraction of these lesions (**Extended Data Fig. 6**). Additionally, clonal SNV counts suggest that polyclonality is prevalent in premalignant and malignant multi-region WES samples from a different published FAP cohort^25^ (**Extended Data Fig. 7, Methods**). A similar analysis using published multi-region WES data from individuals without a hereditary CRC predisposition syndrome^26^ suggests that polyclonal initiation also occurs in sporadic premalignant adenomas but not AdCas (**Extended Data Fig. 7**).

These analyses suggest that polyclonal initiation is likely present in at least 25-40% of polyps in individuals with FAP. Selective sweeps occurring after lesion initiation can obscure a polyclonal population structure, so the true fraction of lesions that initiate polyclonally may be higher. This is particularly true in older lesions or lesions containing driver mutations with a substantial selective advantage, highlighting the importance of studying early premalignant adenomas. Amongst dysplastic polyps, those of monoclonal origin had shorter telomeres than polyclonal lesions (*p*=0.0015; Fisher’s combined p-value from per-patient Wilcoxon rank-sum tests) (**Methods, Extended Data Fig. 8**), consistent with a higher number of total cell divisions due to earlier lesion initiation and/or increased proliferation rate. Early, proliferative lesions would have shorter telomeres when sampled and would be more likely to appear monoclonal, particularly if there were expansion of an advantageous clone.

### Polyclonal polyps have subclonal drivers

We used these monoclonal/polyclonal classifications to further investigate the role of driver mutations in polyp initiation and progression. Most monoclonal and polyclonal polyps have an *APC* second hit driver mutation (**Fig. 3a**). As expected, the ppVAFs of *APC* somatic driver mutations are lower in polyclonal samples than monoclonal samples (**Fig. 3b**; *p*=0.003 for benign, *p*=0.005 for dysplastic polyps, Wilcoxon rank-sum test), suggesting that these mutations are not the sole drivers of lesion initiation and growth and have not caused a selective sweep in the polyclonal polyps. Similarly, *KRAS* mutations are more likely to be found at lower subclonal frequencies in dysplastic polyclonal polyps than dysplastic monoclonal polyps (*p*=0.074 for benign, *p*=0.012 for dysplastic polyps, Wilcoxon rank-sum test, **Fig. 3c-d**). These data are consistent with the notion that premalignant lesions are not always the product of monoclonal expansions associated with common early CRC driver mutations. However, such driver mutations likely have a selective advantage which often increases their frequency and can lead to selective sweeps in more advanced lesions.

**Figure 3.**
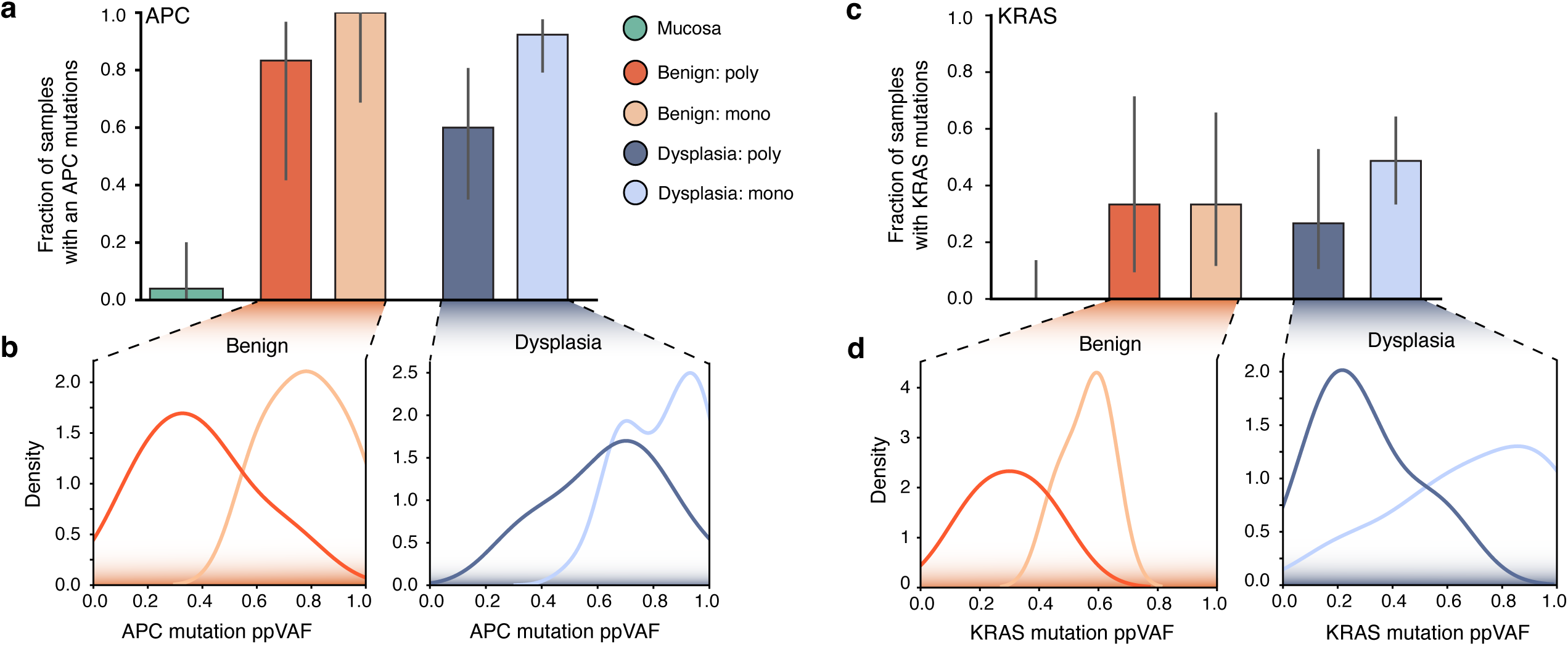
Polyclonal samples had subclonal driver mutations and were not the result of the expansion of an *APC* and/or *KRAS* mutated clone. **a**. Fraction of monoclonal and polyclonal WGS samples with second-hit *APC* somatic driver mutations. **b.** ppVAFs of *APC* second hit mutations in benign (left) and dysplastic (right) polyps. **c.** Fraction of monoclonal and polyclonal WGS samples with *KRAS* driver mutations. **d.** ppVAFs of *KRAS* driver mutations in benign (left) and dysplastic (right) polyps. Error bars in **a** and **c** are 95% Bayesian credible intervals (**Methods**), and fractions in **a** and **c** were estimated from n = 25 mucosal, n = 6 polyclonal benign, n = 9 monoclonal benign, n = 15 polyclonal dysplastic, and n = 39 monoclonal dysplastic samples.

### Single-crypt WGS shows polyclonality

We performed single-crypt isolation and WGS from polyps and AdCas from two additional FAP patients (**Fig. 4, Extended Data Figs. 9-10, Methods, Supplementary Table 4**) to examine the clonal relationships between cell populations within each lesion. FAP patient lesions were dissected into multiple regions before single crypt isolation to better determine the association between spatial proximity and genomic similarity. After filtering out low quality crypts and polyps/tumors, we built phylogenies using single crypt WGS data from the remaining three polyps and one AdCa sample (**Methods, Supplementary Note 3**). We first focused on the polyp for which the most crypts were sequenced (FAP03_P2; 20 single crypts included after filtering). We found that crypts within the polyp diverged early in life, with no mutations shared by all crypts sequenced in the polyp (**Fig. 4c**). This finding suggests that multiple distinct genetic lineages with a very early common ancestor combined to form the polyp. The lack of clonal somatic mutations in this polyp is reflected in the structure of its single-crypt phylogeny (**Fig. 4c**), with long terminal branches and no trunk, as hypothesized in the polyclonal schematic shown in **Fig. 2b**. This high degree of polyclonality appears spatially structured, with one polyp region (R4) appearing monophyletic (**Fig. 4c**), with 1025 mutations shared between and exclusive to all 5 crypts in the region (**Extended Data Fig. 10b**). This implies that the crypts in R4 share a more recent common ancestor than the rest of the polyp, which may have expanded clonally to create that region. In contrast, the other regions are polyclonal mixtures of crypts from different genetic lineages.

**Figure 4.**
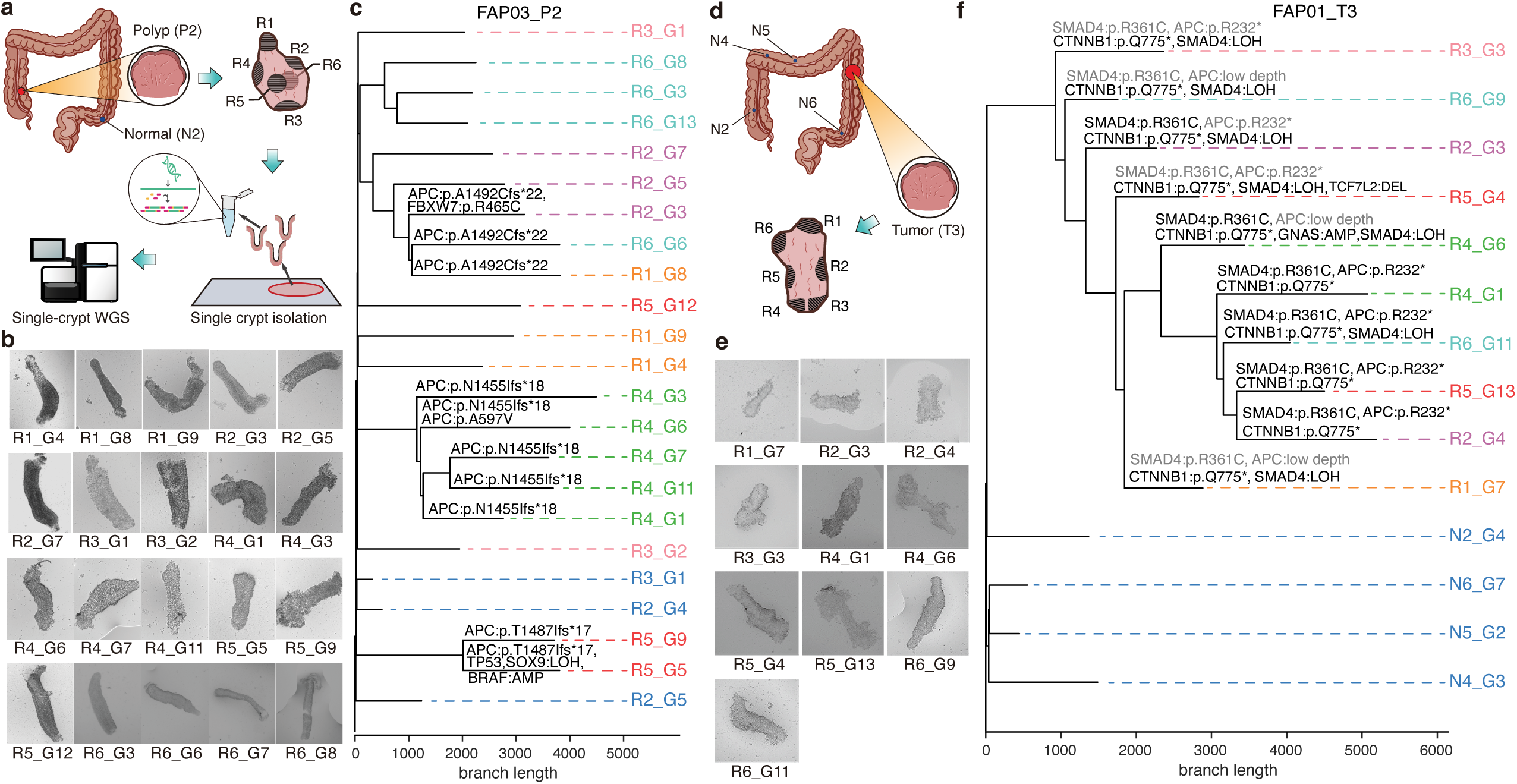
Single-crypt phylogenies based on WGS suggest that FAP polyps are polyclonal, while adenocarcinomas are monoclonal. **a.** Schematic showing collection of polyp P2 from patient FAP03, regional dissection, single crypt isolation, and WGS procedure (**Methods; Supplementary Information)**. **b.** Images of individual isolated crypts from FAP03_P2. **c.** Single crypt phylogeny reconstructed from single-crypt WGS data from polyp P2, patient FAP03 and adjacent normal mucosa (blue). Putative CRC-associated mutations are highlighted. The spatial region from which each crypt originated is indicated by the tip label colors (**Supplementary Table 4**). **d.** Schematic showing collection of CRC lesion T3 from patient FAP01 as well as regional dissection of the lesion. **e.** Images of individual isolated crypts from FAP01_T3. **f.** Single crypt phylogeny reconstructed from lesion T3 from patient FAP01 and normal mucosa (blue). Putative CRC-associated mutations are highlighted, with mutations denoted in grey if they were filtered out of some samples but not others, or if the sequencing depth at the mutated site was too low to detect the variant (**Methods; Supplementary Note 3, Section 5**). The spatial region from which each crypt originated is indicated by the tip label colors. Graphics in **a** and **d** are adapted from BioRender, bioicons (CC BY 4.0) and/or Servier Medical Art (CC BY 4.0).

The single-crypt WGS analysis also revealed putative cancer driver mutations in distinct subpopulations within the polyp. Importantly, no driver mutations are shared by all crypts in the polyp. Instead, there were three truncating frameshift mutations in *APC* occurring in independent subclones (**Fig. 4, Extended Data Fig. 9c**) that either arose after lesion initiation or were present at subclonal frequencies in the initiating cell population. One of these *APC* second hit mutations is found in all crypts in region R4, raising the possibility that it caused a clonal expansion that dominated that part of the polyp. These observations suggest that this polyp did not initiate from a clonal expansion driven by a single *APC* second hit, but rather is a mixture of epithelial cell clones with independent growth advantages.

In contrast, single-crypt WGS of the malignant AdCa tumor sample FAP01_T3 revealed a monoclonal expansion, with 479 mutations shared by all 10 crypts in the sample (**Fig. 4d-f**). This expanded clone includes a second-hit stopgain mutation in *APC*, a missense driver mutation in *SMAD4* (R361C), and a truncating mutation in *CTNNB1* (beta-catenin) (**Fig. 4f**). Lower sequencing depth in some loci in some samples makes it more difficult to determine which driver mutation(s) are found in all crypts and possibly responsible for the initial expansion; in particular, low coverage (≤3x) at the *APC* second-hit mutation locus in crypts R6_G9, R5_G4, and R1_G7 limited our ability to detect the mutation in these samples (**Supplementary Note 3**). However, the phylogenetic structure of this tumor (long trunk from which all tumor crypts originate) clearly suggests it is monoclonal (compare phylogeny to schematics in **Fig. 2c**) and provides a contrasting example to the phylogenetic pattern in the polyclonal polyp FAP03_P2.

Two additional polyps from these patients with single-crypt sequencing data (FAP01_P6 and FAP03_P1) reveal different patterns (**Extended Data Fig. 9**). The 908 clonal mutations in the 7 crypts from polyp FAP01_P6 suggest that it is monoclonal. The lesion has a clonal expansion, possibly resulting from the *APC* LOH event found in all crypts from this polyp and further fueled by a subclonal *KRAS* mutation (Q61R), consistent with the majority of the *APC* second-hit driven monoclonal polyps in our bulk sequencing dataset. In contrast, the single-crypt WGS data from polyp FAP03_P1 are more difficult to interpret. While very few clonal mutations are present (21 shared by all 7 crypts), suggesting early genetic divergence of the crypts within it, the somatic *APC* truncating mutation present in 6 out the 7 crypts (which only share 57 other mutations) raises the possibility that this *APC* second hit occurred very early in life and may have contributed to lesion initiation. In summary, this single-crypt sequencing dataset provides high-resolution orthogonal validation of polyclonal initiation in FAP and highlights the mutational heterogeneity in polyclonal lesions.

## DISCUSSION

While it is widely assumed that malignancies are the product of clonal expansions from single mutant cells of origin^10^, limited case studies in CRC and other cancers indicate that premalignant lesions can be polyclonal^2,3,34,35^. However, systematic assessment of the prevalence of polyclonality in patient samples is still lacking, despite the utility of sequencing to detect this phenomenon. To address this, we used WGS data from 69 colorectal polyps and found 40% with benign histology and 28% with dysplasia from individuals with FAP originated from multiple colon crypts. Furthermore, analysis of single-crypt WGS data supported our conclusion that premalignant colorectal lesions in individuals with FAP can initiate polyclonally, and revealed local expansion of subpopulations within a polyp with unique *APC* second hit mutations. These findings point towards a possible role for cell-extrinsic mechanisms in tumor initiation, and suggest that this process may involve cell-cell interactions in a premalignant multicellular ecosystem as well as cell-intrinsic effects of driver mutations.

The finding that many polyps are polyclonal has implications for understanding the molecular and microenvironmental determinants and dynamics of tumor initiation and progression. Polyclonal initiation provides a genetically-diverse starting point for premalignant evolution. This intralesion heterogeneity can persist since sweeps are relatively rare, with long periods of stasis between such events. The finding that premalignant polyps may experience multiple selective sweeps before accruing genetic alterations and transforming to CRC, is consistent with our previous findings that subsequent evolution is often effectively neutral^19–21^. Common drivers (*APC, KRAS*) may not directly initiate the lesion by causing a clonal expansion but instead may be present at subclonal frequencies within the initiating population or be acquired after lesion initiation. This process can lead to monoclonal conversion of a previously polyclonal lesion and may lead to underestimation of the frequency of polyclonal initiation in our dataset, suggesting that the extent of this phenomenon may be substantially higher than reported here.

Our study has several limitations. First, our bulk sequencing analysis does not directly estimate the purity of each sample individually, but rather uses the distribution of epithelial cell fractions estimated from scATAC-seq data from the HTAN FAP patient cohort as a measure of purity. While this procedure avoids the pitfalls of copy-number based algorithms which render them inappropriate for nonmalignant tissue samples (**Methods**, **Extended Data Fig. 4**, **Supplementary Note 2**), samples with lower purity due to stromal or immune cell inclusion may be falsely called as polyclonal. While this cannot account for all polyclonal samples we identify via bulk genomic sequencing (**Supplementary Note 2**), additional profiling with strategies that either isolate epithelial cells for sequencing (such as single-crypt WGS) or directly measure epithelial cell fraction in the same sample will be instrumental to accurately estimate the fraction of polyclonal samples. Additionally, we focused on CRC initiation in the context of a hereditary cancer predisposition syndrome (FAP). Focus on this patient population also introduces the possibility that some of our polyclonal polyps could be caused by stochastic collision of independently initiated lesions due to the increased density of colorectal polyps in FAP patients. However, recently published studies suggest polyclonality is a general phenomenon in colorectal lesions. Our findings are consistent with studies in the analogous *APC* mutant murine model, which demonstrate cellular cooperativity during colorectal tumorigenesis^35^ and polyclonal tumor origins^36–38^. Moreover, analysis of clonal SNV counts in sporadic CRC adenomas similarly suggests that 29% are polyclonal^36,38^. Thus, polyclonality is unlikely to be restricted to hereditary colon cancers or exclusively caused by random polyp collisions.

Our findings raise questions about the events necessary and/or sufficient for cancer development. Here, we did not identify what mechanistic role these additional epithelial cell clones are playing in tumorigenesis, and follow-up studies will be required to determine how these clones function and interact. Evidence for the role of cell-cell interactions in tumor development between epithelial cells suggests that both cooperation between premalignant clones^40^ and recruitment of neighboring non-malignant epithelium^36,41^ may contribute to polyclonal initiation, though other interactions may also be involved. Furthermore, signals from non-epithelial cells in the microenvironment, including fibroblasts and immune cells^42,43^, may lead to malignant phenotypic changes in multiple colonic crypts at once, resulting in a polyclonal lesion. These findings also raise questions about the role of canonical driver mutations in tumor initiation: how does acquisition of an oncogenic mutation in an individual cell lead to expansion of a diverse group of cells, not all of which have the mutation? More generally, polyclonal initiation may be common across diverse tissue types^34,35^. Indeed, a recent study in premalignant pancreatic cancer lesions demonstrated extreme multi-focality, consistent with polyclonal origins^34^. The conceptual and analytic framework we outline for assessing polyclonality from sequencing data can be extended to other premalignant lesions to systematically investigate this underappreciated phenomenon in cancer initiation.

## Acknowledgements

This work was supported by NCI grant U2CCA233311 to MPS, JMF and CC. AK, CW, DVE, ROS, and CC were additionally supported by NIH grant DP1CA238296 to CC. ROS is supported by a Stanford Dean’s Fellowship. AH was funded in part by the Cancer Systems Biology Scholars Postdoctoral Fellowship Program (5R25CA180993). DVE is supported by the A.P. Giannini Postdoctoral Fellowship. SAN was supported in part by the Stanford Graduate Fellowship in Science and Engineering and the NHGRI Stanford Genome Training Program (5 T32 000044). This work is also supported by NIH S10OD025212. We thank Alvina Adimoelja, Kathleen Houlahan, Seongyeol Park and other members of the Curtis Lab for feedback on the manuscript. This paper is dedicated to FF.

## Contributions

Conceptualization: AH, AK, CC, DVE, ROS

Bioinformatic Analysis: AH, AK, BB, CW, DVE, ROS, SL, SMo, ZhHu

Statistical Analyses: AK, CW, DVE, ROS

Experimental Methods/Data Generation: AH, EDE, HZ, ZM, ZhHu, ZhHe, SMo

Guidance on Methods: AKu, WG

Sample Collection: JF, MEM, RL, SAN, TKG, UL, WRB

Pathology Review: DGE, JC, JS, TL

Supervision: CC, JF, MPS

Writing – original draft: AH, AK, CC, DVE, ROS

Writing – final version: CC, DVE

## Data Availability

DNA sequencing data and metadata have been deposited at the HTAN portal: https://www.ncbi.nlm.nih.gov/projects/gap/cgi-bin/study.cgi?study_id=phs002371.v3.p1. The raw single-crypt WGS data have been deposited in NGDC’s Genome Sequence Archive (GSA) under the accession number HRA009036 (https://ngdc.cncb.ac.cn/gsa-human/browse/HRA009036). Processed source data and resources needed for reproducing key analyses are available through Zenodo (https://doi.org/10.5281/zenodo.13228021).

The whole-exome sequencing data for the sporadic CRC cohort are available through the European Genome-Phenome Archive (EGA) under accession number EGAS00001003066 (https://ega-archive.org/studies/EGAS00001003066). The whole-exome sequencing data for the FAP multi-region cohort are available through the Genome Sequence Archive (GSA) under accession number HRA000127 (https://ngdc.cncb.ac.cn/gsa-human/browse/HRA000127). To facilitate comparisons, both of these published cohorts were uniformly reprocessed using the same software versions within the Isabl platform, as was used for the HTAN Pre-cancer Atlas bulk WGS FAP cohort.

## Code Availability

Source code and source data to generate the figures is available via GitHub: https://github.com/cancersysbio/HTAN-FAP and an archived copy of the code is available on Zenodo (https://doi.org/10.5281/zenodo.17372231).

## Methods

See Supplementary Information

## Supplementary Information

This file contains Methods, Supplementary Figures 1-17 and legends, and Supplementary Notes 1-3.

---

**Extended Data Figure 1.**
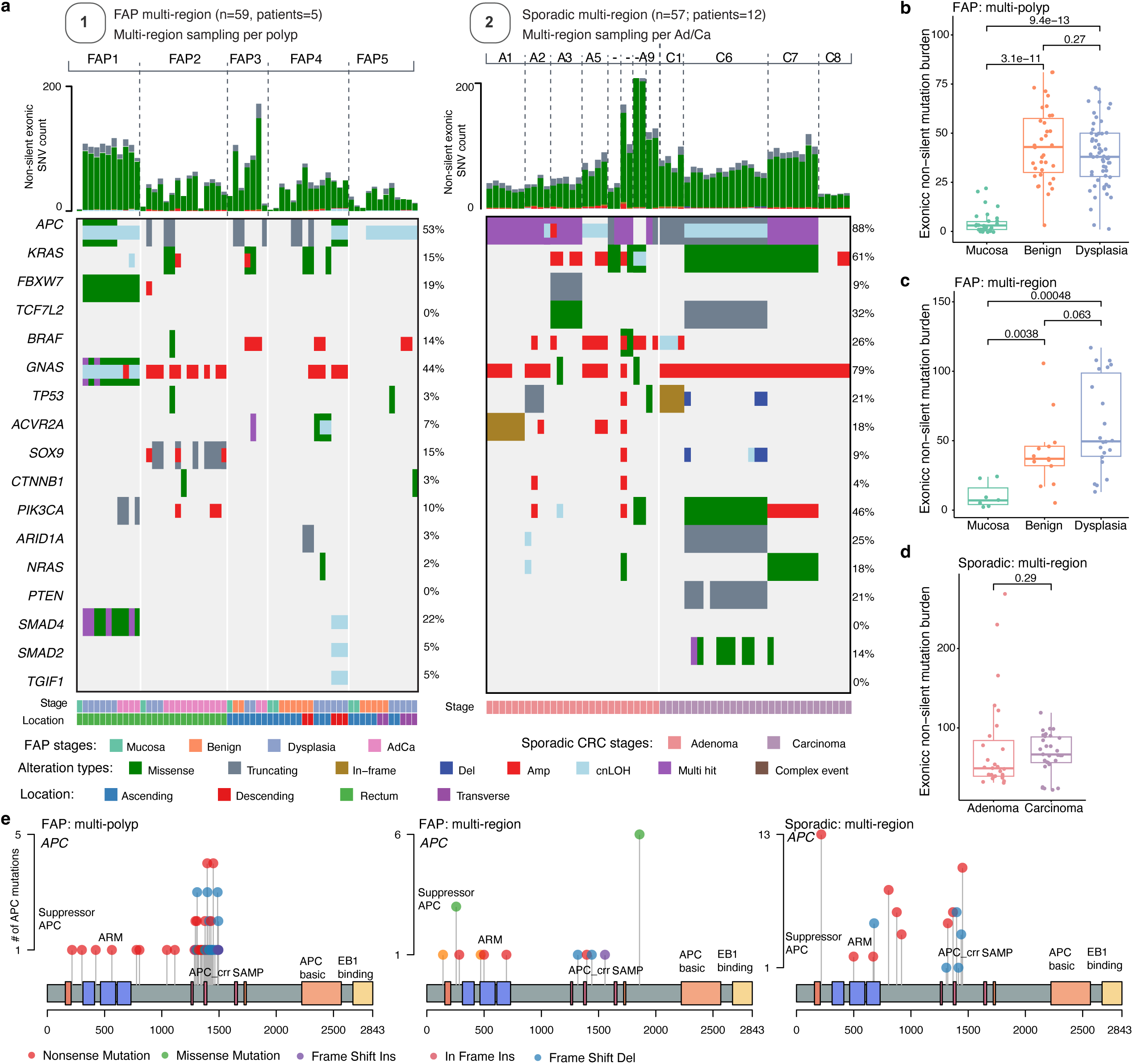
Somatic mutations in FAP and sporadic CRC samples. **a.** Oncoplot summarizing the landscape of non-silent exonic single nucleotide variants (SNVs), small insertions/deletions, copy number amplifications (Amp), deletions (Del), and copy neutral loss of heterozygosity (cnLOH) within CRC driver genes based on WES data from the previously published FAP multi-region cohort^25^ (left) and the sporadic multi-region sequencing cohort^26^ (right). Only somatic mutations are shown. **b-d.** Comparison of non-silent exonic mutations across polyp stages between the two FAP cohorts and the CRC cohort (p-values are estimated using two-sided Wilcoxon rank sum tests). In box-and-whisker plots, the box represents the interquartile range (IQR) with the center line representing the median, and the whiskers are the largest and smallest data values within 1.5 times the IQR from the box edges. Each point represents one sequenced normal or polyp sample, with n = 29 mucosal, n = 35 benign, and n = 57 dysplastic samples in the multi-polyp HTAN FAP cohort (**b**), n = 7 mucosal, n = 13 benign, and n = 22 dysplastic samples in the multi-region FAP cohort (**c**), and n = 27 adenoma and n = 30 carcinoma samples in the sporadic CRC cohort (**d**). **e.** Lollipop plots showing the distribution and classes of mutations in *APC* across two FAP cohorts and one sporadic CRC cohort.

**Extended Data Figure 2.**
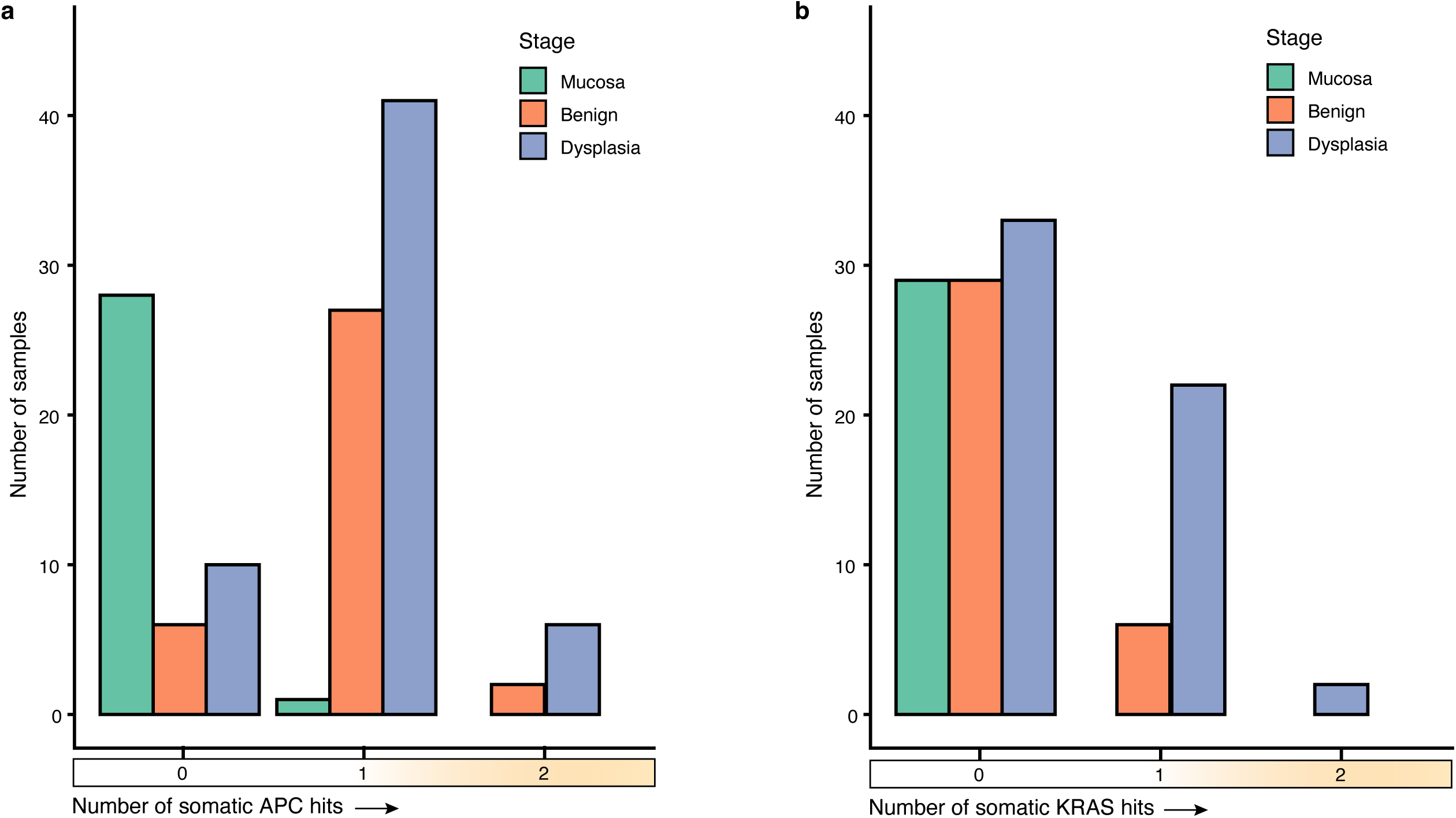
Distribution of *APC* and *KRAS* somatic mutations in FAP polyps and normal mucosa. a. Number of normal mucosa, benign and dysplastic polyps with 0, 1, or 2 somatic APC mutations (‘hits’). b. Number of normal mucosa, benign and dysplastic polyps with 0, 1, or 2 somatic *KRAS* mutations.

**Extended Data Figure 3.**
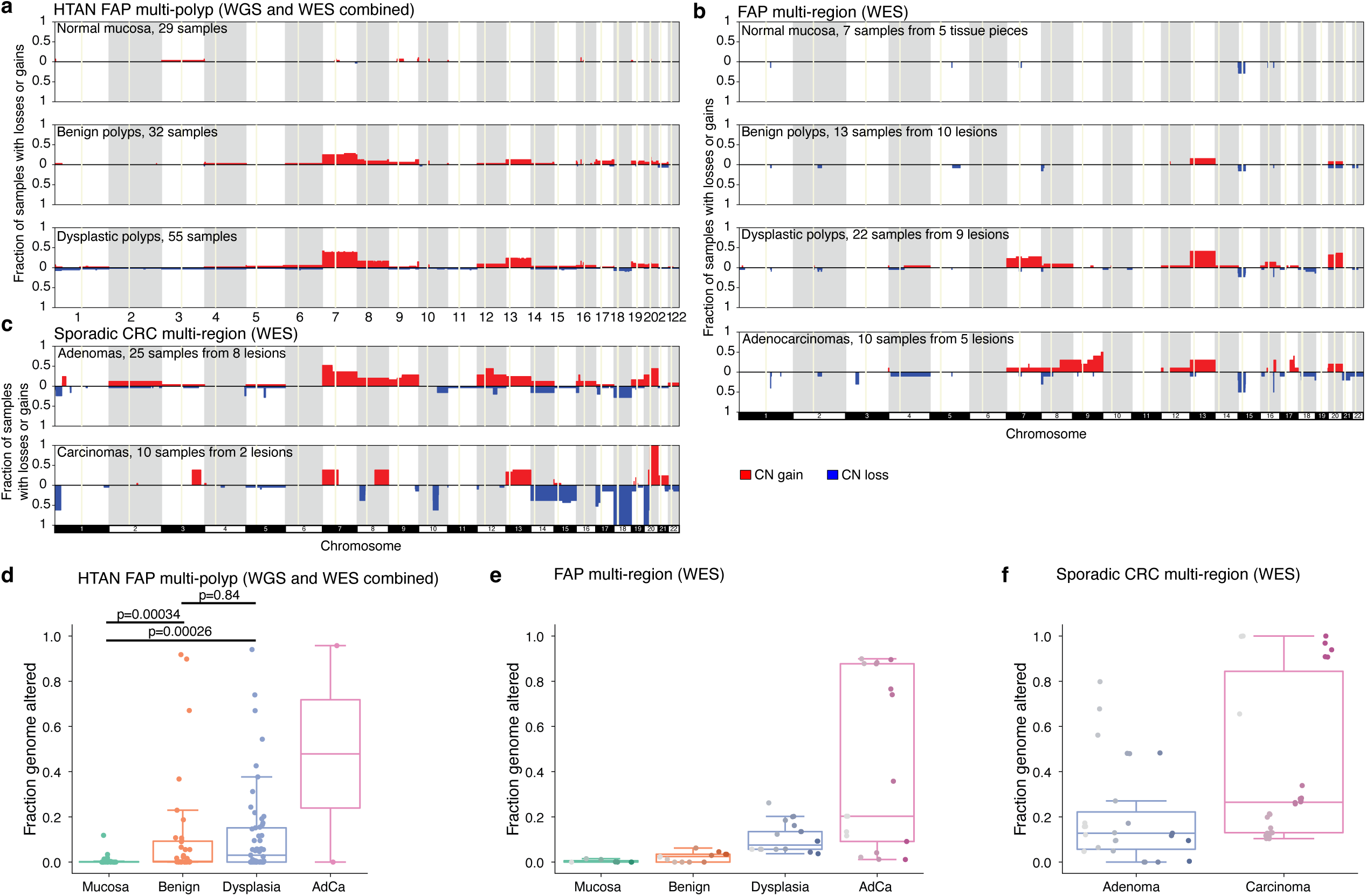
Landscape of somatic copy-number alterations in FAP lesions and sporadic colorectal tumors. **a-c.** Fraction of samples with copy-number gains (red, positive y-axis) or losses (blue, negative y-axis) in the HTAN FAP multi-polyp cohort (**a**), FAP multi-region cohort^25^ (**b**), and sporadic CRC multi-region cohort^26^ (**c**). Centromeres are shown as light yellow vertical regions. Only samples without whole-genome doubling (WGD) inferred by FACETS are shown, and the HTAN AdCas were not shown since only one sample was in this group. **d-f.** Comparison of the fraction genome altered (FGA) across disease stages for the HTAN FAP cohort (**d**), multi-region FAP cohort (**e**), and the sporadic CRC cohort (**f**). In **e-f**, the points of the same color and at the same x-axis position within each disease stage indicate multiple regions sequenced from the same lesion. WGD samples were included in these plots. In total there were n = 29 mucosal, n = 35 benign, and n = 57 dysplastic samples in the multi-polyp HTAN FAP cohort (**d**), n = 7 mucosal, n = 13 benign, n = 22 dysplastic, and n = 17 AdCa samples in the multi-region FAP cohort (**e**), and n = 27 adenoma and n = 30 carcinoma samples in the sporadic CRC cohort (**f**). P-values were computed using two-sided Wilcoxon tests. In box-and-whisker plots, the box represents the IQR with the center line representing the median, and the whiskers are the largest and smallest data values within 1.5 times the IQR from the box edges.

**Extended Data Figure 4.**
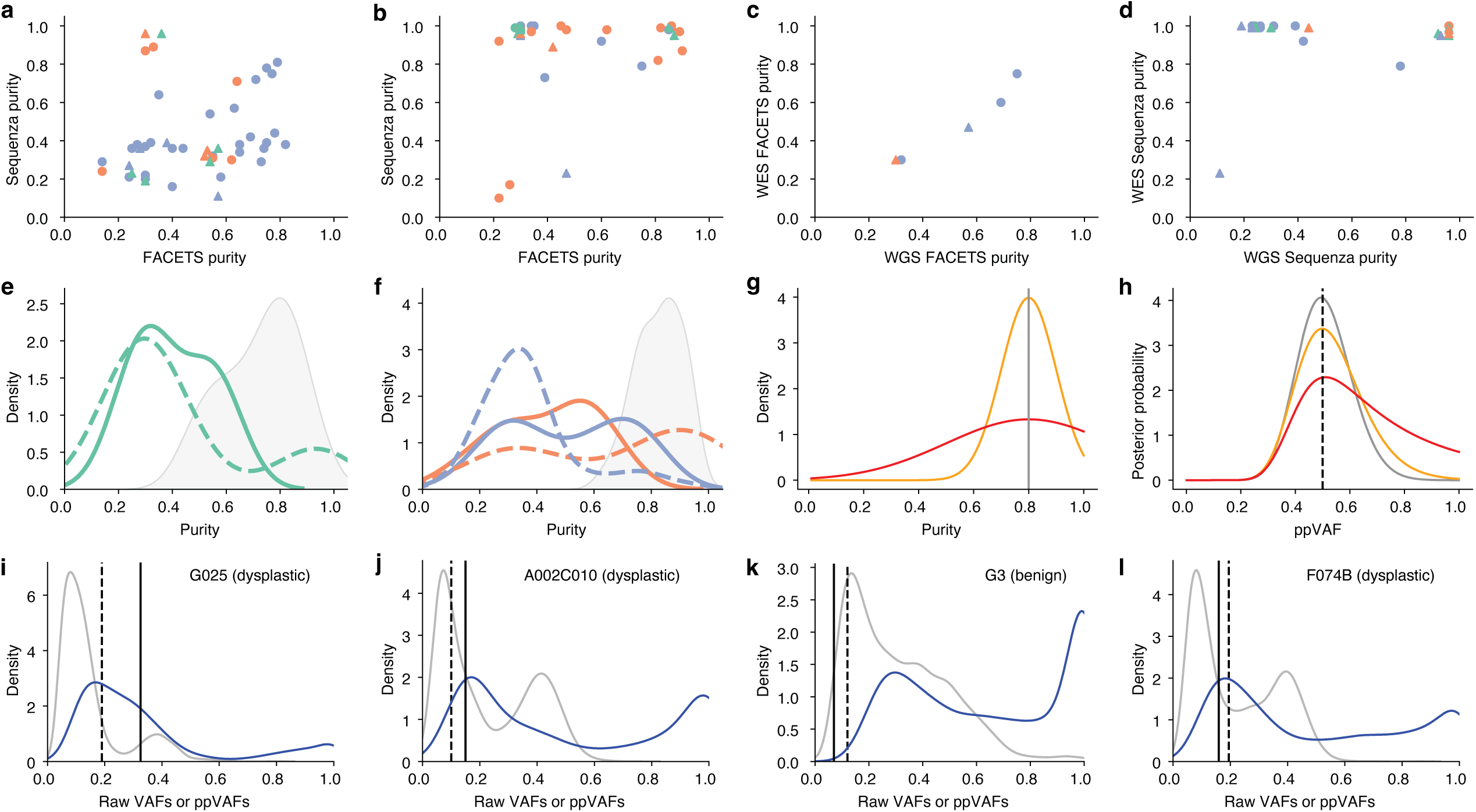
Purity estimation and ppVAF transformation in normal and premalignant samples. **a-b.** Comparison between sample purities estimated by two copy-number based algorithms (FACETS and Sequenza) in the HTAN WGS (**a**) and WES (**b**) samples. Colors indicate the type of sample (green: mucosa, orange: benign polyp, blue: dysplastic polyp) and shape indicates the inferred clonal origin of the sample from the bulk sequencing data (circle: monoclonal, triangle: polyclonal). **c-d.** Comparison between WES and WGS sample purities estimated by FACETS (**c**) or Sequenza (**d**) for samples profiled with both modalities. As before, colors indicate the type of sample (green: mucosa, orange: benign polyp, blue: dysplastic polyp) and shape indicates the inferred clonal origin of the sample from the bulk sequencing data (circle: monoclonal, triangle: polyclonal). **e-f.** The distributions of purities inferred by Sequenza (dashed colored lines; mucosa shown in green in **e**, benign (orange) and dysplastic (blue) polyps are shown in **f**) or FACETS (solid colored lines) are very different from the epithelial cell fractions measured using scATAC-seq (grey filled distributions). **g-h.** Toy examples showing estimation of ppVAFs using uncertain sample purity values, for a mutation with 20 mutant and 80 wild-type reads. Using a known sample purity with no uncertainty (vertical line in sample purity distributions in **g**), the posterior distribution for the ppVAF has the narrowest possible width (corresponding posterior distribution in **h**). As the sample purity distribution gets wider (orange and red distributions in **g**), the ppVAF posterior distributions for a mutation with the same reference and alternate allele sequencing counts get wider as well (corresponding orange and red distributions in **h**). The true ppVAF of the mutation given an 80% pure sample is noted by the dashed vertical line in **g**. **i-l.** Raw VAF distributions (grey) and corresponding ppVAF distributions (blue), computed using the scATAC-seq measured polyp sample purity distribution (shown in **f**) from four example samples where computational purity estimation using copy-number based algorithms (FACETS and Sequenza) produced poor results. In these examples, many mutations, including the bulk of the clonal mutation peak, have substantially higher VAFs than the expected clonal heterozygous VAF calculated from the FACETS purity (vertical solid line) and/or the Sequenza purity (vertical dashed line).

**Extended Data Figure 5.**
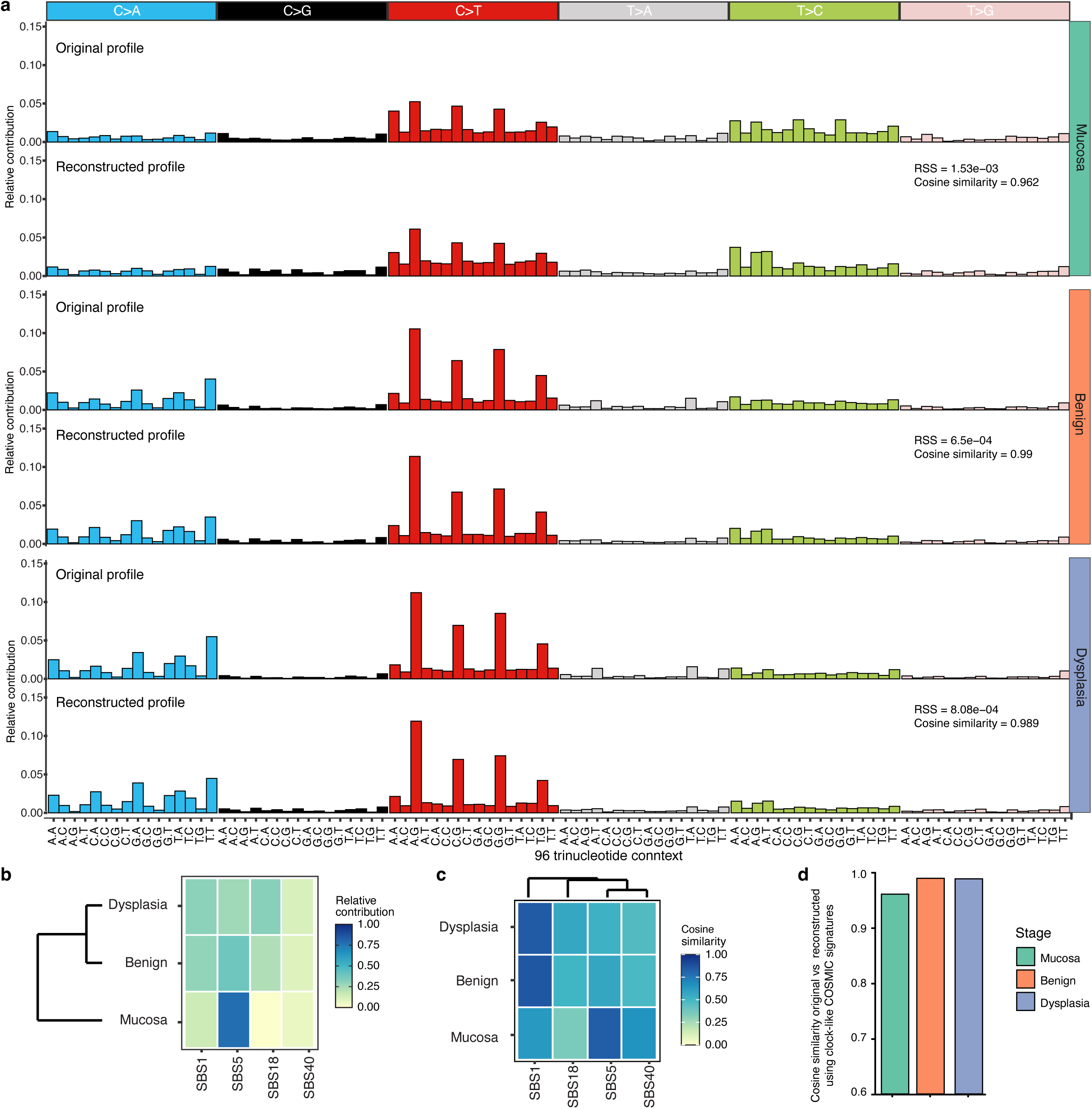
SNV signatures using WGS data and known COSMIC signatures. **a**. Original and reconstructed mutation profiles using COSMIC (v.3.2) signatures (SBS1,5,18 and 40) for mutations in normal mucosa, benign, and dysplastic polyps based on WGS. Each pair of profiles shows the original (top) and reconstructed (bottom) mutation profiles categorized by nucleotide substitution type (C>A, C>G, C>T, T>A, T>C, T>G) across 96 trinucleotide contexts. Cosine similarity and residual sum of squares (RSS) between original and reconstructed profiles are shown for each stage. **b**. Relative contributions of SBS1, SBS5, SBS18, and SBS40 signatures in each stage. The color gradient indicates the level of contribution, with darker shades representing higher contributions. **c**. Cosine similarity of signature contributions across the three stages. Darker shades represent higher cosine similarity between the stage and signature. **d.** Cosine similarity between the original and reconstructed mutation profiles using the combined signatures including the clock-like signatures (SBS1, SBS5) for normal mucosa, benign, and dysplastic polyps.

**Extended Data Figure 6.**
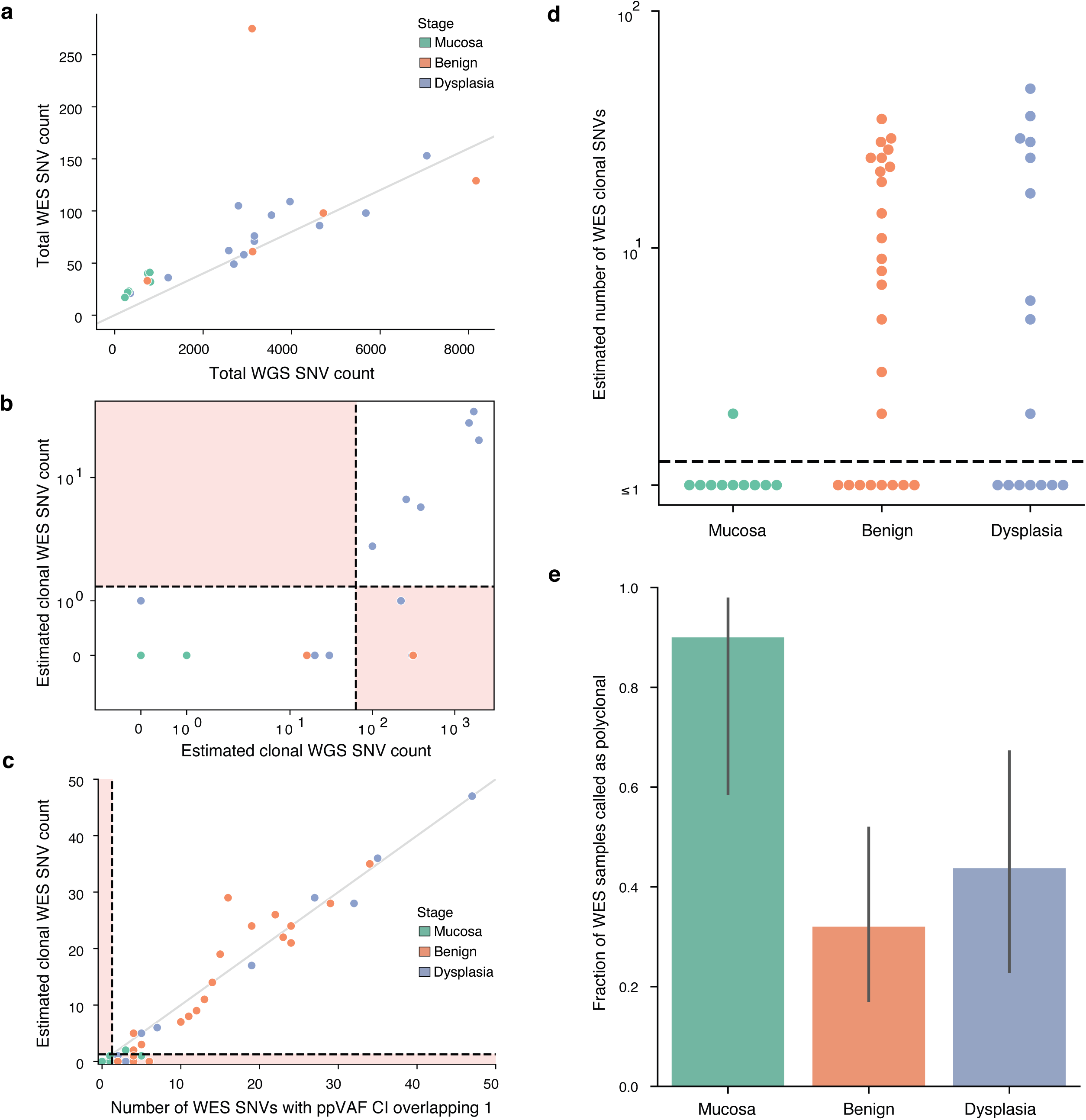
Detecting polyclonality using clonal SNV counts in WES FAP samples. **a.** Comparison of the number of total SNVs detected with WES and WGS in samples with data from both modalities. The diagonal line indicates the approximate expected relationship between WES and WGS (approximately 2% of the human genome is in the exome). **b.** Comparison between early/late MRCA classifications using WES and WGS data for samples with both modalities. Classification thresholds shown as dashed lines and regions in light red indicate samples that were classified differently using the two modalities. **c.** Comparison between counting clonal SNVs using our expected clonal counting procedure (**Methods**) and counting mutations with ppVAF upper bounds (defined as the upper ppVAF value where the posterior probability is half the maximum) equal to 1, as was used in^38^. Classification thresholds are shown as dashed lines and regions in light red indicate samples that were classified differently using the two procedures. The diagonal line shows the expected relationship if the counting procedures were equivalent (line is at y=x). In panels **a-c**, green points indicate mucosal samples, orange points indicate benign polyps, and blue points indicate dysplastic polyps (coloring as in **d-e**). **d.** Number of estimated clonal SNVs detected in WES data from FAP polyps. Samples with fewer than 0.63 clonal SNVs (dashed line, corresponding to an MRCA at < 1 year old) were classified as having an early MRCA and are likely polyclonal. **e.** Fraction of WES samples classified as early MRCA (putative polyclonal) based on estimated clonal SNV count, calculated using n = 10 mucosal, n = 25 benign, and n = 16 dysplastic samples. Error bars are 95% Bayesian credible intervals (**Methods**).

**Extended Data Figure 7.**
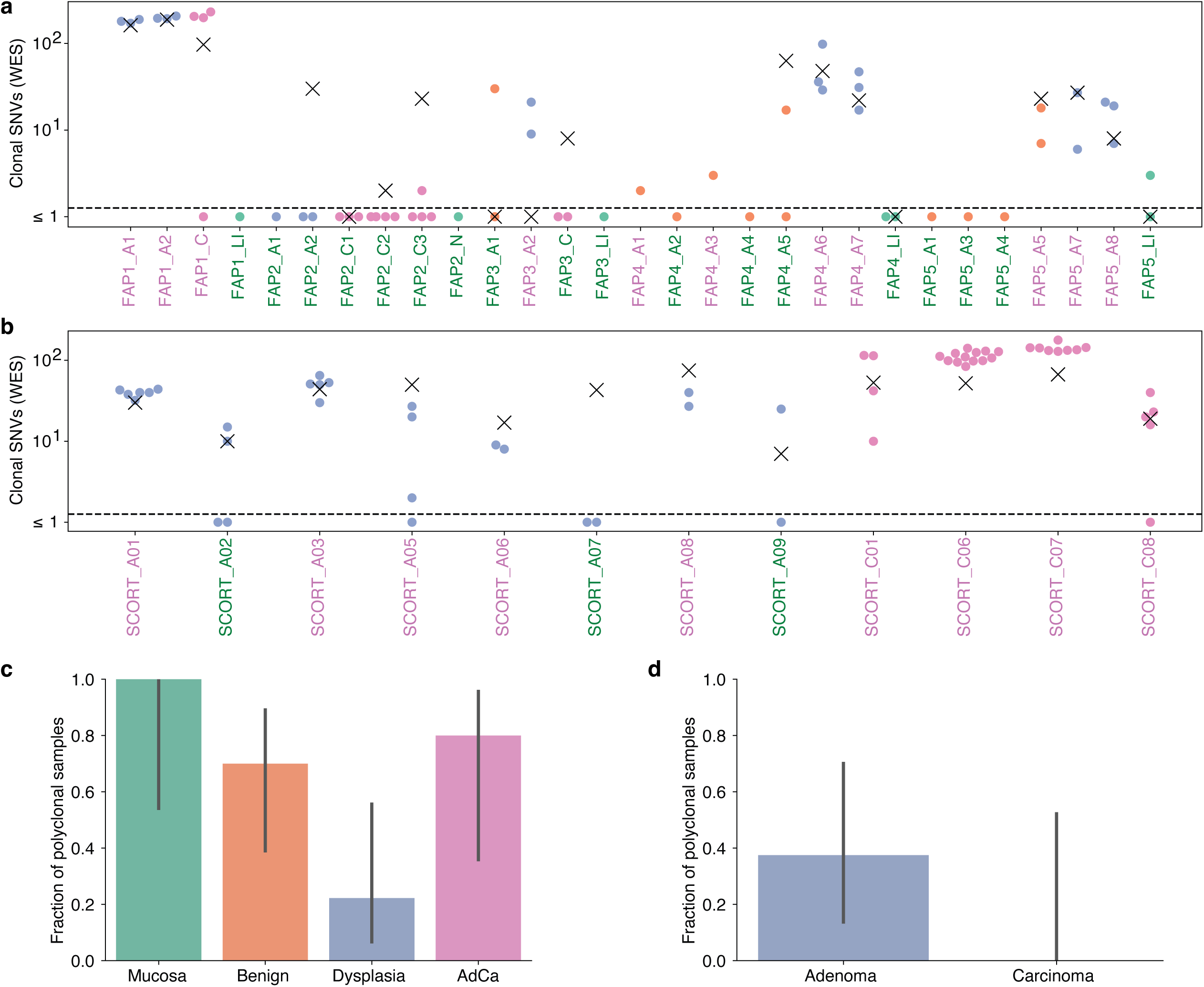
Clonal SNV counts and monoclonal/polyclonal classifications in published multi-region WES FAP and sporadic CRC cohorts. **a.** Estimated clonal SNV counts in an additional published FAP cohort with multi-region WES data^24^. Each lesion (polyp or adenocarcinoma tumor) is at an x-axis location, and sequencing data from each region in the lesion are shown as individual points. Point colors indicate sample type; green: mucosa, orange: benign, blue: dysplastic, and pink: AdCa. X’s indicate the number of SNVs shared between all regions of each lesion, and the dashed horizontal line is the polyclonal cutoff. Samples marked in green text on the x-axis were classified as polyclonal, while samples marked in purple text are monoclonal. **b.** Estimated clonal SNV counts in a published sporadic CRC cohort with WES data from adenomas and adenocarcinomas^26^. Each lesion (adenoma or adenocarcinoma tumor) is at an x-axis location, and sequencing data from each region in the lesion are shown as individual points. Point colors indicate sample type; blue is adenomas and pink is adenocarcinomas. X’s indicate the number of SNVs that are shared between all regions of each lesion, regardless of VAF, and the dashed horizontal line is the polyclonal cutoff. Samples marked in green text on the x-axis were classified as polyclonal, while samples marked in purple text are monoclonal. **c-d.** Fraction of polyclonal samples (where each sample is a lesion or normal mucosal region, not an individual sequenced region within a lesion/region) in the multi-region datasets (FAP cohort data in **c** with n = 5 mucosal, n = 10 benign, n = 9 dysplastic, and n = 5 AdCa samples, and sporadic CRC cohort data in **d** with n = 8 adenomas and n = 4 carcinomas). Colors indicate sample type as in **a** and **b**. Error bars are 95% Bayesian credible intervals (**Methods**).

**Extended Data Figure 8.**
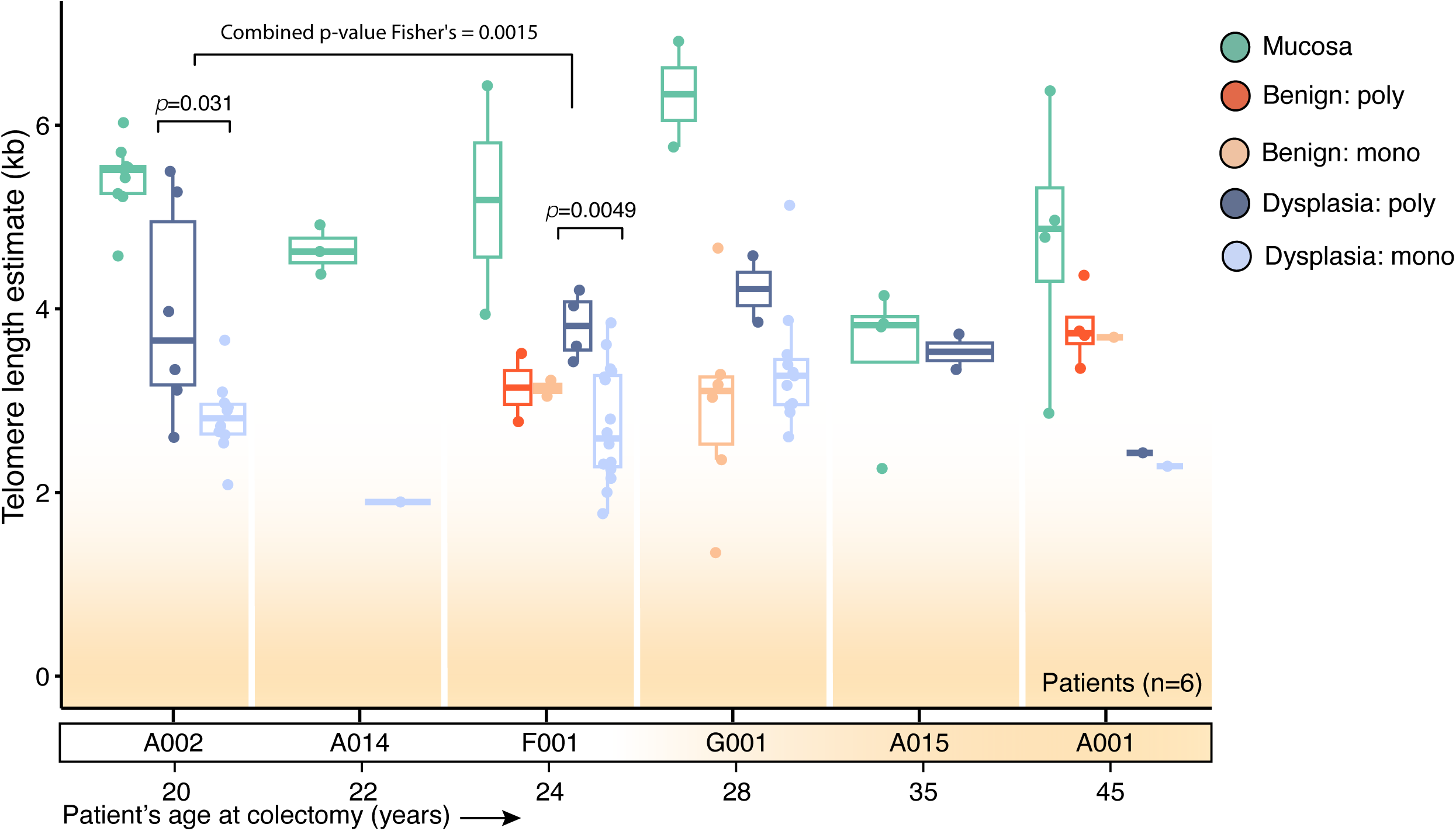
Telomere length in monoclonal and polyclonal polyps. Boxplot showing computationally estimated telomere length (TL) (**Methods**) from WGS data for normal mucosa and benign and dysplastic polyps, separated by patient. Polyclonal and monoclonal lesions are plotted separately for benign and dysplastic samples. The age at which colectomy was performed for each patient is noted under the patient IDs and patients are ordered by age. Each point is one sequenced sample from one polyp or normal mucosal region, with n = 9 mucosal, n = 6 polyclonal dysplastic, and n = 10 monoclonal dysplastic samples for A002, n = 4 mucosal and n = 1 monoclonal dysplastic samples for A014, n = 2 mucosal, n = 2 polyclonal benign, n = 2 monoclonal benign, n = 4 polyclonal dysplastic and n = 16 monoclonal dysplastic samples for F001, n = 2 mucosal, n = 6 monoclonal benign, n = 2 polyclonal dysplastic, and n = 11 monoclonal dysplastic samples for G001, n = 4 mucosal and n = 2 polyclonal dysplastic samples from A015, and n = 4 mucosal, n = 4 polyclonal benign, n = 1 monoclonal benign, n = 1 polyclonal dysplastic, and n = 1 monoclonal dysplastic samples from A001. P-values were computed to compare polyclonal and monoclonal TL in dysplastic polyps with at least two data points within individual patients using two-sided Wilcoxon rank-sum tests. The combined p-value was computed using Fisher’s method. In box-and-whisker plots, the box represents the IQR with the center line representing the median, and the whiskers are the largest and smallest data values within 1.5 times the IQR from the box edges.

**Extended Data Figure 9.**
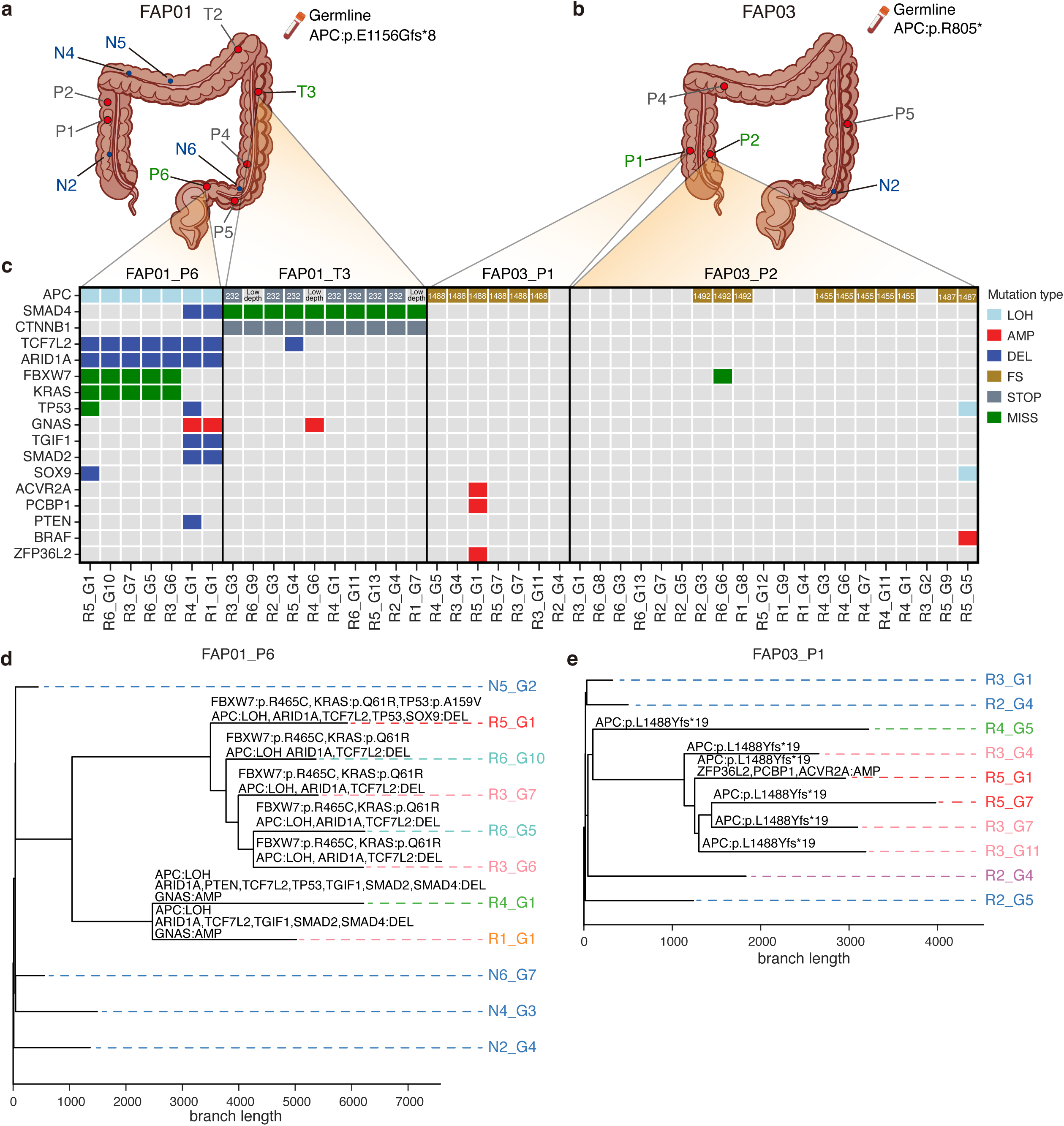
FAP sgWGS sampling distribution and additional phylogenetic trees. **a-b.** Distribution of sampled lesions for patient FAP01 (**a**) and FAP03 (**b**). Samples taken from the normal mucosa are shown in blue. Lesions that passed our filters are highlighted in green. Each patient’s germline APC mutation is annotated as well. **c.** Oncoplot heatmap of mutations in a list of 20 known CRC drivers across the four sgWGS-sampled lesions that passed our filters. For lesion FAP01_T3, this includes mutations originally filtered out (Figure 4f**; Supplementary Note 3, Section 5**). CNVs are grouped into loss-of-heterogeneity (LOH), amplification (AMP) and deletion (DEL) variants. SNVs are grouped into frame-shift (FS), premature stop-codon (STOP) and miss-sense (MISS) variants. For APC SNVs, the amino-acid position affected by the mutation is indicated as well. **d-e.** Phylogenetic trees from lesions FAP01_P6 (**d**) and FAP03_P1 (**e**). Trees were reconstructed from all SNVs and 1-base INDELs at VAF ≥ 0.2, using the neighbor-joining algorithm. Colors indicate the spatial region of the polyp or lesion from which glands were extracted, or glands from normal mucosa (in blue). Terminal branches are annotated with missense mutations, nonsense mutations, frameshift mutations and CNVs in a list of 20 known CRC drivers, if the sample below that branch carries such a mutation. R, region; P, polyp; N, normal. Graphics in **a** and **b** are adapted from BioRender, bioicons (CC BY 4.0) and/or Servier Medical Art (CC BY 4.0).

**Extended Data Figure 10.**
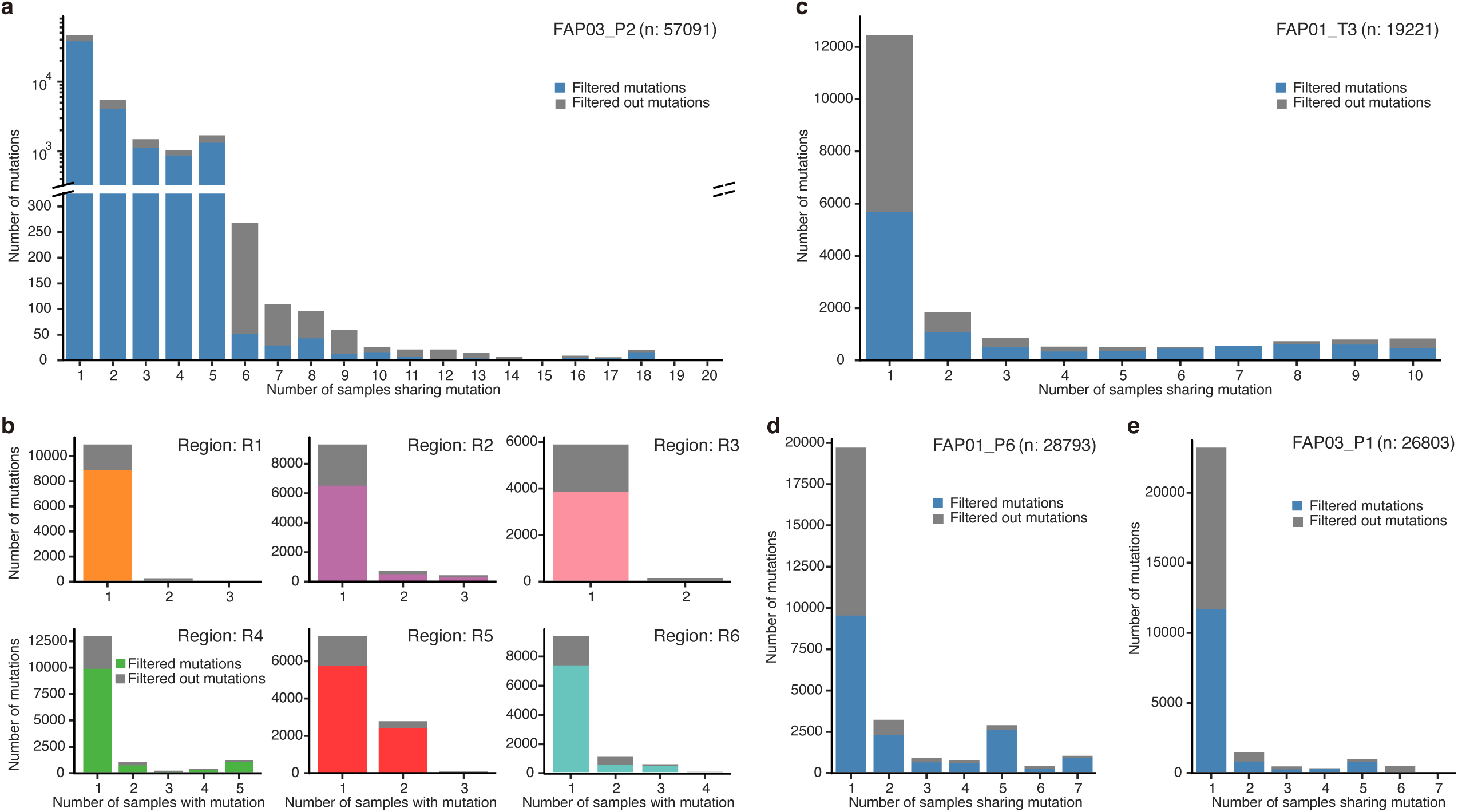
Mutation sharing across sgWGS samples within lesions. **a.** Mutation sharing across the 20 samples from FAP03_P2. The y-axis was split to be able to faithfully represent the extent of higher-level mutation sharing, while still capturing (on a log10 scale) the mutations shared by 5 or fewer samples. **b.** Mutation sharing within the six regions sampled from lesion FAP03_P2. **c.** Mutation sharing across the 10 samples from FAP01_T3. **d.** Mutation sharing across the 7 samples from FAP01_P6. **d.** Mutation sharing across the 7 samples from FAP03_P1. **a-e.** In all plots, the colored bars indicate the counts of mutations that passed our coverage and VAF filters, while the grey bar indicates the counts of mutations that got filtered out (**Supplementary Note 3**).

