## Supplementary Information for "Polyclonal origins of human premalignant colorectal lesions"

1 Department of Genetics, Stanford School of Medicine, Stanford, CA 94305
2 Stanford Cancer Institute, Stanford School of Medicine, Stanford, CA 94305
3 Key Laboratory of Quantitative Synthetic Biology, Shenzhen Institute of Synthetic Biology, Shenzhen Institute of Advanced Technology, Chinese Academy of Sciences, Shenzhen, China
4 Key Laboratory of Human Microbiome and Chronic Diseases, Sun Yat-sen University, Ministry of Education, Guangzhou, China
5 Department of Medicine, Stanford School of Medicine, Stanford, CA 94305
6 Animal Reference Pathology, Salt Lake City, UT 84107
7 Department of Pathology, Stanford School of Medicine, Stanford, CA 94305
8 Department of Computer Science, Stanford University, Stanford, CA 94305
9 Department of Biomedical Data Science, Stanford University, Stanford, CA 94305
10 Chan Zuckerberg Biohub, San Francisco, CA, 94158
11 Present address: Natera Inc, San Carlos, CA 94070
12 Present address: Department of Computational Biology, Mohamed bin Zayed University of Artificial Intelligence (MBZUAI), Abu Dhabi, UAE
*These authors contributed equally to this work
#These authors jointly supervised this work and are co-senior authors

Supplementary Figure 16. VAF distributions for WES of dysplastic polyclonal polyps. ... 22
Supplementary Figure 17. VAF distributions for WES of dysplastic monoclonal polyps.. 22
Supplementary Note 1: Estimating the number of clonal mutations from bulk sequencing data 23

### Methods

##### *Sample collection*

Samples were procured from colons of consented FAP patients (IRB-47044; PreCancer Atlas for Familial Adenomatous Polyposis) during total colectomy procedures at Stanford Hospital. Patient characteristics are reported in **Supplementary Table S1**. Patient colons were directly transported from the surgical suite to the pathology gross room at the time of resection. Colons were then rinsed with water and flayed open using a scalpel lengthwise. Polyp-adjacent normal mucosa, polyps, and adenocarcinoma tissues were carefully removed from the colon and stored in cryovial tubes and flash frozen in liquid nitrogen. For patients A001, A002, A014, and A015, exact locations of sampling were recorded by replacing the resected tissue with numbered markers; once sample collection was completed, photographs were taken to determine the spatial coordinates of the tissue samples in pixel units, the diameter of the markers were used as a scaling factor to convert pixel units to centimeters. Samples were classified as normal mucosa, benign, dysplasia, or adenocarcinomas (AdCa) by expert pathologists.

##### *DNA isolation, library preparation, and sequencing*

Flash frozen tissue sample DNA was isolated by pulverizing tissue with mortar and pestle. Pulverized tissue was prepped for DNA extraction using the Qiagen All-prep kit (cat. 80204). Buffy coat DNA was isolated by placing 200 ul directly into 600 µl Buffer RLT with β-ME at 1:100 dilution following the manufacturers guidelines. DNA libraries were created for samples whose DNA yield was > 300 ng. WGS and WES libraries were prepared using either NEBNext DNA Library Prep Kit (cat. no. E7645S) or TruSeq Nano DNA Kit (cat. No. 20015964) following the manufacturers' guidelines. The SureSelect Human All Exon v6 exome capture kit was used for samples subject to WES. WGS and WES libraries were sequenced on the Illumina NovaSeq 6000 to a depth of 60x and 90x, respectively. Some fresh frozen samples were profiled with both WGS and WES, as indicated (**Supplementary Fig. 1**; **Supplementary Tables 2, 3**), when both were available WGS was used for analysis. Given the limited number of AdCa samples (n=2) and the focus on pre-malignancy, comparisons of SNVs within the FAP multi-polyp cohort were restricted to mucosa, benign and dysplastic polyps.

##### *Sequence mapping to the reference genome*

After sequencing, using an in-house pipeline for mapping[18,45], we mapped FASTQ files with bwa mem (v1.10, bwa mem -r 1.2 -t 40 -R) for hg38. We created a target interval list file with GATK's RealignerTargetCreator (v3.4-46, --allow_potentially_misencoded_quality_scores) and IndelRealigner (v3.4-46, -compress 5) with Mills_and_1000G_gold_standard.indels and dbSNP138 indels as references. To reduce technical errors by the sequencer, called bases were recalibrated with BaseRecalibrator (v3.4-46) against the previous indel references and the SNPs from dbSNP138. PrintReads (v3.4-46, -DIQ --emit_original_quals -BQSR *BaseRecalibrator-output*) outputs well-formed reads with the bases recalibrated. Finally, to create the final .bam file, duplicate reads are marked with MarkDuplicates (Picard v2.18.7, REMOVE_DUPLICATES = false, VALIDATION_STRINGENCY = LENIENT, ASSUME_SORTED = true).

For the two external WES cohorts, described below, raw fastq files were mapped using the Nextflow-based pipeline nfcore/sarek (v2.7.1)[46] with the reference genome GRCh38/hg38. The rest of the downstream analysis was performed uniformly for all the three cohorts.

##### *Published multi-region sporadic CRC and FAP cohort analysis*

Comparative analysis was performed on two published WES multi-region cohorts, including five FAP patients and 12 sporadic CRC patients (8 adenomas and 4 adenocarcinomas). Raw sequencing data were downloaded from the Genome Sequence Archive (HDAC000073) for the FAP cohort (Li *et al.*[24]) with average coverage of ~53x and from the European Genome Phenome Archive (EGAS00001003066) for the sporadic CRC cohort (Cross *et al.*[25]) with average coverage ~53x (**Supplementary Fig. 1**). For the multi-region sporadic cohort we excluded patients reported to have high level microsatellite instability, in the original study. Since no patient sex information was available for the sporadic CRC cohort, we manually inspected the WES coverage at the *SRY* locus of the normal reference samples to determine the sex of each patient in this cohort. All downstream analysis was performed using the same approaches as detailed for the samples collected in this study.

In the multi-region FAP cohort, samples were classified adenomas as either stage Normal, I, II, III, or AdCa, or assigned pathology statuses of LGIN (Low-Grade Intraepithelial Neoplasia), HGIN (High-Grade Intraepithelial Neoplasia), MDAC (Moderately Differentiated Adenocarcinoma). In order to ensure consistency and comparability with the HTAN FAP cohort classification rubric, which includes 1) mucosa, 2) benign, 3) dysplasia (low-grade, high-grade), and 4) adenocarcinoma, we applied the following criteria: All samples with stage normal were classified as mucosa. Polyps with stage I and pathology LGIN were classified as benign. Only two polyps (FAP5_A5_R1, FAP5_A5_R2) with stage II and pathology LGIN, but with a polyp size of 5mm or less, were classified as benign. Polyps with stage II or III and pathology LGIN or HGIN were classified as dysplasia. Polyps with stage AdCa and pathology MDAC were classified as adenocarcinoma (AdCa).

##### *Somatic variant calling, filtering, and annotation*

Somatic variants were called on fresh frozen samples by taking consensus calls from two different methods against a patient-specific blood normal reference. First, we used nf-core/sarek (v2.7.1)[46] Mutect2[47] following the GATK (v4.1.7.0)[48] best practices guideline with default options using the GRCh38 reference and gnomAD (r2.1.1) germline resource. Raw somatic variants were filtered first using GATK's *FilterMutectCalls* using default options and a contamination table produced by *CalculateContamination*. Second, we separately called somatic mutations on each sample using Strelka2 (v2.9.10)[49], using input candidate indels called with Manta (v1.6.0)[50]. Only variants called by both Mutect2 and Strelka2 were retained and further filtered by a custom python script, *'VariantFilter'* without filtering clustered mutation events (https://github.com/rschenck/VariantFilter/tree/strelka). For a variant to be retained by this script we required a minimum of 2 variant reads in the sample, a minimum variant allele frequency of ≥0.01, and minimum coverage of 10 reads at the locus for both the sample and the normal reference. Variants reporting any number of variant reads from Mutect2 within the normal sample were removed. Upon inspection of the aligned reads in *APC*, we found that one sample (G005) had a complex frameshift deletion (**Supplementary Fig. 7**) that was denoted differently by Strelka2 and Mutect2 and thus was not retained by VariantFilter. We manually added the relevant calls back to the consensus variant set.

Consensus variants were further filtered to remove mapping errors and other technical artifacts. Variants on the Y chromosome were filtered out in samples from female individuals. We removed variants in telomeres and other problematic genomic regions (e.g., low mappability regions) identified by ENCODE, as well as variants within $10^5$ bp of centromeres. In our in-house HTAN bulk WES and WGS datasets, we also removed variants present in more than one sample. While some of these shared variants may represent true somatic mutations that occurred early in development or spread through the colon as part of a high-fitness epithelial clonal expansion, the patterns of sharing suggest the majority are likely technical artifacts. However, to retain high-frequency driver mutations that may occur independently in multiple lesions, we retained mutations found in previously-identified colorectal cancer driver genes[51], even if they were found in multiple samples.

We used vcf2maf (v1.6.19; https://github.com/mskcc/vcf2maf) to convert VCF files to MAF and annotated using VEP (v103.1) from Ensembl[52] with --ncbi-build argument set to GRCh38.

Since the cohorts in this study include samples with WGS and WES, we compared mutation burden between samples and disease stages by counting non-silent exonic coding mutations. For samples with both WGS and WES we used WGS data for these calculations.

##### *Copy number analysis*

We used FACETS-SUITE (v2.0.8; https://github.com/mskcc/facets-suite) with allele-specific CNA caller FACETS (v0.6.1)[53] to identify CNAs and whole-genome doubling (WGD) events using WGS and WES data. We performed parameter tuning with six combinations of parameters for WGS and WES: purity_cval (WGS: 2000,1000; WES: 1000, 500), cval (WGS: 1000, 500; WES: 500, 250) with normal_depth set to 25, 30, and 35. The optimal parameter selection for each sample was determined by an aggregated score based on six possible extremes: extreme diplogR defined as abs(dipLogR) > 0.8, number of whole chromosome losses, number of whole chromosome loss of heterozygosity, extreme fraction of homozygous deletions defined as > 0.8, extreme loss of heterozygosity defined as > 0.5 of the genome, extreme ploidy defined as >6 or <1.8. We selected the parameter choice that minimized these extreme cases, as detailed elsewhere[54].

The FACETS expectation-maximization total and minor copy number calls were then further filtered to exclude problematic genomic regions and other artifacts. We excluded all segments with >=30% overlap with centromeres, telomeres, or other ENCODE blacklisted areas of the genome or that were shorter than 1 Mbp. Low-frequency subclonal events were excluded by filtering out all segments with FACETS estimated cell fraction < 0.15. For gene and mutation copy number identification, the remaining segments were extended to cover the entire genome between the telomeres of each chromosome. For each gap in the filtered copy number segments, the flanking segment with copy number closest to normal was extended to span the gap. All mutations on chromosome Y were assumed to have a total copy number of 1 in males and 0 in females. In the oncoplots, genes with total copy number greater than normal were reported to have a copy number gain, genes with total copy of 0 were reported to have a deletion, and genes with a minor copy number of 0 and total copy number greater than 1 were reported to have loss of heterozygosity (LOH). Copy number alterations were not plotted for samples inferred to have WGD.

The fraction of genome altered (FGA) was computed as in FACETS-SUITE, by summing the lengths of the segments (extended to cover the whole genome, as explained above) with abnormal total or minor copy number and normalizing to total genome length between telomeres. Only autosomes were used for this calculation.

For samples with both WGS and WES data, we exclusively used the WGS copy number calls, including for correcting VAF values for copy number as described below.

##### *Purity estimation*

In order to estimate the fraction of epithelial cells that harbor each somatic mutation (the ppVAF), we first estimated the fraction of sampled cells that were epithelial, which we term the sample purity in this study. We initially attempted to use individual allele-specific copy number callers (FACETS and Sequenza (v3.0.0)[55]) designed for tumor sequencing data to estimate purity. However, we found that none of these computational methods were satisfactory for assigning purity to our samples, likely due to the low number of CNAs found in our premalignant samples relative to the tumor samples these algorithms were designed to process. In particular, these algorithms were not able to resolve the purity of the normal mucosal samples, which have very few CNAs. Therefore, many polyp samples had raw VAF distributions that were unlikely given the computationally inferred purity values from the genomic data, with several mutations at VAF values substantially higher than would be expected for clonal heterozygous mutations at the inferred purities (**Extended Data Fig. 4**).

More generally, the design principle behind all of these algorithms that estimate purity from bulk genomic data alone is not ideal for polyclonal, nonmalignant tissues. This is because these purity callers do not directly estimate the fraction of cells of a desired phenotype (here, epithelial cells), but rather estimate the fraction of cells in the sample that belong to the highest-frequency subclone defined by CNAs and/or somatic mutations. In polyclonal samples, the frequency of the major subclone is often not representative of the overall frequency of the desired cell type in the sample, leading to underestimation of sample purity.

To address these issues, we instead used single-cell ATAC sequencing (scATAC-seq) data from the same patient cohort[28] to estimate the distribution of epithelial cell fractions in normal, premalignant, and malignant samples using the published cell type assignments. The scATAC-seq measured distributions of epithelial cell fractions were very different from the computationally-inferred purity value distributions (**Extended Data Fig. 4**), highlighting issues with estimating purity with these algorithms on largely diploid samples. We therefore used these orthogonally-measured distributions to estimate ppVAF values, both in our patient cohort and in the FAP and sporadic multiregion cohorts we reanalyzed, which had similar issues with computational purity estimation.

##### *Calculating somatic mutation ppVAFs*

We estimated the purity and ploidy adjusted variant allele frequencies (ppVAFs) for each somatic mutation. The ppVAF is the nonmalignant/premalignant analogue to the cancer cell fraction (CCF), and describes the fraction of epithelial cells which have a mutation. We calculated the posterior probability distribution for ppVAF values using the same general approach as for CCF calculation[76] by maximizing the probability of generating the observed alternate allele read count given a ppVAF value and purity. The alternate allele count is a random variable distributed binomially,

$$t_{alt} \sim Binom(t_{depth}, p), \text{ where } p = CN_{mut}\,\rho f/(2(1-\rho) + \rho CN_{total}),$$

given the total sequencing depth at the locus $t_{depth}$, the total copy number and mutant copy number at the locus $CN_{total}$ and $CN_{mut}$ (computed by FACETS and postprocessed as described above), the ppVAF value *f*, and a purity value ρ. $CN_{mut}$ was assumed to be equal to the FACETS-estimated major allele copy number or 1 if the FACETS was not able to estimate a major allele copy number. Mutations were excluded from further analysis if the estimated total copy number was zero. Since we could not accurately estimate a purity value for each sample individually, we instead used the distribution of purity values *P*(ρ) from the scATAC-seq data as described in the previous section to compute the expected probability of each ppVAF value given the observed read counts, marginalized over the distribution of sample purities. The posterior probability of a given ppVAF value *f* can therefore be calculated as

$$P(ppVAF \;=\; f \mid t_{alt}) \;=\; \frac{\int_0^1 (t_{alt} \mid \rho, f)\; P(\rho = r) dr\, P}{\int_0^1 \int_0^1 P(t_{alt} \mid \rho, f)\; P(\rho = r) dr df}$$

assuming a uniform prior distribution over *f*, where $P(t_{alt} \mid \rho, f)$ is the probability mass function for the binomial random variable $t_{alt}$ defined above. This posterior probability was computed numerically by evaluating $P(t_{alt} \mid \rho, f)$ across a mesh and summing appropriately. The maximum a posteriori (MAP) point estimate of the ppVAF was computed by numerically maximizing this posterior probability over a 1000-element grid of ppVAF values from 0.001 to 1, and was computed and used as the ppVAF estimate for each somatic mutation detected.

##### *Identifying clonal mutations and estimating the clonal SNV count using bulk sequencing data*

The number of clonal SNVs per sample was estimated using the posterior probability ppVAF distributions $P(ppVAF \;=\; f \mid t_{alt})$ for somatic SNVs in each sample calculated as defined in the section above. Our general strategy was to classify somatic SNVs with cumulative posterior density at ppVAF above value *F* greater than some probability threshold *c* (i.e., $P(ppVAF \;>\; \varepsilon) \;>\; c$) as "clonal". However, there is no obvious optimal value for either $\varepsilon$ or $c$, since the posterior ppVAF density depends not only on the true underlying ppVAF of each mutation, but also on the total sequencing depth at the mutation locus (**Supplementary Note 1**). Therefore, we empirically optimized this procedure to enhance discrimination between clonal mutations binomially simulated with the same sequencing depth and purity distribution as in our HTAN polyp samples and subclonal mutations from our HTAN mucosal samples (**Supplementary Note 1**). In our final optimization procedure, we first selected the value of *c* that maximizes the accuracy of the clonal count estimate assuming 50% of mutations were clonal, then used the preliminary estimated clonal mutation fraction to iteratively refine our threshold *c* until the estimated number of clonal mutations remained stable across iterations (for further details, see **Supplementary Note 1**). This method of counting clonal SNVs was compared to the method used in [36], which defined the number of clonal SNVs as the number of SNVs with ppVAF upper bound = 1 (defined as the ppVAF value $f_{upper}$ where $P(ppVAF \;=\; f_{upper}) \;=\; 0.5\, P(ppVAF \;=\; f)$, where *f* is the MAP best point estimate for the SNV ppVAF, as previously described[53]).

##### *Identifying polyclonal samples using clonal SNV counts*

The number of clonal SNVs can be used to estimate the age at which the cell that founded the sample or lesion (the most recent common ancestor, or MRCA) existed. All somatic mutations present in the MRCA will be found in all crypts in the polyp and will be clonal. Assuming somatic

SNVs accumulate and fix over time as a constant Poisson process in each crypt, the number of clonal somatic SNVs in the resulting lesion is a measure of how long after somatic mutations began accruing (likely at conception) the MRCA existed. Published data quantifying SNV accumulation rates in human intestinal organoids suggest that approximately 36 SNVs accumulate per year in each intestinal stem cell[56]. This per-cell mutation rate is the same as the average rate of accumulation of clonal mutations within each crypt, since the mutations from each individual stem cell in a crypt fix and become clonal in the crypt periodically. The expected number of clonal SNVs in a crypt is therefore approximately 36 multiplied by the number of years between conception and the MRCA time, so the MRCA time in years after birth was therefore estimated as $n_{clonal}/36 - 0.75$. The polyclonal cutoff of 63 clonal SNVs for WGS data therefore corresponds to an MRCA time of 1 year after birth. The polyclonal cutoff for WES (1.26 clonal SNVs) was chosen by assuming approximately 2% of the human genome is in the exome (**Extended Data Fig. 6**). For the published multiregion WES cohorts (both FAP and sporadic), a sample/lesion was considered polyclonal if at least one of the regions had one or fewer clonal SNVs.

##### *Oncoplots*

Oncoplots were generated using the oncoplot function from the R/Bioconductor package maftools (2.17.0)[57] by using the curated list of COADREAD driver genes[51]. Genes with exonic non-silent mutations in more than one sample were included in **Fig. 1b** and **Extended Data Fig. 1a**.

##### *Mutational signature analysis*

We used the R/Bioconductor package MutationalPatterns (v3.9.1)[58] to identify known COSMIC Single Base Substitution (SBS) signatures in our cohort using all WGS somatic mutations. We generated mutational profiles across 96 contexts using the *fit_to_signatures* function to align with known COSMIC (v3.2) SBS mutational signatures including the clock-like signatures (SBS1, SBS5, SBS40) and SBS18. The cosine similarity was then calculated by comparing the original mutation matrix with the reconstructed matrix using the *cos_sim* function.

##### *Telomere length estimation and comparison*

Telomere content was estimated from WGS data from our cohort using TelSeq (v0.0.2)[59] with default parameters. We compared estimated telomere lengths from monoclonal and polyclonal dysplastic samples for each patient separately to account for differences in age and telomere shortening rate, using only patients with at least two monoclonal and at least two polyclonal dysplastic samples (A002 and F001). Each intrapatient comparison was done with a two-sided Wilcoxon rank sum test and p-values were combined using Fisher's method.

##### *Single-crypt whole genome sequencing*

We isolated individual colorectal crypts from two FAP patients (FAP01, male, 35 years old and FAP03, male, 38 years old) recruited from the Sixth Affiliated Hospital of Sun Yat-sen University (**Supplementary Table 4**) under the approved IRB protocol 2019ZSLYEC-06. The method of single-crypt isolation and WGS is based on adaptation of a previously published protocol[60]. In order to sample distinct spatial regions from each lesion, we first cut out different regions from the fresh frozen tissue. Then, the tissue pieces were placed on a clean glass slide under the microscope with 20μl PBS. Subsequently, individual crypts were manually dissociated using a 23G (0.6*25mm) injection needle and aspirated using a pipette. Each crypt was transferred to a PCR tube with 20μl protease buffer[91]. The crypts with enough extracted DNA (>1ng) were selected for WGS library construction using the Vazyme TruePrep DNA Library Prep Kit V2 (TD501/502/503, Vazyme, China) or the Hieff NGS® OnePot Pro DNA Library Prep Kit V2 (Yeasen Biotechnology, China) according to the manufacturer's instructions. Libraries were sequenced on an Illumina NovaSeq platform (Illumina, US).

SNVs from the single-crypt WGS data were called using an equivalent pipeline to our bulk WGS data analysis. In brief, we again called consensus variants across two callers, Mutect2[47] and Strelka[49], using VariantFilter (https://github.com/rschenck/VariantFilter/tree/strelka2) with default settings. Resulting VCF files were converted to MAF using vcf2maf (v1.6.19; https://github.com/mskcc/vcf2maf) and annotated using VEP (v103.1) from Ensembl[52] with --ncbi-build argument set to GRCh38.

To call CNVs from the sgWGS data, we used Sequenza v3.0.0[55]. Sequenza segment calls were filtered to be over 10kb in size, and we used depth ratio cutoff values of (2.5/2) and (1.5/2) for amplification (AMP) and deletion (DEL) respectively. Loss of heterozygosity (LOH) was called at depth ratios in between those cutoffs, and we added an additional cutoff to remove LOH calls if the B-allele frequency was above 0.2.

To generate the variant set used for phylogenetic inference, we first filtered to only include SNVs and 1-base INDELs. We then applied a set of sample-level and variant-level filters inspired by the published Sequoia pipeline[61] to select for glands consisting of a single dominant clone and to reduce false-positive variant calls. To address the latter, we filtered out variants with a VAF below 0.2 (increasing stringency over the 0.1 default of VariantFilter, see **Supplementary Note 3, Section 1**), or with a total sequencing depth below 10 or above 100 (**Supplementary Note 3, Sections 3 & 4**). To select for samples consisting of a single clone, we implemented a truncated binomial mixture model, similar to Sequoia. In brief, we fit binomial mixtures with one, two and three components, truncated at the VAF threshold noted above. We then used the Akaike information criterion (AIC) to select the best-fitting model among the three. Given the best-fitting model, we classified a sample as sufficiently clonal if its largest mixture component (as estimated by the mixture weights) was estimated to be at a VAF of 0.3 or higher (**Supplementary Note 3, Section 2**). This sample-level clonality filter removed 59/129 samples, leaving us with 70 samples passing the filter. Among the samples passing the filter, we further filtered for lesions where at least 50% of samples passed the filter, and the total number of samples passing the filter was at least 5. This further reduced the dataset to 44 samples across 4 lesions, two from each patient.

To generate phylogenetic trees from the lesions passing our filters, we first arrayed all SNVs and 1-base INDELs from the samples of that lesion in a multiple sequence alignment (MSA) matrix, where a sample carried the reference allele if a variant was absent, the alternative allele if its VAF was at least 0.3, or an ambiguous base (N) if a variant was present but at VAF below 0.3. This was done to further guard against false positive variant alleles. The MSA was then used to estimate a neighbor-joining tree, using the Biopython Phylo package v1.85 (see **Supplementary Note 3, Section 6** for a comparison with trees inferred using maximum parsimony). We annotated the phylogenetic trees with missense mutations, nonsense mutations, frameshift mutations and CNVs in a set of 20 known CRC drivers, using the previously mentioned list of COADREAD driver genes[51]. To discover SNVs in driver mutations for annotation, we first scanned for them in the filtered mutation set of all samples in a lesion. We then went through the filtered-out mutations to see if any of the putative drivers had been filtered out, for example due to falling below the detection limit in a low-coverage sample (**Supplementary Note 3, Section 5)**. If so, we added the filtered-out mutations as annotations to the tree, marking them in grey.

##### *Statistical analysis*

Bayesian credible intervals for binomial data (e.g., fraction of clonal mutations, fraction of polyclonal samples) were computed by updating a uniform prior with the observed count data. The posterior distribution of this fraction given the observed number of successes ($s$) and failures ($f$) is therefore distributed as Beta($1+s$, $1+f$). The plotted 95% credible intervals describe the uncertainty around the point estimate due to sampling.

### Supplementary figures and legends

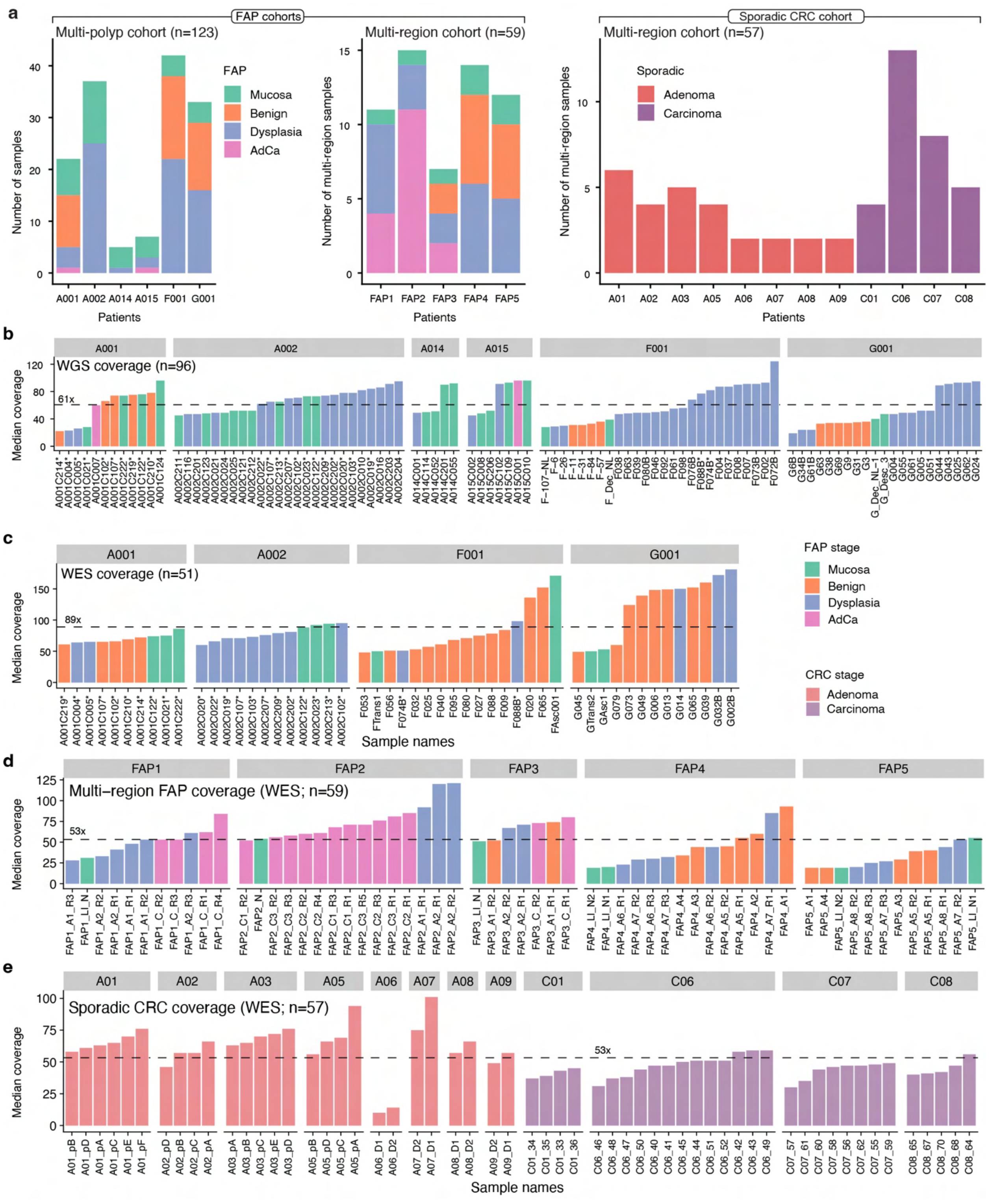

**Supplementary Figure 1. Stage distribution and sequencing coverage for samples across three cohorts.**

**a**. Stacked bar plots showing the distribution of sample stages for the two FAP (multi-polyp and multi-region) and sporadic CRC (multi-region) cohorts. **b**. Median sequence coverage for whole genome sequencing (WGS) samples (n=96; mean coverage=61x). **c**. Median sequence coverage for whole exome sequencing (WES) samples (n=51; mean coverage=89x). Samples with both WGS and WES are labeled with asterisks. **d**). Median sequence coverage for WES samples from FAP multi-region cohort (n=59; mean coverage=53x). **e**). Median sequence coverage for WES samples from sporadic CRC (multi-region) cohort (n=57; mean coverage=53x). Coverage values computed by mosdepth[62].

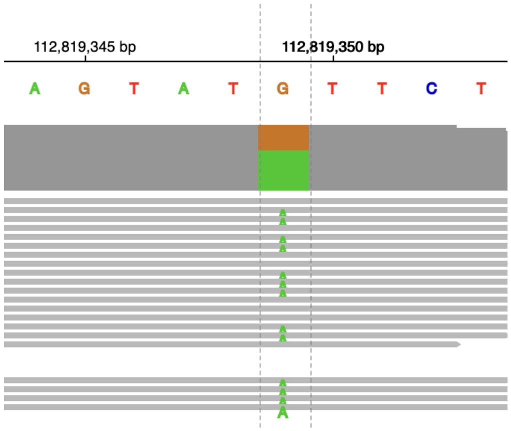

**Supplementary Figure 2. Germline APC mutation for patient A001.**
Integrative Genomics Viewer (IGV) showing supporting reads for germline *APC* mutation in patient A001.

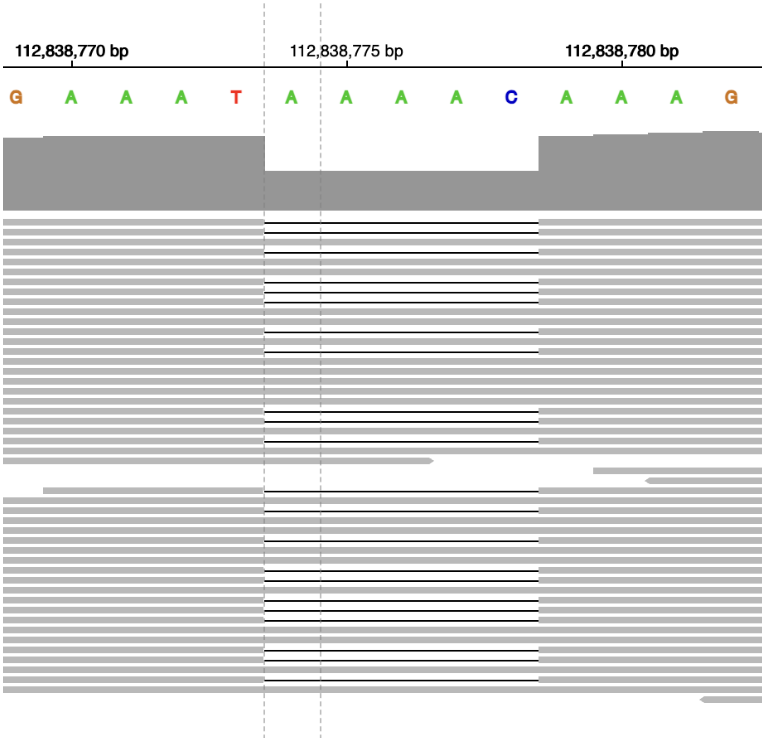

**Supplementary Figure 3. Germline APC mutation for patient A002.**
Integrative Genomics Viewer (IGV) showing supporting reads for germline *APC* mutation in patient A002.

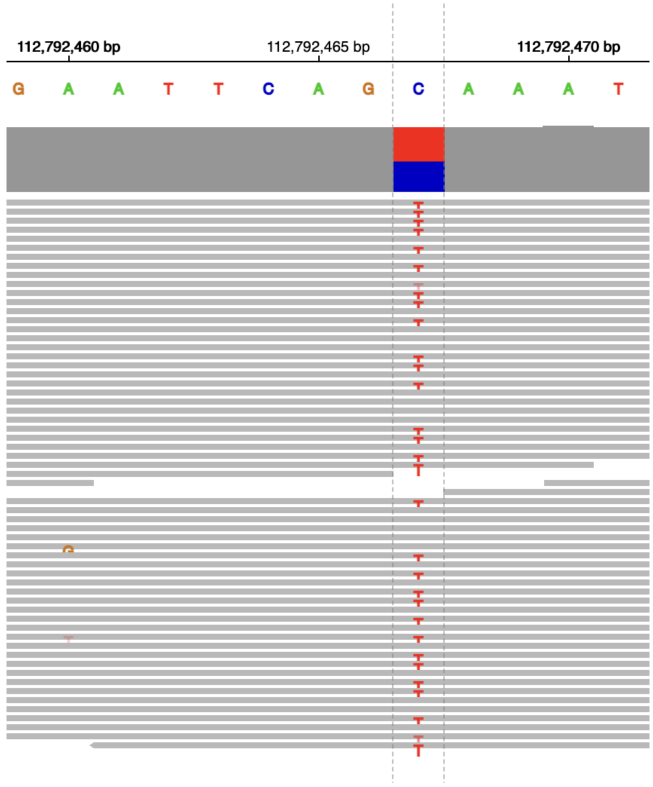

**Supplementary Figure 4. Germline APC mutation for patient A015.**
Integrative Genomics Viewer (IGV) showing supporting reads for germline *APC* mutation in patient A015.

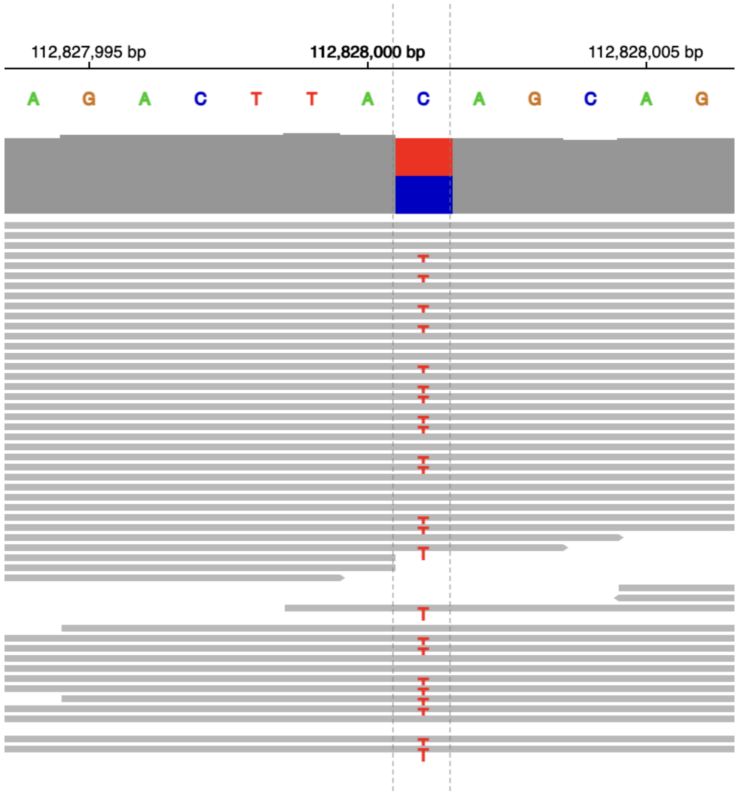

**Supplementary Figure 5. Germline APC mutation for patient F001.**
Integrative Genomics Viewer (IGV) showing supporting reads for germline *APC* mutation in patient F001.

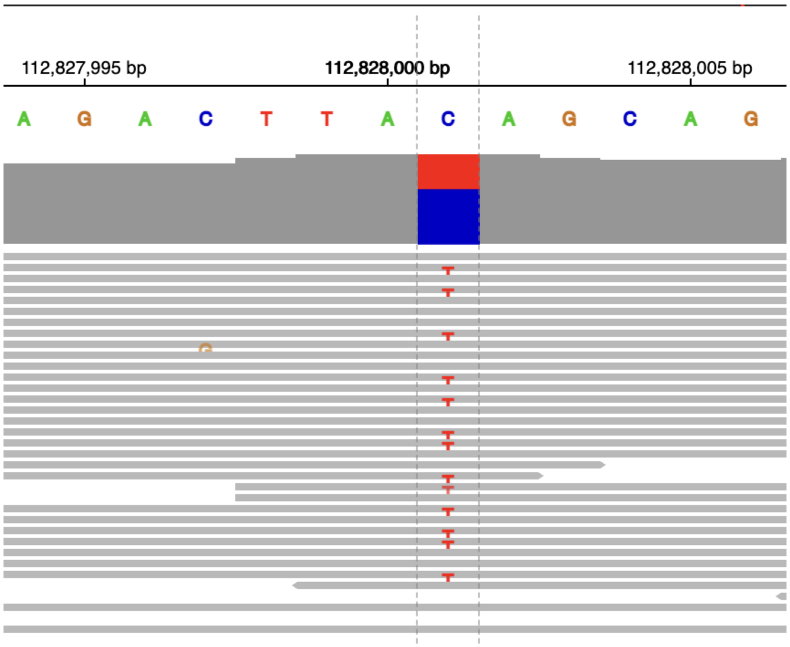

**Supplementary Figure 6. Germline APC mutation for patient G001.**
Integrative Genomics Viewer (IGV) showing supporting reads for germline *APC* mutation in patient G001.

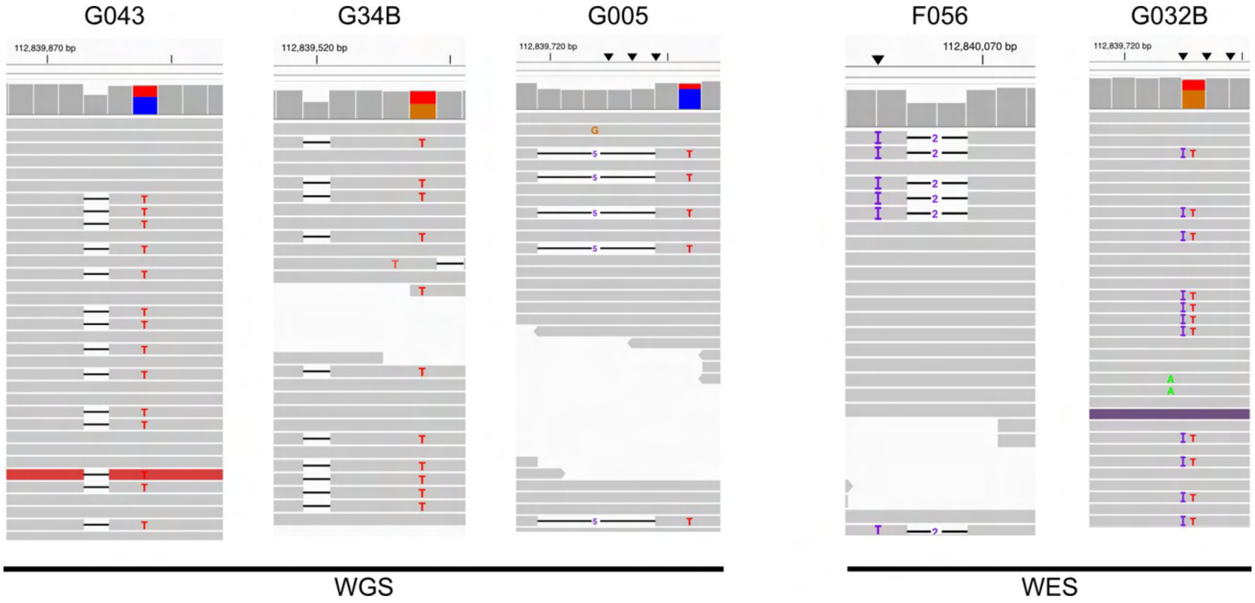

**Supplementary Figure 7. Clustered 2nd hit *APC* events.**
Integrative Genomics Viewer (IGV) showing supporting reads for samples containing clustered 2nd hit *APC* mutations defined as two or more SNV or INDELs within 10bp of one another, on chromosome 5 denoted in the main text (**Figure 1e**).

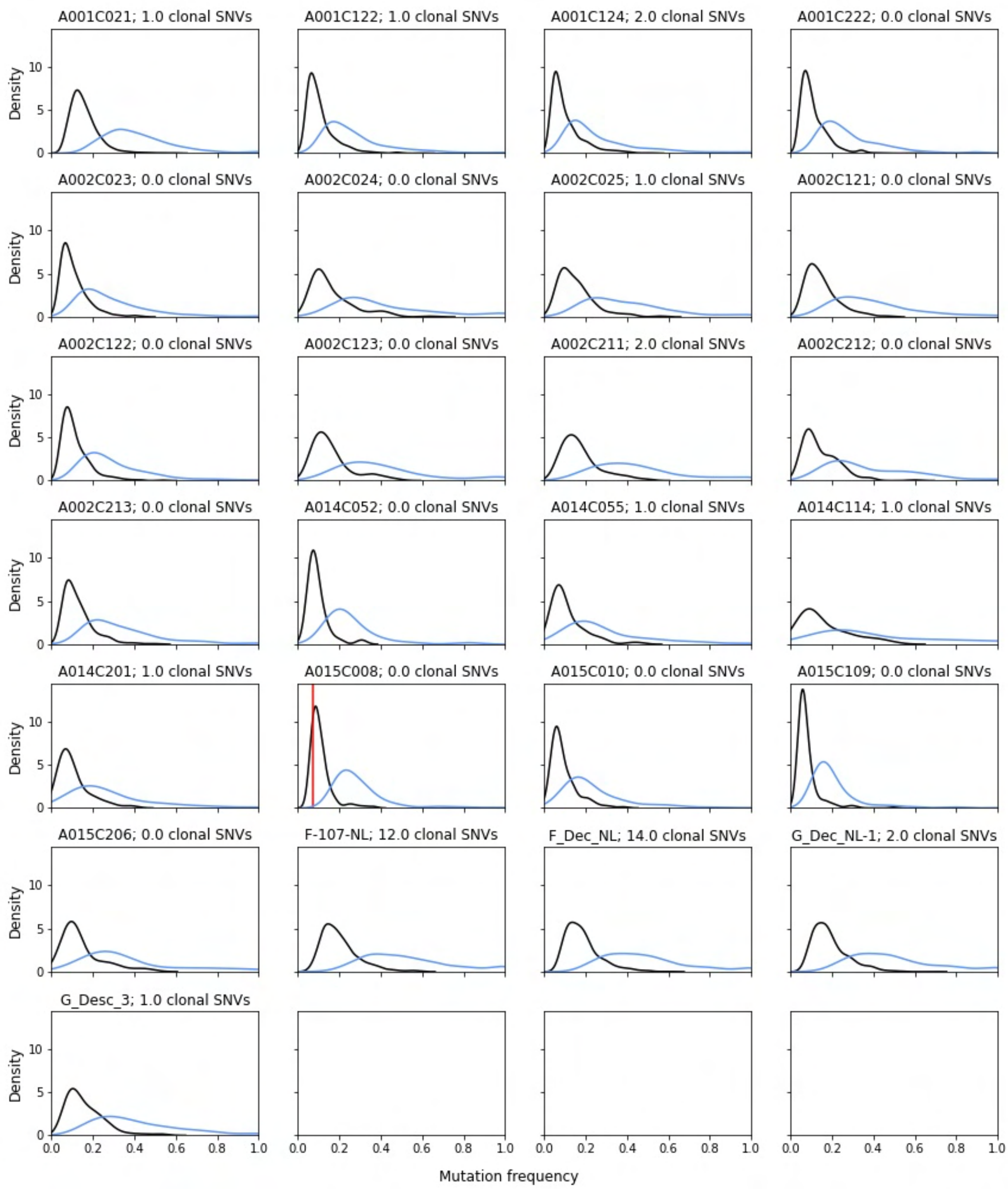

**Supplementary Figure 8. VAF distributions for WGS of normal mucosa.**

The black curve is the raw VAF distribution and the blue curve is the ppVAF distribution. Samples that have 2nd hit *APC* driver mutations have vertical lines in red at the VAF of the mutation(s), and samples with *KRAS* driver mutations have vertical lines at the VAFs in orange.

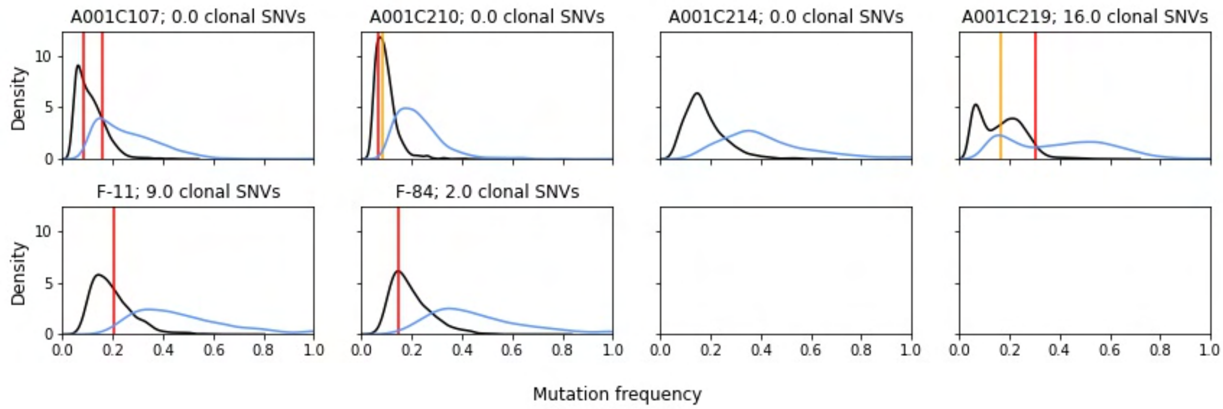

**Supplementary Figure 9. VAF distributions for WGS of benign polyclonal polyps.**
The black curve is the raw VAF distribution and the blue curve is the ppVAF distribution. Samples that have 2nd hit *APC* driver mutations have vertical lines in red at the VAF of the mutation(s), and samples with *KRAS* driver mutations have vertical lines at the VAFs in orange.

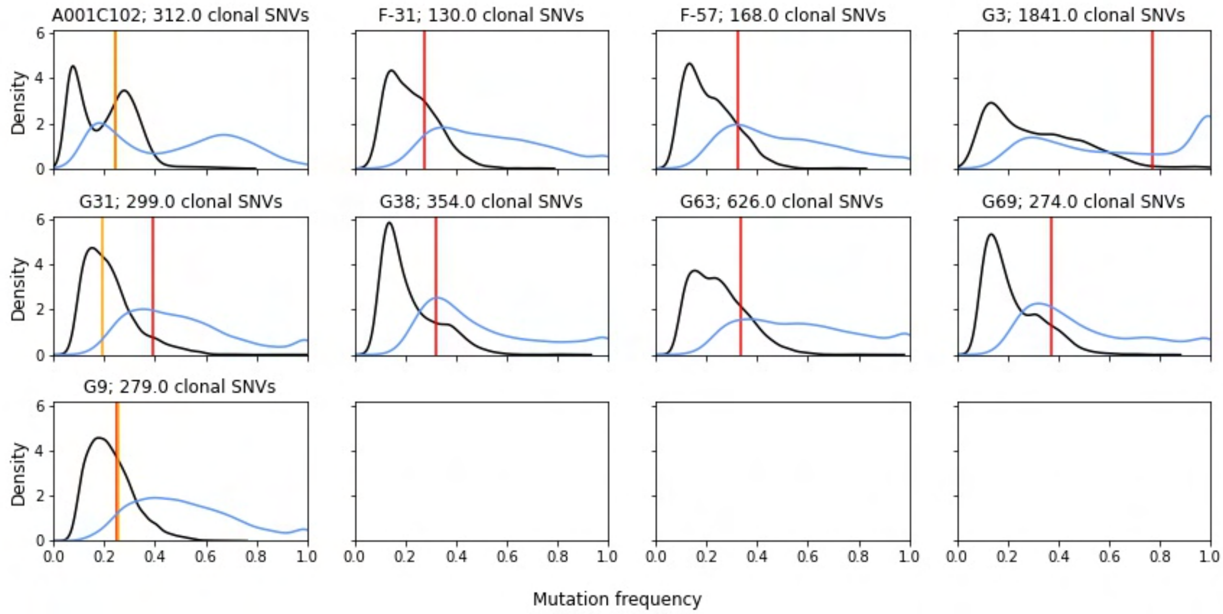

**Supplementary Figure 10. VAF distributions for WGS of benign monoclonal polyps.**
The black curve is the raw VAF distribution and the blue curve is the ppVAF distribution. Samples that have 2nd hit *APC* driver mutations have vertical lines in red at the VAF of the mutation(s), and samples with *KRAS* driver mutations have vertical lines at the VAFs in orange.

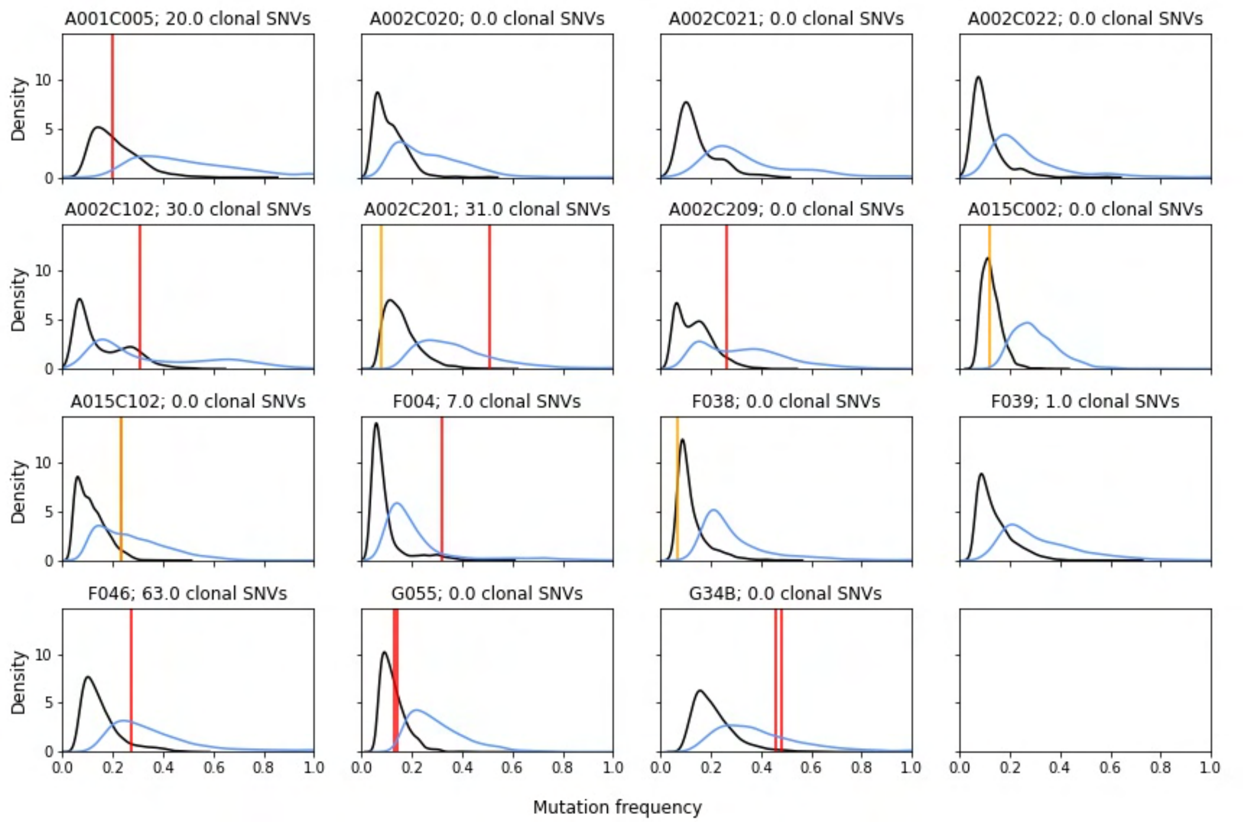

**Supplementary Figure 11. VAF distributions for WGS of dysplastic polyclonal polyps.**
The black curve is the raw VAF distribution and the blue curve is the ppVAF distribution. Samples that have 2nd hit *APC* driver mutations have vertical lines in red at the VAF of the mutation(s), and samples with *KRAS* driver mutations have vertical lines at the VAFs in orange.

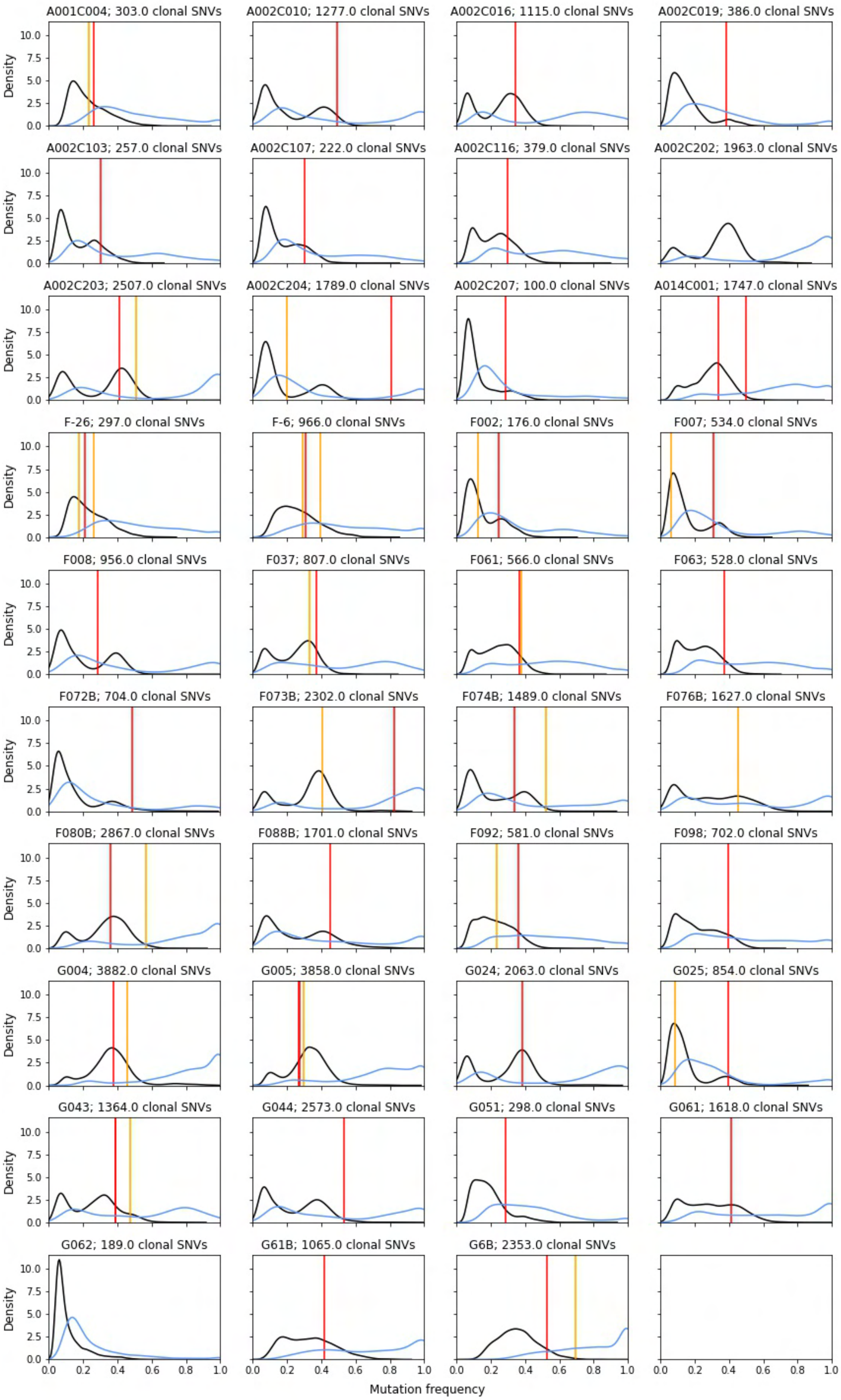

**Supplementary Figure 12. VAF distributions for WGS of dysplastic monoclonal polyps.**
The black curve is the raw VAF distribution and the blue curve is the ppVAF distribution. Samples that have 2nd hit *APC* driver mutations have vertical lines in red at the VAF of the mutation(s), and samples with *KRAS* driver mutations have vertical lines at the VAFs in orange.

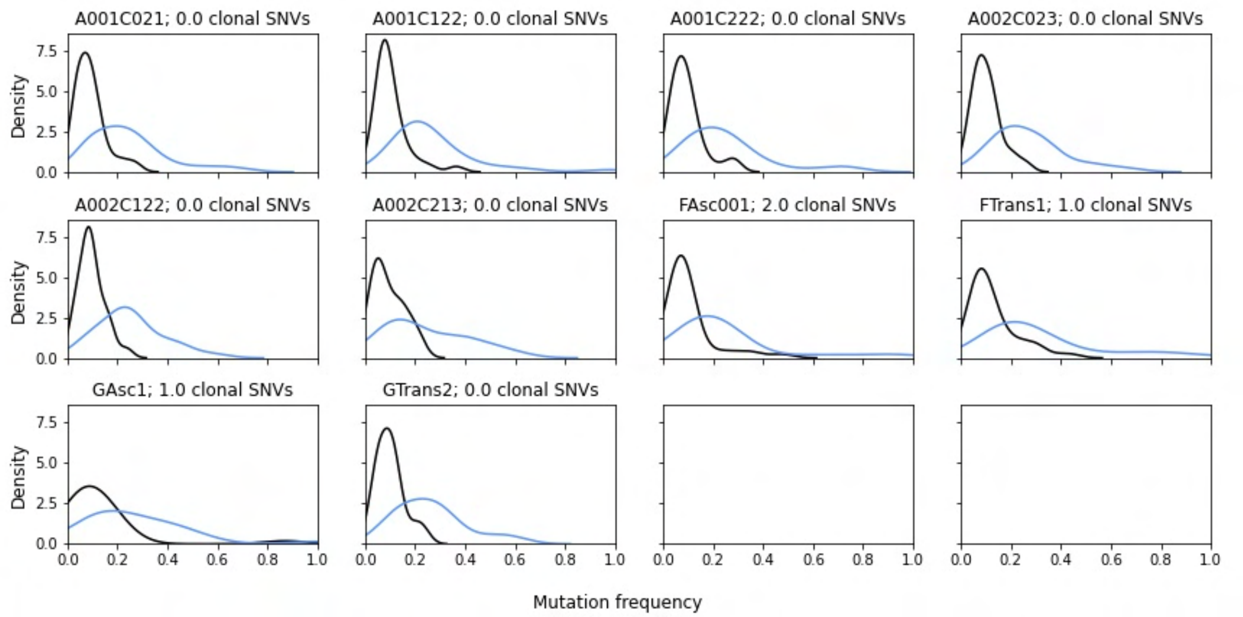

**Supplementary Figure 13. VAF distributions for WES of normal mucosa.**
The black curve is the raw VAF distribution and the blue curve is the ppVAF distribution. Samples that have 2nd hit *APC* driver mutations have vertical lines in red at the VAF of the mutation(s), and samples with *KRAS* driver mutations have vertical lines at the VAFs in orange.

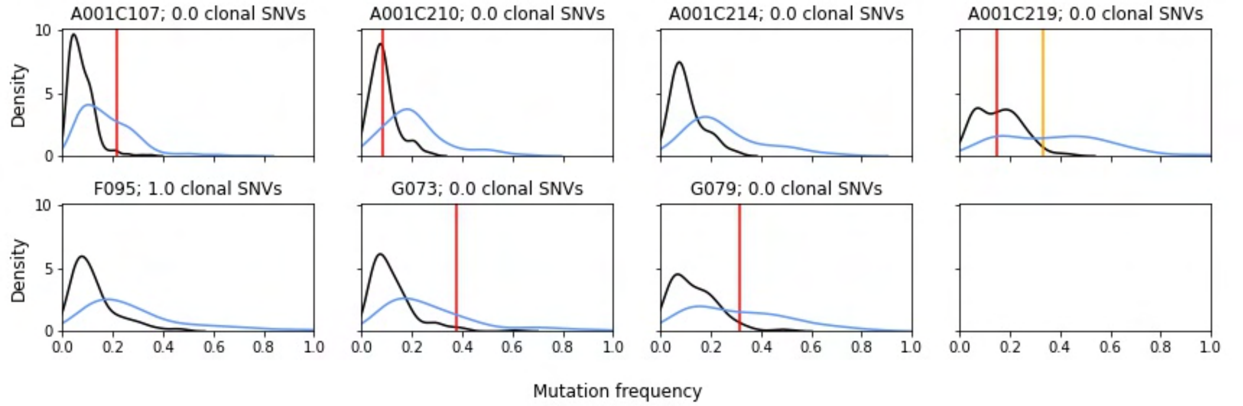

**Supplementary Figure 14. VAF distributions for WES of benign polyclonal polyps.**
The black curve is the raw VAF distribution and the blue curve is the ppVAF distribution. Samples that have 2nd hit *APC* driver mutations have vertical lines in red at the VAF of the mutation(s), and samples with *KRAS* driver mutations have vertical lines at the VAFs in orange.

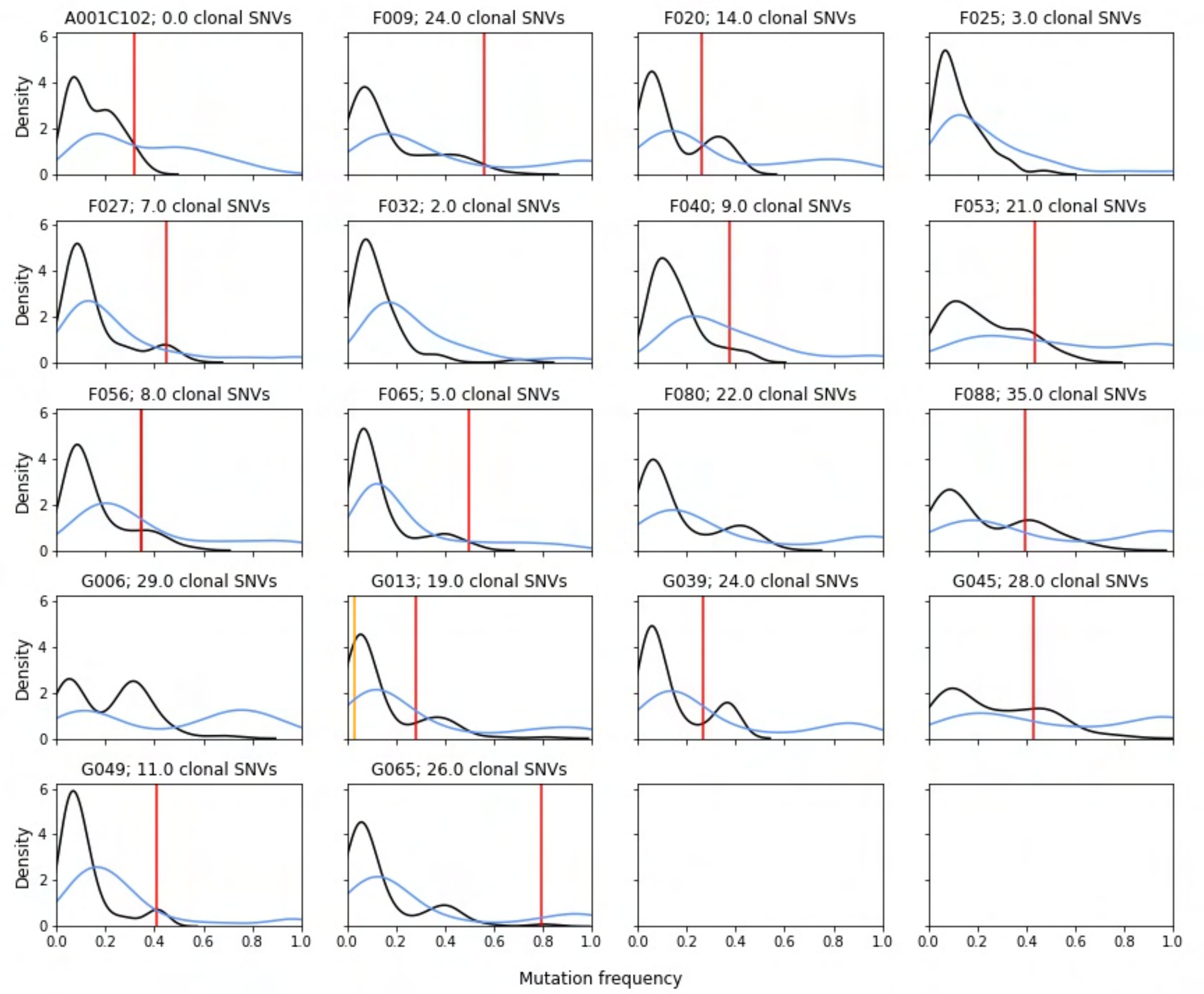

**Supplementary Figure 15. VAF distributions for WES of benign monoclonal polyps.**

The black curve is the raw VAF distribution and the blue curve is the ppVAF distribution. Samples that have 2nd hit *APC* driver mutations have vertical lines in red at the VAF of the mutation(s), and samples with *KRAS* driver mutations have vertical lines at the VAFs in orange.

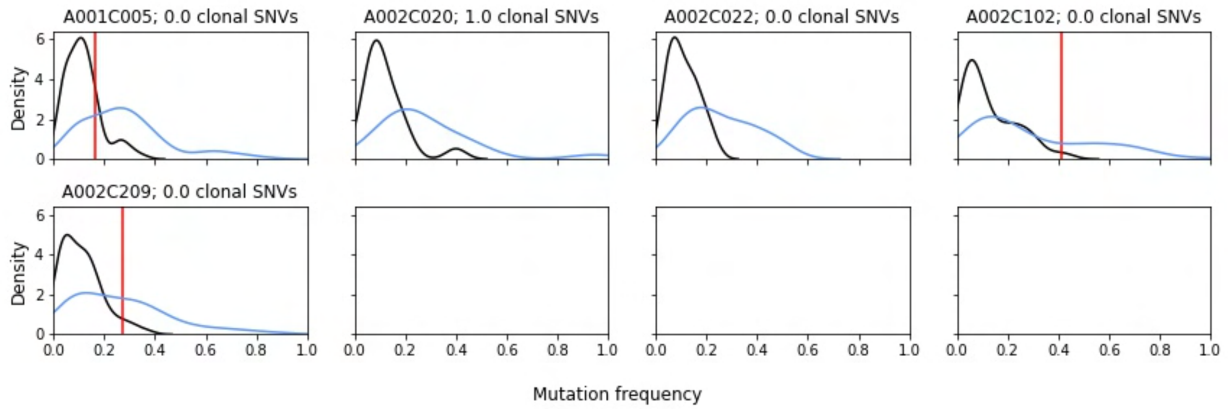

**Supplementary Figure 16. VAF distributions for WES of dysplastic polyclonal polyps.**
The black curve is the raw VAF distribution and the blue curve is the ppVAF distribution. Samples that have 2nd hit *APC* driver mutations have vertical lines in red at the VAF of the mutation(s), and samples with *KRAS* driver mutations have vertical lines at the VAFs in orange.

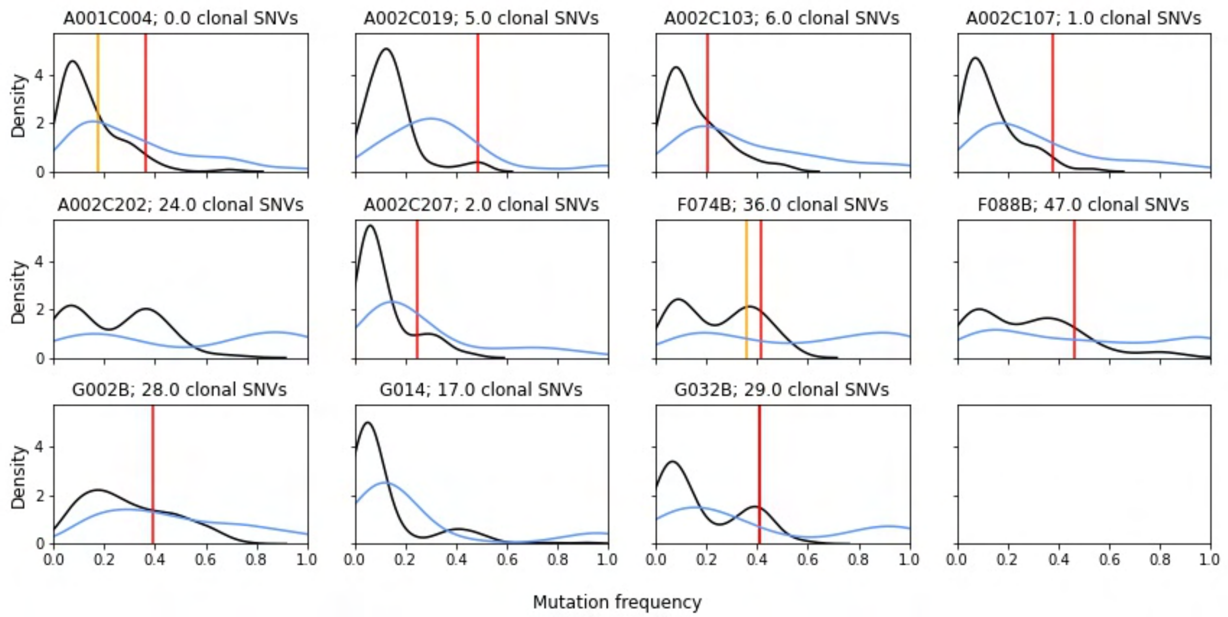

**Supplementary Figure 17. VAF distributions for WES of dysplastic monoclonal polyps.**
The black curve is the raw VAF distribution and the blue curve is the ppVAF distribution. Samples that have 2nd hit *APC* driver mutations have vertical lines in red at the VAF of the mutation(s), and samples with *KRAS* driver mutations have vertical lines at the VAFs in orange.

### Supplementary Note 1: Estimating the number of clonal mutations from bulk sequencing data

To determine whether a bulk sequenced sample is monoclonal or polyclonal, we estimated the number of clonal SNVs present in the sample. To do this, we determined whether each SNV in a sample was likely clonal or subclonal based on its reference and mutant read counts. However, determining whether a mutation is clonal or subclonal can be difficult, since truly clonal mutations can sometimes stochastically produce reference and mutant read counts similar to those produced by truly subclonal mutations (**Fig. SN1.1**). As explained in the **Methods**, we assumed the mutant read count $t_{alt}$ is binomially distributed

$$t_{alt} \sim Binom(t_{depth}, p), \text{ where } p = CN_{mut}\, \rho f / (2(1-\rho) + \rho CN_{total})$$

given the total sequencing depth at the locus $t_{depth}$, the total copy number and mutant copy number at the locus $CN_{total}$ and $CN_{mut}$, the ppVAF value $f$, and a purity value ρ. Discriminating between clonal and subclonal mutations becomes even more difficult at lower sequencing depths, in samples where subclonal mutations are at higher frequency, and at lower sample purities (**Fig. SN1.1**).

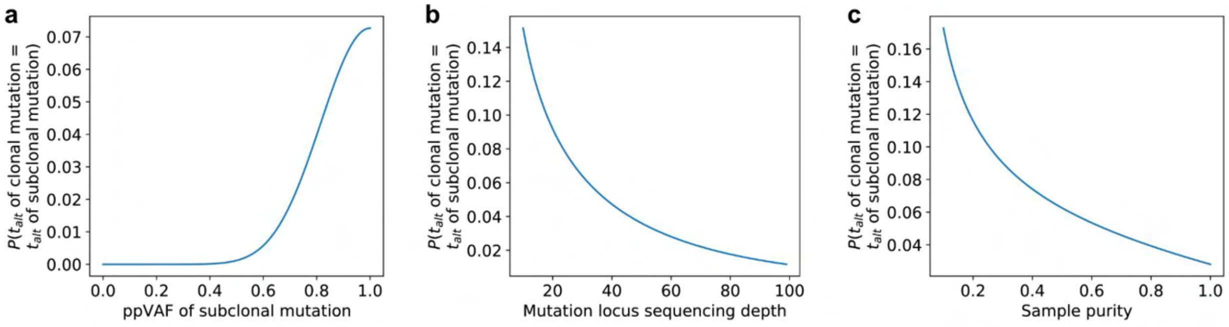

**Figure SN1.1. Overlap of binomial mutant read count probability mass functions between subclonal and clonal mutations. a.** Probability that a clonal mutation and a subclonal mutation with true ppVAF given by the x-axis value yield the same number of mutant reads when sequenced. The total sequencing depth is assumed to be 60 reads and the sample purity is assumed to be 100%. **b.** Probability that a clonal mutation and a subclonal mutation with true ppVAF 0.75 yield the same number of mutant reads when sequenced to the depth given on the x-axis. The sample purity is assumed to be 100%. **c.** Probability that a clonal mutation and a subclonal mutation with true ppVAF 0.75 yield the same number of mutant reads when the sample purity is equal to the value on the x-axis. The total sequencing depth is assumed to be 60 reads.

Ideally, we would use some function of the posterior ppVAF probability density $P(ppVAF = f \mid t_{alt})$ to determine whether a mutation is clonal or subclonal, since if we collect an infinite number of sequencing reads this posterior density should converge to 1 at the true ppVAF value ($f$=1 for clonal mutations, $f$<1 for subclonal mutations). However, at lower sequencing depths, this posterior density distribution is more diffuse, representing the uncertainty about the true ppVAF and clonal/subclonal status of the mutation (**Fig. SN1.2a**). Therefore, for finite sequencing depths, truly clonal mutations will have some posterior probability density below ppVAF=1 and truly subclonal mutations will have some posterior density at 1. This means that while the naive estimator for the clonal mutation count $\sum_{all\ muts} \lim_{\varepsilon \to 1} P(ppVAF \leq \varepsilon \mid t_{alt})$ is asymptotically consistent with respect to the sequencing depth, in practice it can perform poorly

(**Fig. SN1.2b-c**). In general the entire class of estimators $\sum_{all\ muts} P(ppVAF \leq \varepsilon \mid t_{alt})$ performs poorly with realistic sequencing depths for any threshold $\varepsilon$ due to this uncertainty, often underestimating the number of clonal mutations at higher values of $\varepsilon$ and overestimating at lower values (though the error depends on the true number of clonal and subclonal mutations present).

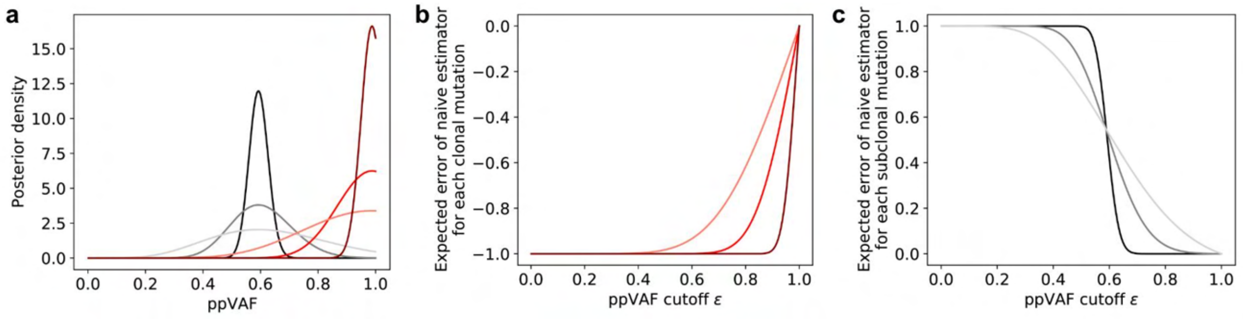

**Figure SN1.2. ppVAF posterior density overlap and estimator error at finite sequencing depths. a.** ppVAF posterior densities of clonal (shades of red) and subclonal mutations with true ppVAF = 0.6 (shades of grey). Mutations were simulated at total sequencing depths {25, 100, 1000}, with increasing depths corresponding to increasing color intensity. **b-c.** The expected contribution of each clonal (**b**) and subclonal (**c**) mutation to the naive estimator of clonal mutation count that sums the ppVAF posterior density of each mutation above a threshold $\varepsilon$. As in panel **a**, increasing sequencing depths correspond to increasing color intensity, with values {25, 100, 1000}.

To address this issue, we constructed an estimator of the clonal mutation count that was not only asymptotically consistent, but unbiased at all sequencing depths (i.e., $E[\hat{n}_{clonal}] = n_{clonal}$ for the estimator $\hat{n}_{clonal}$ and the true value $n_{clonal}$). We used an estimator of the form $\hat{n}_{clonal} = \sum_{i \in all\ muts} z_i$ where $z_i = 1$ iff mutation *i* has $P(ppVAF > \varepsilon) > c$, essentially classifying all mutations with cumulative posterior ppVAF density $P(ppVAF > \varepsilon)$ greater than some threshold *c* as clonal. This estimator is unbiased only at specific values of *c*, which we can calculate from its expected value

$$
\begin{aligned}
\mathbb{E}[\hat{n}_{\text{clonal}}] &= \mathbb{E}[\sum_{i \in \text{all muts}} z_i] \\
&= n_{\text{clonal}} \mathbb{E}[z_i | i \text{ is clonal}] + n_{\text{subclonal}} \mathbb{E}[z_i | i \text{ is subclonal}] \\
&= n_{\text{clonal}} P(z_i = 1 | i \text{ is clonal}) + n_{\text{subclonal}} P(z_i = 1 | i \text{ is subclonal}) \\
&= n_{\text{clonal}} (P(z_i = 1 | i \text{ is clonal}) + \frac{n_{\text{subclonal}}}{n_{\text{clonal}}} P(z_i = 1 | i \text{ is subclonal}))
\end{aligned}
$$

From the last line above, this expected value is equal to the true value $n_{clonal}$ when

$$n_{\text{clonal}}(1 - P(z_i = 1 | i \text{ is clonal})) = n_{\text{subclonal}} P(z_i = 1 | i \text{ is subclonal}).$$

This condition occurs when the contribution to the estimator from false negative clonal classifications (mutations that are truly clonal but are classified as subclonal based on their ppVAF posterior density; occurs with frequency $P(z_i = 0 \mid i \text{ is clonal})$ and expressed on the left-hand side of the above equation) exactly balances that of the false positive classifications (mutations that are truly subclonal but are classified as clonal; right-hand side of above equation).

We can tune $P(z_i = 1 \mid i$ is clonal) and $P(z_i = 1 \mid i$ is subclonal) by adjusting the value of the cutoff *c* to satisfy the above equation and make the estimator $\hat{n}_{clonal}$ unbiased. However, to use this strategy we must approximate some ground truth values that we don't know *a priori*:

1. $P(z_i = 1 \mid i$ is clonal) and $P(z_i = 1 \mid i$ is subclonal) for any cutoff *c*, which require posterior ppVAF densities of known true clonal and true subclonal mutations. These values will depend on the sequencing depth and underlying ppVAF distributions of true subclonal mutations in the dataset.
2. The true number of clonal mutations $n_{clonal}$ in a sample- exactly the quantity we are trying to estimate!

We addressed point (1) by generating approximate ground truth sequencing data from which to estimate $P(z_i = 1 \mid i$ is clonal) and $P(z_i = 1 \mid i$ is subclonal) which match the sequencing depth of the dataset (**Fig. SN1.3a**). $P(z_i = 1 \mid i$ is clonal) was estimated from synthetic datasets in which true diploid clonal mutations (ppVAF = 1) were binomially simulated, given the observed distribution of sequencing depths at mutation loci and the observed scATAC-seq sample purities (same distribution as used to estimate ppVAFs). In other words, the mutant read count $t'_{alt}$ was simulated by drawing from $t'_{alt} \sim Binom(t'_{depth}, p')$ where $t'_{depth}$ is drawn from the observed sequencing depth distribution, $p' = \rho'/2((1-\rho') + \rho')$, and $\rho'$ is drawn from the observed scATAC-seq purity distribution. A set of true subclonal mutations was approximated by using the read counts for somatic mutations found in the mucosal samples. Most mutations in the polyclonal normal mucosa should be subclonal, and using mucosal data from the same dataset ensures that sequencing depth and other key parameters are preserved. For both the simulated clonal and ground truth subclonal mutations, the depths and mutant read counts were used to estimate ppVAF posteriors as described in the **Methods**. $P(z_i = 1)$ for true subclonal and clonal mutations was then empirically estimated from these ground truth datasets by computing the frequency of mutations with $P(ppVAF > \varepsilon) > c$ given any cutoff values for *c* and $\varepsilon$.

To address point (2), we employed an iterative procedure to reduce the bias in the estimator as much as possible by estimating the number of clonal mutations in each sample using an initial value for the threshold *c* (conceptually similar to the expectation step of an expectation-maximization algorithm), and using this estimate of $n_{clonal}$ to update the value of *c* (similar to the maximization step). We repeated this cutoff optimization until the estimated number of clonal mutations did not change further.

This procedure is guaranteed to minimize the expected error in $n_{clonal}$

$$\mathbb{E}[\hat{n}_{\text{clonal}} - n_{\text{clonal}}] = n_{\text{subclonal}} P(z_i = 1 | i \text{ is subclonal}) - n_{\text{clonal}} P(z_i = 0 | i \text{ is clonal}))$$

and converge on the true value of $n_{clonal}$. The algorithmic correctness of this procedure can be proved by showing that both steps bring the estimator value closer to the true number of clonal mutations.

**STEP 1: update the estimated number of clonal mutations**

We will show that, given an initial classification threshold computed from the previous estimated number of clonal mutations $\hat{n}_{clonal}$ that differs from the true value $n_{clonal}$ by $\sigma = \hat{n}_{clonal} - n_{clonal}$ mutations, the expected updated estimate $\hat{n}'_{clonal}$ will be closer to the true number of clonal mutations (i.e., $|\hat{n}'_{clonal} - n_{clonal}| < |\hat{n}_{clonal} - n_{clonal}|$ ).

$$
\begin{aligned}
\mathbb{E}[\hat{n}'_{\text{clonal}}] &= n_{\text{clonal}}\mathbb{E}[z_i|i \text{ is clonal}] + n_{\text{subclonal}}\mathbb{E}[z_i|i \text{ is subclonal}] \\
&= (\hat{n}_{\text{clonal}} - \sigma)P(z_i = 1|i \text{ is clonal}) + (\hat{n}_{\text{subclonal}} + \sigma)P(z_i = 1|i \text{ is subclonal}) \\
&= \left(\hat{n}_{\text{clonal}}P(z_i = 1|i \text{ is clonal}) + \hat{n}_{\text{subclonal}}P(z_i = 1|i \text{ is subclonal})\right) \\
&\quad + \sigma(P(z_i = 1|i \text{ is subclonal}) - P(z_i = 1|i \text{ is clonal})) \\
&= \hat{n}_{\text{clonal}} + \sigma(P(z_i = 1|i \text{ is subclonal}) - P(z_i = 1|i \text{ is clonal}))
\end{aligned}
$$

The value of ($P(z_i = 1 \mid i$ is subclonal) - $P(z_i = 1 \mid i$ is clonal)) should be negative (i.e., a true clonal mutation is more likely to be classified as clonal than a true subclonal mutation) if the ground truth subclonal and clonal mutations used to tune these values were chosen appropriately. Therefore, the new expected value of $\hat{n}'_{clonal}$ will be less than the initial estimate if $\sigma > 0$ and $\hat{n}'_{clonal} > \hat{n}_{clonal}$ if $\sigma < 0$, indicating that $|\hat{n}'_{clonal} - n_{clonal}| < |\hat{n}_{clonal} - n_{clonal}|$.

**STEP 2: update the classification threshold**

As stated above, the classification threshold is chosen to provide the new values of $P'(z_i = 1 \mid i$ is subclonal) and $P'(z_i = 1 \mid i$ is clonal)) that best satisfy

$$
\hat{n}'_{\text{clonal}}(1 - P'(z_i = 1|i \text{ is clonal})) = \hat{n}'_{\text{subclonal}}P'(z_i = 1|i \text{ is subclonal})
$$

given the new estimated number of clonal mutations $\hat{n}'_{clonal}$. Assuming that the estimated number of clonal mutations changed by $\sigma' = \hat{n}'_{clonal} - \hat{n}_{clonal}$ in the previous iteration of **STEP 1**, this condition is equivalent to

$$
\begin{aligned}
&\hat{n}_{\text{clonal}}\left(P'(z_i = 0|i \text{ is clonal}) - P(z_i = 0|i \text{ is clonal})\right) \\
&\quad - \hat{n}_{\text{subclonal}}\left(P'(z_i = 1|i \text{ is subclonal}) - P(z_i = 1|i \text{ is subclonal})\right) \\
&\qquad = -\sigma'\left(P'(z_i = 0|i \text{ is clonal}) + P'(z_i = 1|i \text{ is subclonal})\right).
\end{aligned}
$$

The right-hand side of the above equation always has the same sign as $\sigma'$. Denoting $P'(z_i = 1 \mid i$ is subclonal) - $P(z_i = 1 \mid i$ is subclonal) as $\Delta P(z_i = 1 \mid i$ is subclonal) and $P'(z_i = 0 \mid i$ is clonal) - $P(z_i = 0 \mid i$ is clonal) as $\Delta P(z_i = 0 \mid i$ is clonal), we therefore have

$$
\begin{aligned}
&\hat{n}_{\text{clonal}}\Delta P(z_i = 0|i \text{ is clonal}) > \hat{n}_{\text{subclonal}}\Delta P(z_i = 1|i \text{ is subclonal}) \text{ if } \sigma' < 0 \\
&\hat{n}_{\text{clonal}}\Delta P(z_i = 0|i \text{ is clonal}) < \hat{n}_{\text{subclonal}}\Delta P(z_i = 1|i \text{ is subclonal}) \text{ if } \sigma' > 0.
\end{aligned}
$$

$\Delta P(z_i = 0 \mid i$ is clonal) and $\Delta P(z_i = 1 \mid i$ is subclonal) always have opposite signs. This is because the value of $P(z_i = 1) = P(ppVAF > \varepsilon) > c$ for both subclonal and clonal mutations is set by adjusting the value of the cutoff *c.* Increasing *c* will classify fewer mutations as clonal and therefore lower the value of $P(z_i = 1)$ and increase $P(z_i = 0)$ for all mutations, while decreasing *c* will do the opposite. Thus, when $\sigma' < 0$, $\Delta P(z_i = 0 \mid i$ is clonal) must be positive and $\Delta P(z_i = 1 \mid i$ is subclonal) must be negative, increasing *c* and lowering the number of mutations classified as clonal in the next iteration. Similarly, when $\sigma' > 0$, *c* decreases and the number of mutations classified as clonal in the next iteration increases. Therefore, upon a single iteration of both steps, the threshold value *c* is updated such that the estimated number of clonal mutations gets closer than the true value, and repeating the steps until convergence of $\hat{n}_{clonal}$ leads to an unbiased and consistent estimate of $n_{clonal}$.

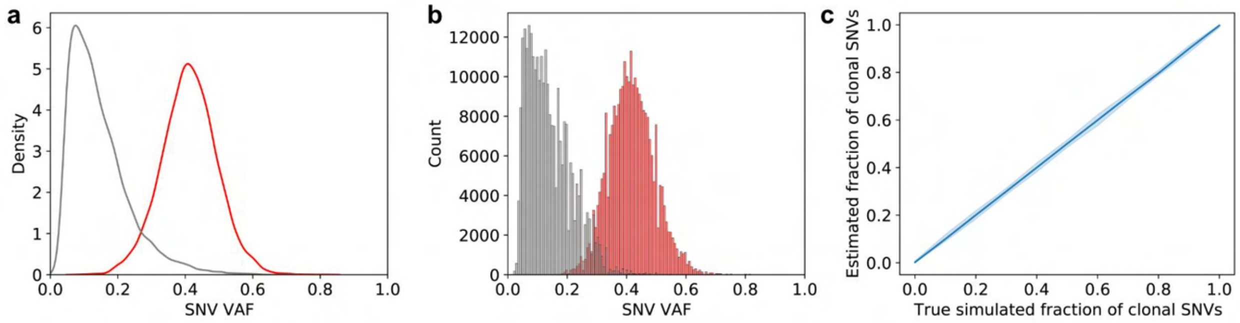

**Figure SN1.3. VAFs and estimated classifications of ground truth clonal and subclonal mutations.** **a.** VAF distributions of ground truth simulated clonal SNVs (red) and subclonal SNVs taken from mucosal WGS data (grey). **b.** VAF distributions of ground truth data classified by the iterative procedure described above into prospective clonal SNVs (red) and subclonal SNVs (grey). **c.** Comparison of the true fraction and estimated fraction of clonal mutations of 1000 SNV datasets simulated by subsampling the true clonal and subclonal mutation sets. Error ribbon is the 95% percentile interval, 50 simulated datasets per condition.

This algorithm leads to good estimates of the number of clonal mutations in datasets resampled from our ground truth clonal and subclonal mutations used to construct the estimator (**Fig. SN1.3b-c**). We can also compare these estimates and the clonal/subclonal mutation classifications to a procedure that uses k-means clustering on mutation VAFs to identify clonal mutations (**Fig. SN1.4**). These two strategies seem to both perform well and produce similar clonal/subclonal classifications in samples with a visible clonal peak at VAF ~ 0.4 (**Fig. SN1.4a-b**). However, unsupervised clustering of mutations cannot reliably produce sensible clonal/subclonal mutation classifications for samples with only one major peak in the VAF distribution, making it unsuitable for determining whether samples are polyclonal or monoclonal more generally (**Fig. SN1.4c-d**). Overall, our ppVAF-based procedure iterative produces estimates of $n_{clonal}$ with clear clonal peaks that are similar to the clustering results, except for samples with a second peak at a lower VAF than would be expected for clonal mutations (**Fig. SN1.4e-f**). While the clustering strategy simply considers all mutations in the higher-VAF peak as clonal, our iterative classification procedure uses the absolute ppVAF values to perform the classification, resulting in lower clonal mutation counts (**Fig. SN1.4e**). In these cases, the true clonality of the mutations in this second VAF peak is unclear, since there are two possibilities that are indistinguishable without additional information on the purity of the sample:

1. Overestimation of the purity of the sample when using the cohort-level purity distribution to estimate the ppVAF.
2. Presence of a major subclone that is present in the sample at a truly-subclonal frequency. Additional epithelial subclones are also present, and the sample is indeed polyclonal.

While this uncertainty may contribute to some error in our polyclonal classifications for samples in our bulk sequencing datasets, we do not believe that it explains all of the evidence for polyclonality in our premalignant samples (explored more in **Supplementary Note 2**).

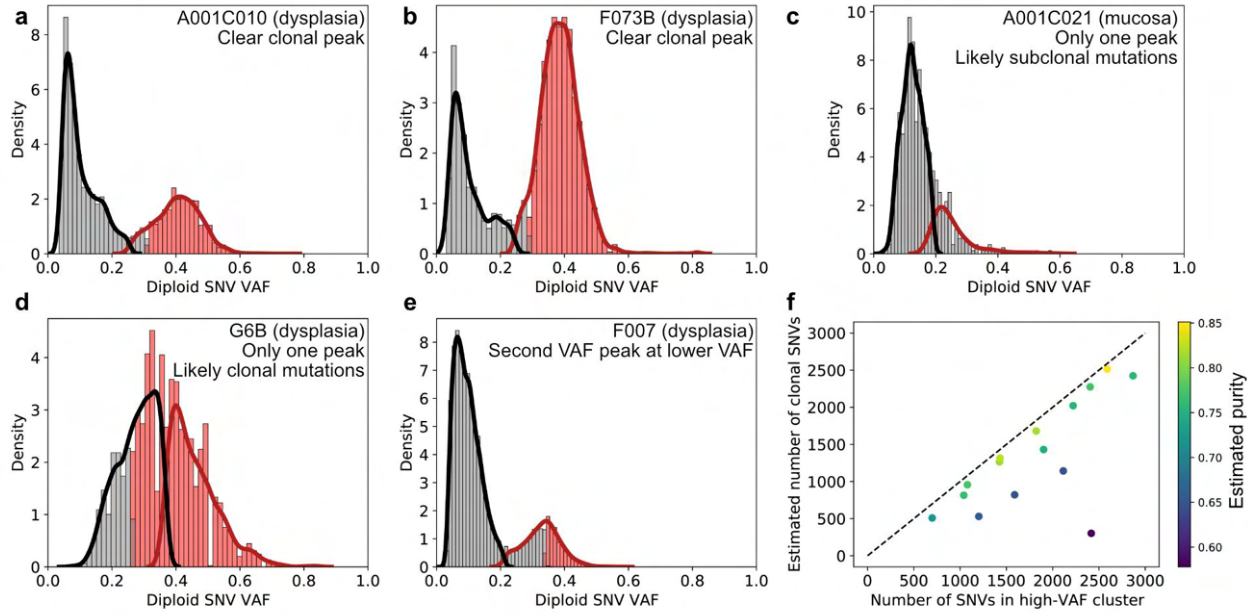

**Figure SN1.4. Estimated clonal and subclonal diploid mutation classifications of HTAN WGS samples. a-e.** Comparison between k-means clustering classifications on VAF values (KDE lines) and our clonal classification strategy (histograms) on WGS samples with clear high-VAF clonal peaks (**a-b**), only one visible VAF peak (**c-d**), or a distinct lower VAF peak that may or may not represent clonal mutations (**e**). **f.** Comparison between the clustering-estimated number of diploid SNVs (x-axis) and our estimated number of diploid SNVs (y-axis) for WGS samples with visible clonal mutation peaks. Point color indicates the estimated purity of the sample, assuming the high-VAF mutation peak is clonal (estimated as 2*mean VAF of high-VAF cluster). Perfect concordance between estimates is shown by the dashed line.

In addition, we empirically found our clonal mutation count estimation procedure also approximately minimizes the expected error of the individual clonal/subclonal mutation classifications (**Fig. SN1.5a-b**). Therefore, we used these optimized clonal/subclonal classifications to identify individual driver mutations that were clonal or subclonal in **Fig. 1e-f**.

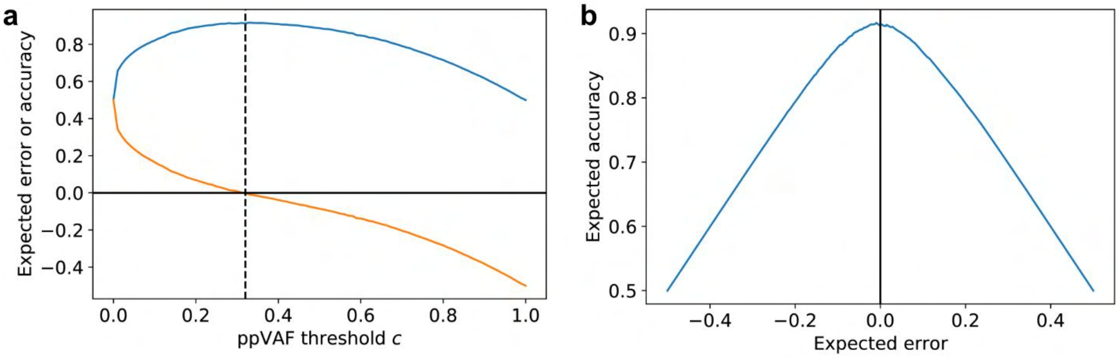

**Figure SN1.5. Expected accuracy of clonal/subclonal mutation classifications with respect to the ppVAF classification threshold and expected error of the clonal mutation count estimate. a.** Expected accuracy (blue curve) of the clonal/subclonal classifications for different clonal classification thresholds *c*, estimated as $n_{subclonal}/n_{total}P(z_i = 0 \mid i$ is subclonal) + $n_{clonal}/n_{total}P(z_i = 1 \mid i$ is clonal), assuming half of mutations are clonal. The expected error of the clonal mutation count estimate (orange curve) is also plotted, along with the optimal ppVAF threshold *c* (dashed line). The expected classification accuracy appears to peak at the optimal ppVAF threshold *c.* **b.** Expected classification accuracy (y-axis) at different values of the expected error in $\hat{n}_{clonal}$ (y-axis). The optimal cutoff used for clonal/subclonal classifications is at expected error = 0.

#### Supplementary Note 2: Effect of purity misestimation on polyclonal/monoclonal classifications from bulk WGS data

Our procedure for estimating the initiation time and thereby classifying bulk samples as polyclonal or monoclonal relies on estimating the number of clonal mutations in a sample. As explained in **Supplementary Note 1**, we do this by thresholding on the ppVAF value for each somatic SNV to classify it as clonal or subclonal. However, this strategy requires us to accurately correct the raw VAF for sample purity to estimate the fraction of epithelial cells in a sample with the mutation. In most cancer samples, this is done by inferring the sample purity using the copy number profile. Unfortunately this method is not appropriate for our samples, since they are largely premalignant samples where not all epithelial cells have clonal copy number changes due to an overall lack of copy number changes and/or possible polyclonality (**Methods**, **Extended Data Fig. 4**).

Therefore, we used the epithelial cell fraction estimated from scATAC-seq data from the same HTAN FAP patient cohort[28] to correct for sample purity (**Methods**). These samples are taken mostly from patients who were also profiled in this study (A001: 14 samples, A002: 15 samples, A014: 13 samples, A015: 15 samples, F001: 4 samples), though we used an additional 7 polyp samples from other FAP patients in this study cohort (**Supplementary Table SN2.1**). Since most samples do not have a matched sample with scATAC-seq data taken from the same mucosal region or polyp, we used the distribution of epithelial cell fractions taken from the scATAC-seq cohort for all samples to correct for sample purity. This strategy avoids the pitfalls associated with copy number based algorithms for estimating purity in individual samples using the bulk WGS or WES data, but introduces a source of error in the ppVAF estimates. Samples with a true epithelial cell fraction less than the expected scATAC-seq sample purity will have somatic mutation ppVAF estimates that are generally lower than the true ppVAF (and vice versa). This error can lead to undercounting clonal mutations in these lower-purity samples, leading to false polyclonal classifications.

| | All HTAN FAP scATAC-seq samples (used for ppVAFs) | scATAC-seq samples from the 6 patients in the HTAN WGS data | scATAC-seq samples with a matched WGS sample |
|---|---|---|---|
| Mucosa | 18 | 18 | 12 |
| Polyp (benign or dysplastic) | 48 | 41 | 13 |
| AdCa | 6 | 2 | 2 |

**Table SN2.1. Number of HTAN FAP samples with scATAC-seq data used to estimate the purity distributions and overlap with HTAN WGS data.**

To estimate an upper bound for the false positive polyclonal classification rate, we simulated lower-purity monoclonal samples by computationally lowering the purity of HTAN WGS polyp samples we classified as monoclonal in our original analysis. For each simulated lower-purity sample, we randomly chose a monoclonal polyp sample and lowered the VAF of each mutation by increasing the number of reference reads while preserving the original total number of reads at each locus. We then randomly chose a mucosal sample to provide mutations from the additional non-epithelial cells that would also be present in a lower-purity sample, lowering the effective purity of these mutations by adding reference reads as for the monoclonal polyp sample. Adding new mutations from a separate mucosal sample doesn't exactly recapitulate the effect of increasing the fraction of non-epithelial cells in a sample, which would be better represented by increasing the VAFs of existing mutations in the non-epithelial cells in the monoclonal sample rather than mixing in mutations from another sample altogether. However, as we cannot determine which mutations are present in the epithelial or non-epithelial cells in bulk sequencing, we must approximate this effect by mixing in polyclonal mutations from another sample. After artificially lowering these purities, we combined the mutations from the monoclonal sample and the normal sample, filtered out mutations with VAF < 0.01 as in our standard filtering pipeline, and used our clonal SNV counting method to determine whether these simulated samples would be called monoclonal or polyclonal.

We found that lower-purity samples are indeed more likely to be called polyclonal, with nearly all of these simulated samples with purity lowered by 30% or more being classified as polyclonal (**Fig. SN2.1a**). This effect is due to the reduction in ppVAF of the mutations previously considered clonal in these samples, leading to a decrease in the number of estimated clonal SNVs in the sample.

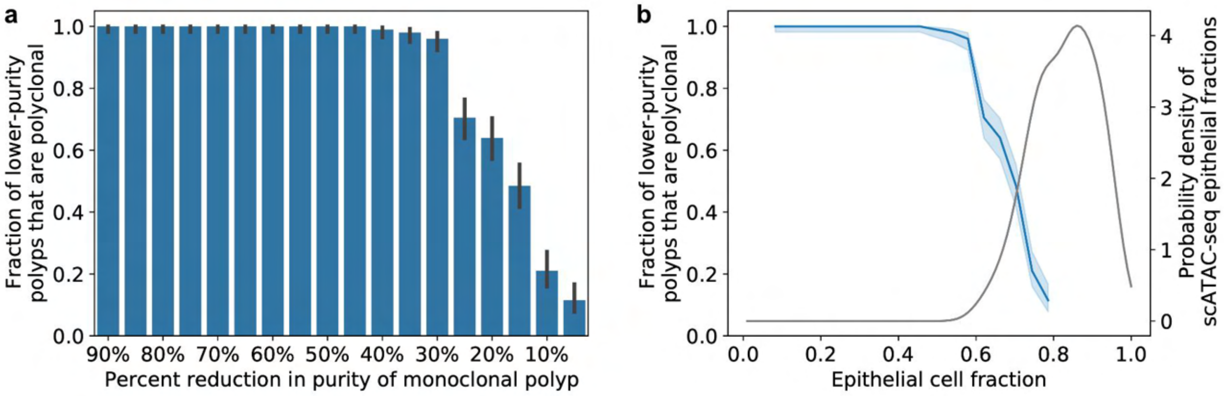

**Figure SN2.1. Simulations show that samples with lower purity cannot account for the degree of polyclonality observed in our bulk WGS data. a.** Fraction of simulated monoclonal polyps with sample purity computationally reduced that are classified as polyclonal. Error bars are 95% Bayesian CIs, n=200 simulation runs per condition. **b.** Fraction of simulated monoclonal polyps at different effective sample purities that are called polyclonal (blue curve with left y-axis, simulation data same as in panel **a**, error ribbon is 95% Bayesian CI). This curve was convolved with the scATAC-seq measured sample purity distribution for polyps (grey curve, right y-axis) to compute the upper bound for the expected fraction of true monoclonal samples that will be misclassified as polyclonal due to low purity.

To calculate the overall effect of these lower-purity samples on our estimated polyclonal frequency, we can estimate how often samples at purities below the mean purity truly occur. Assuming the scATAC-seq data provides a good approximation for the distribution of sample purities, we can estimate the frequency of samples with a given epithelial cell fraction using the smoothed distribution for polyp samples (the same one used for ppVAF estimation). We observed that samples with purities that result in mostly polyclonal classifications are relatively unlikely according to the scATAC-seq measured distribution (**Fig. SN2.1b**). We calculated the expected frequency of false polyclonal classifications, assuming the scATAC-seq purity distribution $P(\rho)$ represents the true distribution well, by approximating

$$\int_0^{\bar{\rho}} P(polyclonal \mid \rho) P(\rho) d\rho$$

using trapezoidal sums. The expected effective sample purity $\rho$ was computed for the simulated samples by multiplying the mixing fraction of the monoclonal sample in the simulated dataset by the sample mean scATAC-seq epithelial fraction for polyps $\bar{\rho}$. Using this strategy, we estimated the expected false polyclonal rate due to purity misestimation as 14.9%, which is lower than our estimated frequency of polyclonal polyps in the HTAN WGS data (40% of benign polyps, 28% of dysplastic polyps). This false positive rate represents an upper bound on the error of the estimated polyclonal polyp fraction, since it does not account for possible false negative monoclonal classifications, where truly polyclonal samples with higher purities than expected could be incorrectly classified as monoclonal.

We can also inspect the scATAC-seq measured epithelial cell fractions of samples that were profiled with both WGS and scATAC-seq. Focusing on the 13 polyp samples with overlap between these modalities (all dysplastic), we found that in the 9 samples we classified as monoclonal the scATAC-seq estimated purity generally provided a better approximation than the copy number based purity callers based on the position of the clonal peak, further supporting our approach using the scATAC-seq data to correct for purity in the ppVAFs (**Fig. SN2.2**).

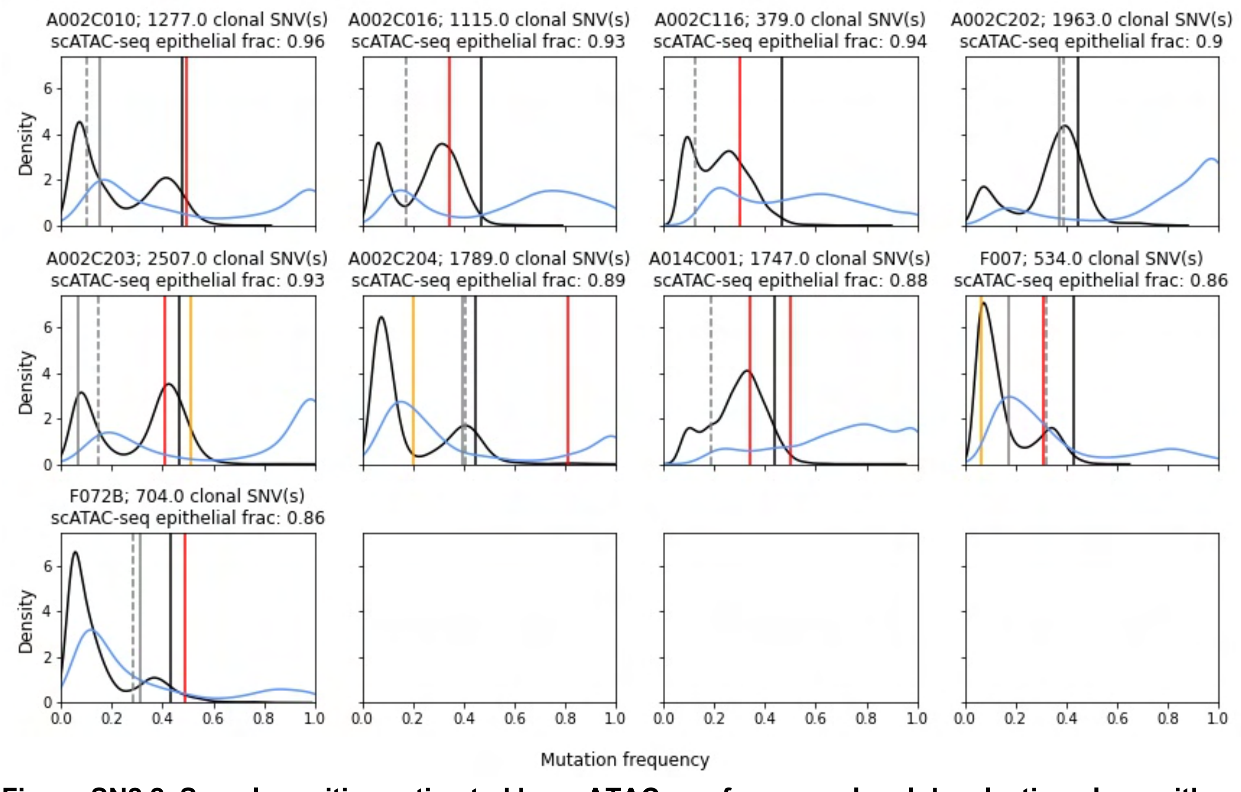

**Figure SN2.2. Sample purities estimated by scATAC-seq for monoclonal dysplastic polyps with matched WGS and scATAC-seq data.** The black distribution is raw VAFs and the blue curve is the ppVAF distribution. Red vertical lines are APC driver VAFs and yellow vertical lines are KRAS driver VAFs. Importantly, the black solid vertical lines are the expected heterozygous mutation VAFs given the matched sample scATAC-seq epithelial cell fraction. The grey vertical lines are the expected heterozygous mutation VAFs given the copy-number inferred purities (Sequenza: dashed, FACETS: solid).

Importantly, we also found that the 4 polyclonal dysplastic samples with data from both modalities do not have dramatically lower scATAC-seq estimated purity values than other samples (**Fig. SN2.3-4**, $p$=0.07, Wilcoxon rank-sum test). This suggests the inferred polyclonality in these 4 samples cannot be well explained by lower sample purity.

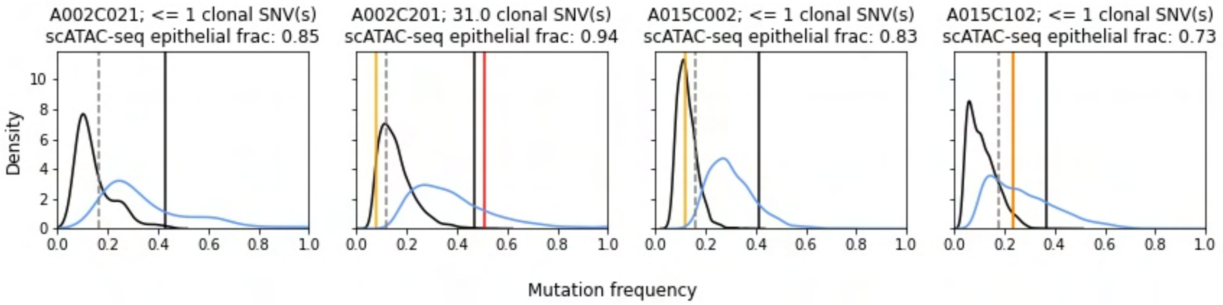

**Figure SN2.3. Sample purities estimated by scATAC-seq for polyclonal dysplastic polyps with matched WGS and scATAC-seq data. a.** The black distribution is raw VAFs and the blue curve is the ppVAF distribution. Red vertical lines are *APC* driver VAFs and yellow vertical lines are *KRAS* driver VAFs. Importantly, the black solid vertical lines are the expected heterozygous mutation VAFs given the matched sample scATAC-seq epithelial cell fraction. The grey vertical lines are the expected heterozygous mutation VAFs given the copy-number inferred purities (Sequenza: dashed, FACETS: solid).

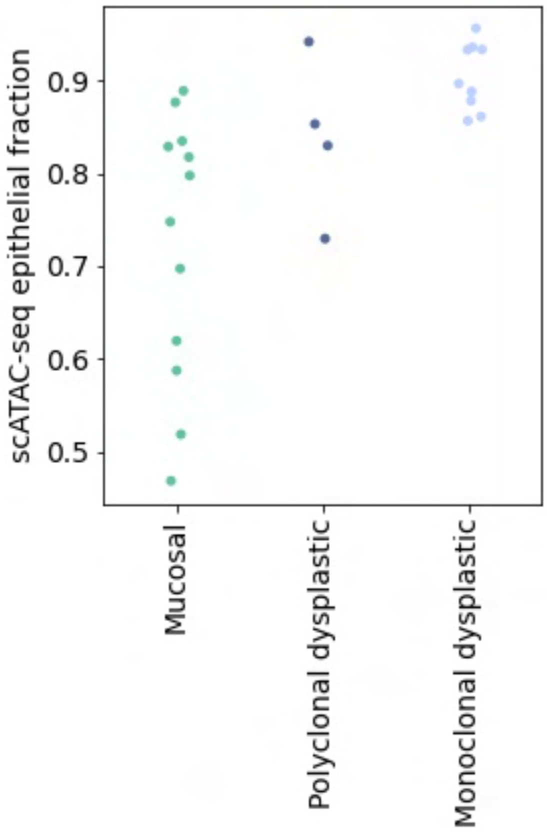

**Figure SN2.4. Comparison of scATAC-seq derived purities in polyclonal dysplastic, monoclonal dysplastic, and normal mucosal samples with matched single-cell data.**

### Supplementary Note 3: Single-gland whole-genome sequencing analysis

#### Contents

**1 Raw VAF Histograms** **34**

**2 Filtered VAF Histograms** **39**

**3 Raw Coverage Histograms** **44**

**4 Filtered Coverage Histograms** **49**

**5 Low-Coverage Driver Recovery Analysis** **54**

**6 Tree Inference Method Comparisons** **57**

### 1 Raw VAF Histograms

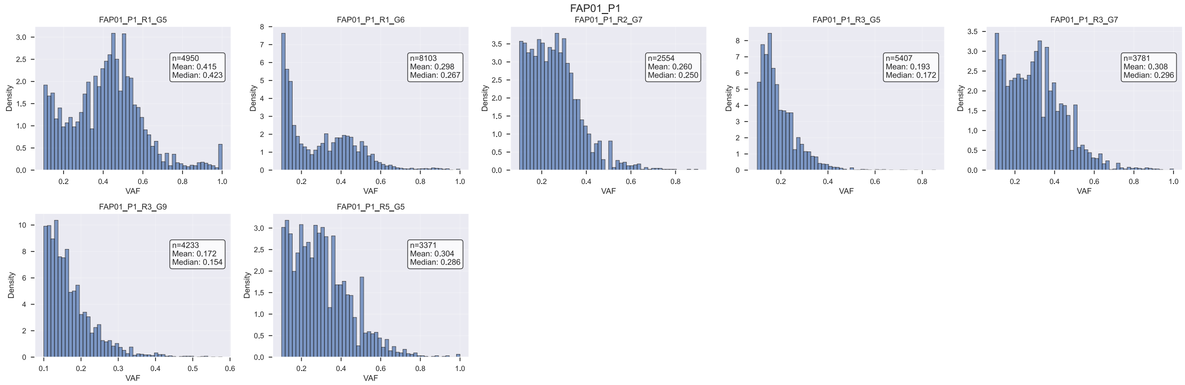

Figure SN3.1: Raw variant allele frequency (VAF) histograms for lesion FAP01_P1.

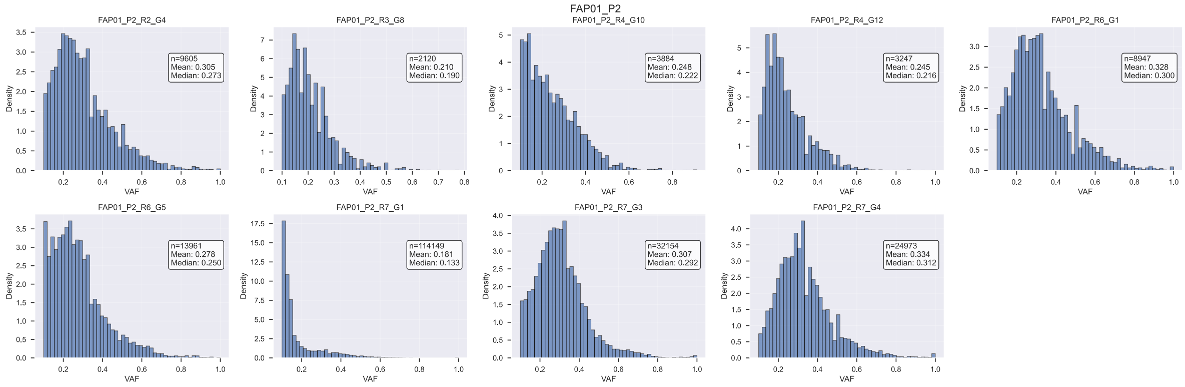

Figure SN3.2: Raw variant allele frequency (VAF) histograms for lesion FAP01_P2.

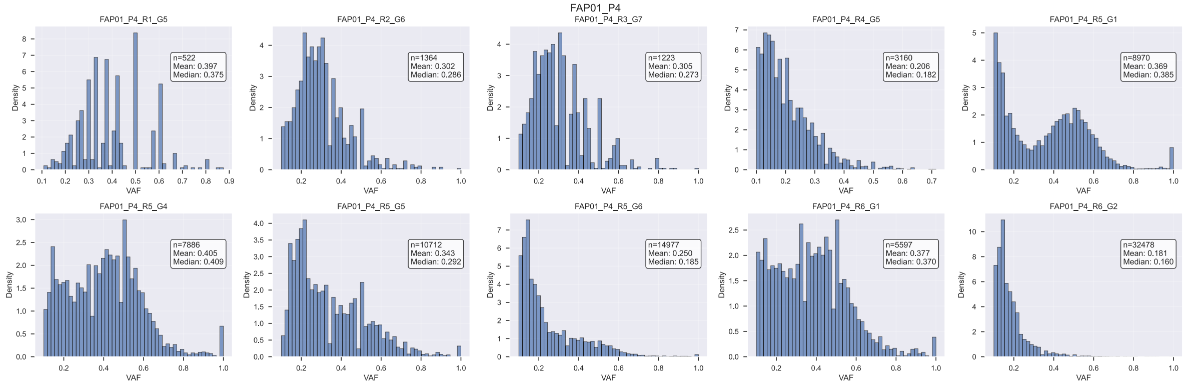

Figure SN3.3: Raw variant allele frequency (VAF) histograms for lesion FAP01_P4.

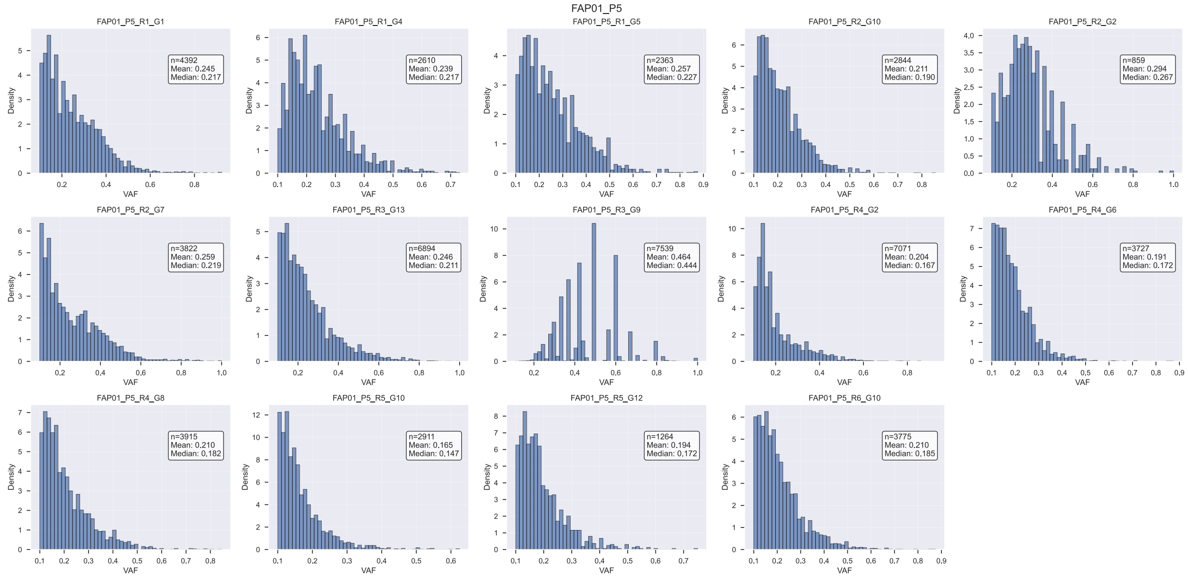

Figure SN3.4: Raw variant allele frequency (VAF) histograms for lesion FAP01_P5.

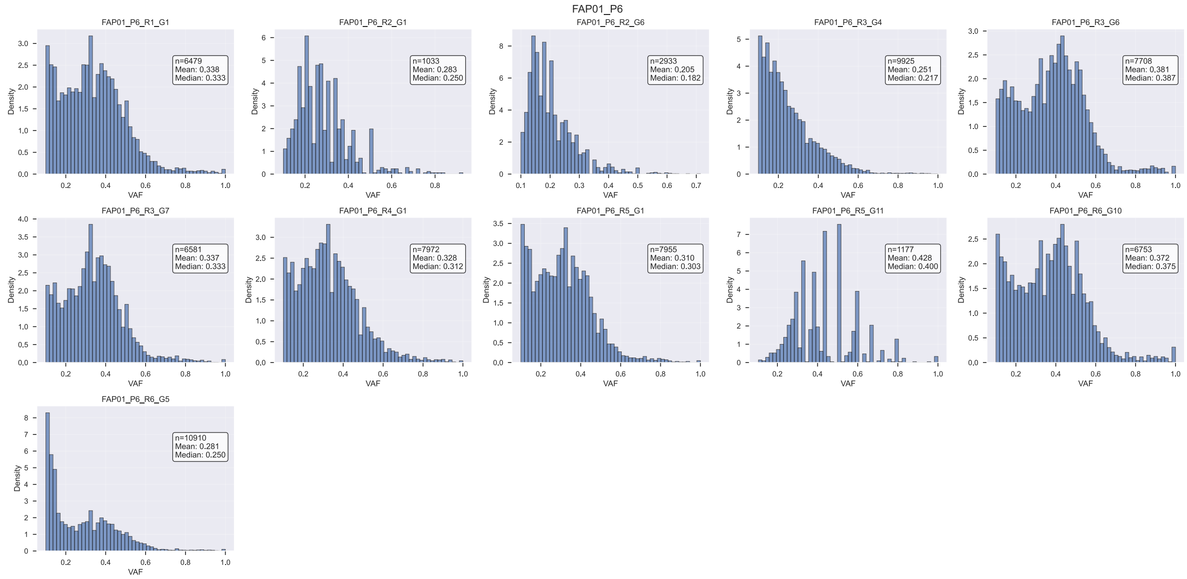

Figure SN3.5: Raw variant allele frequency (VAF) histograms for lesion FAP01_P6.

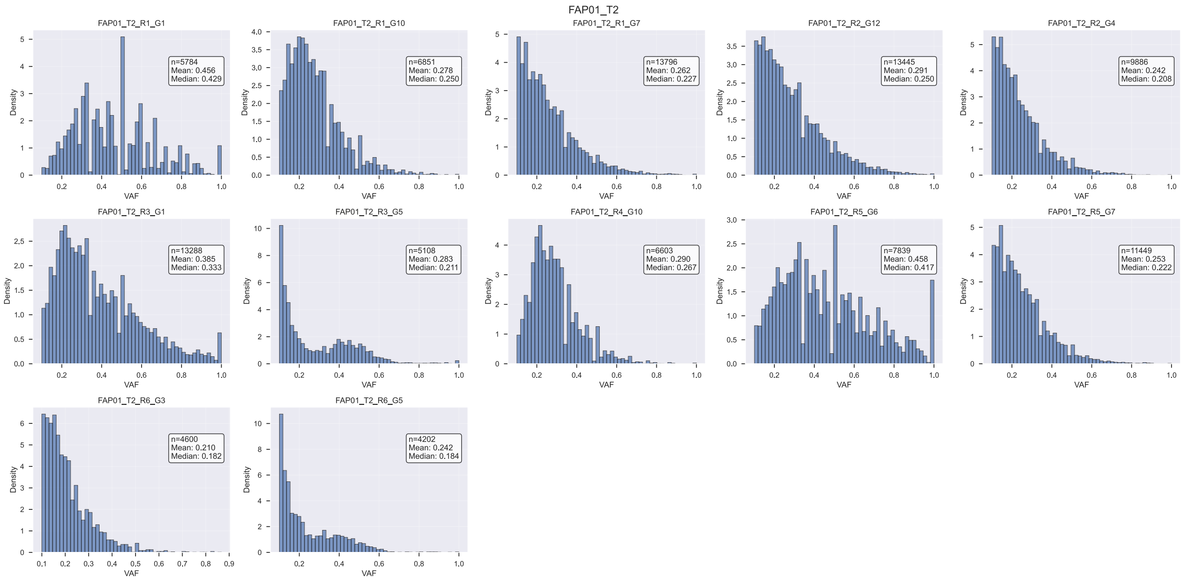

Figure SN3.6: Raw variant allele frequency (VAF) histograms for lesion FAP01_T2.

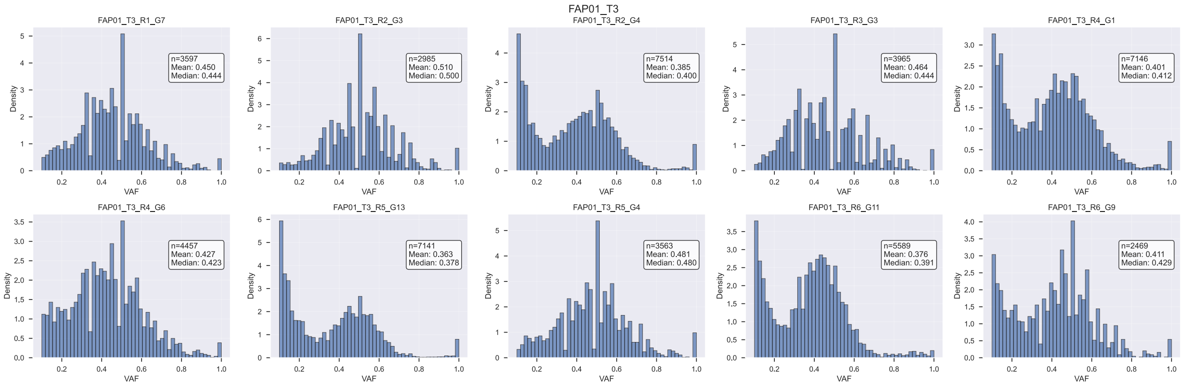

Figure SN3.7: Raw variant allele frequency (VAF) histograms for lesion FAP01_T3.

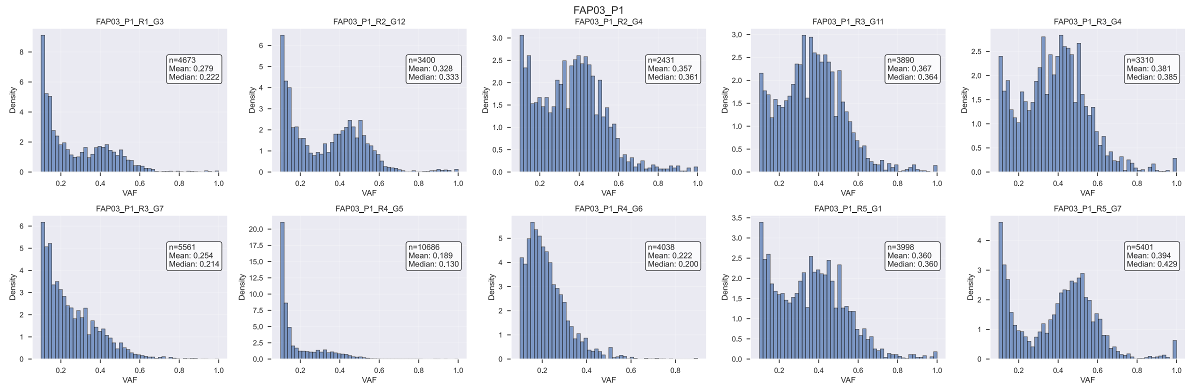

Figure SN3.8: Raw variant allele frequency (VAF) histograms for lesion FAP03_P1.

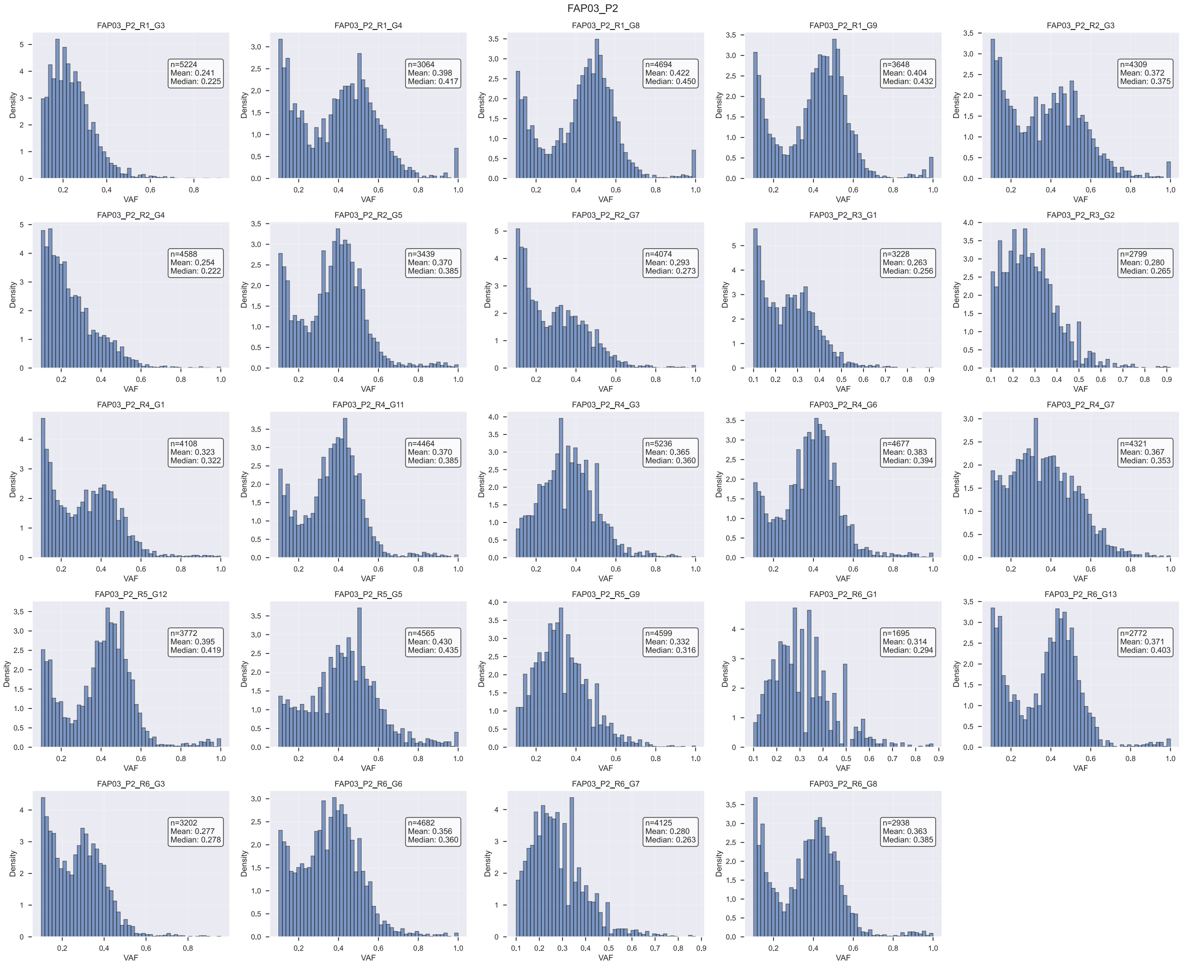

Figure SN3.9: Raw variant allele frequency (VAF) histograms for lesion FAP03_P2.

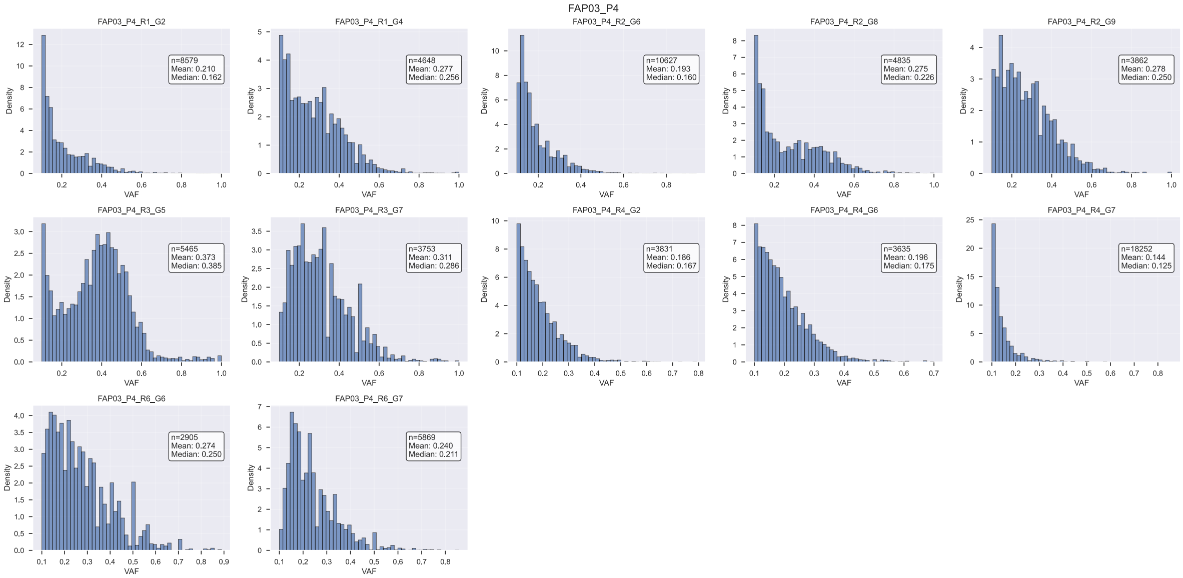

Figure SN3.10: Raw variant allele frequency (VAF) histograms for lesion FAP03_P4.

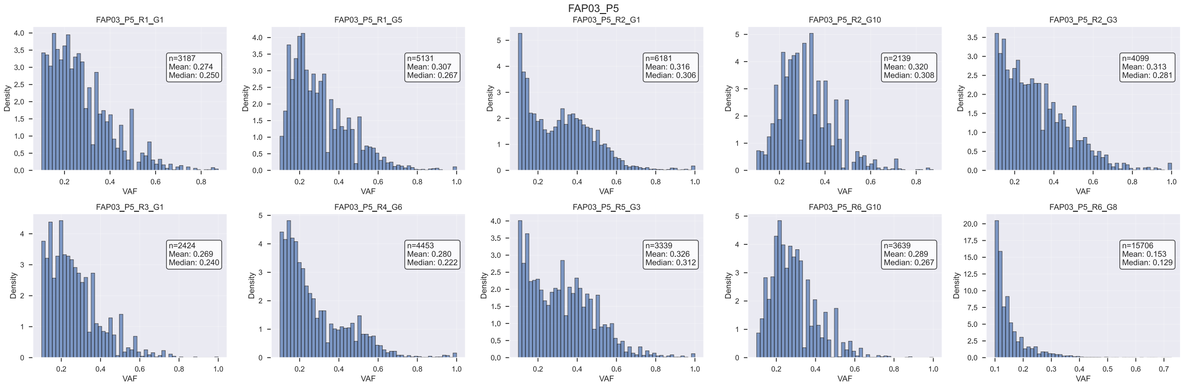

Figure SN3.11: Raw variant allele frequency (VAF) histograms for lesion FAP03_P5.

#### 2 Filtered VAF Histograms

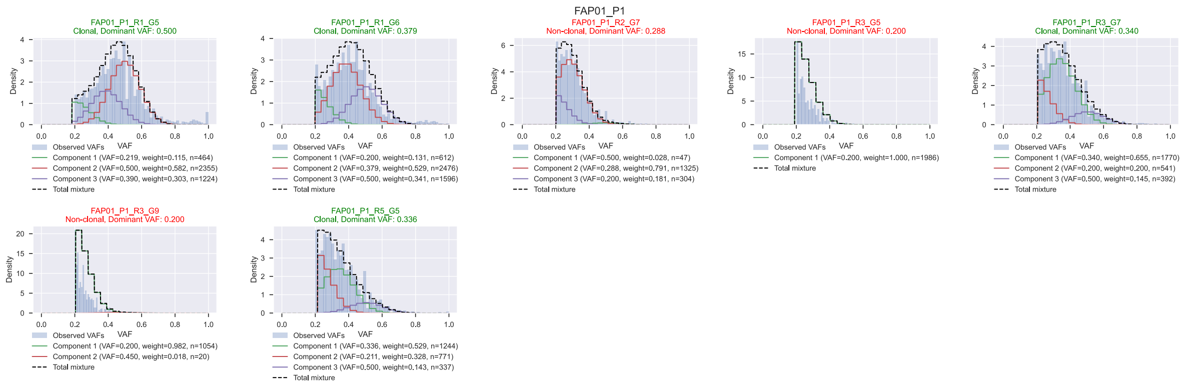

Figure SN3.12: Filtered variant allele frequency (VAF) histograms for lesion FAP01_P1. Also shown are the results of the truncated binomial model fit to the VAFs.

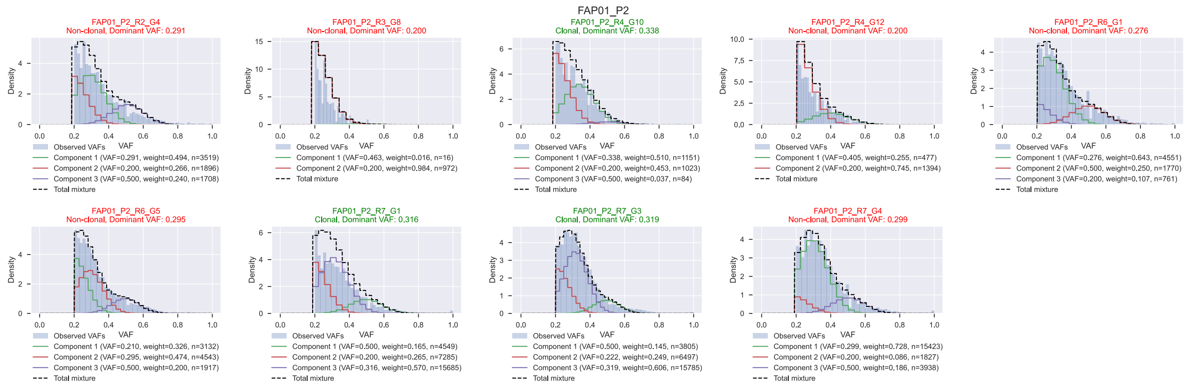

Figure SN3.13: Filtered variant allele frequency (VAF) histograms for lesion FAP01_P2. Also shown are the results of the truncated binomial model fit to the VAFs.

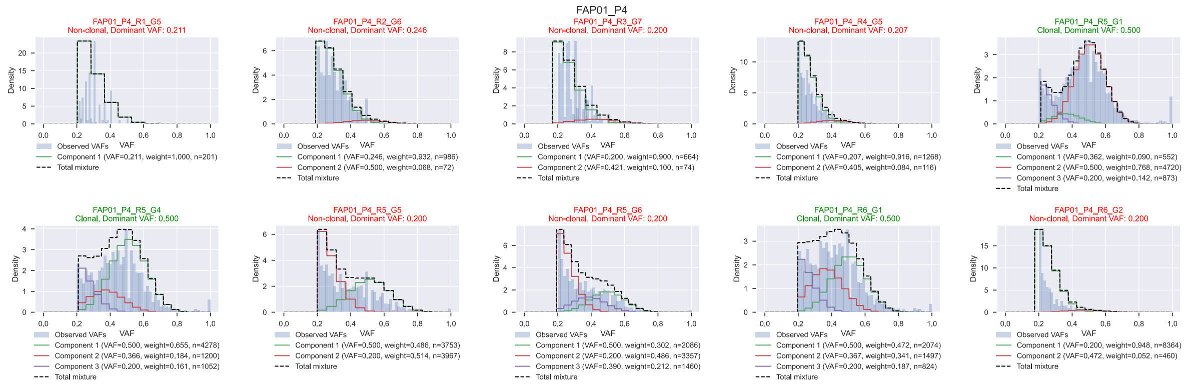

Figure SN3.14: Filtered variant allele frequency (VAF) histograms for lesion FAP01_P4. Also shown are the results of the truncated binomial model fit to the VAFs.

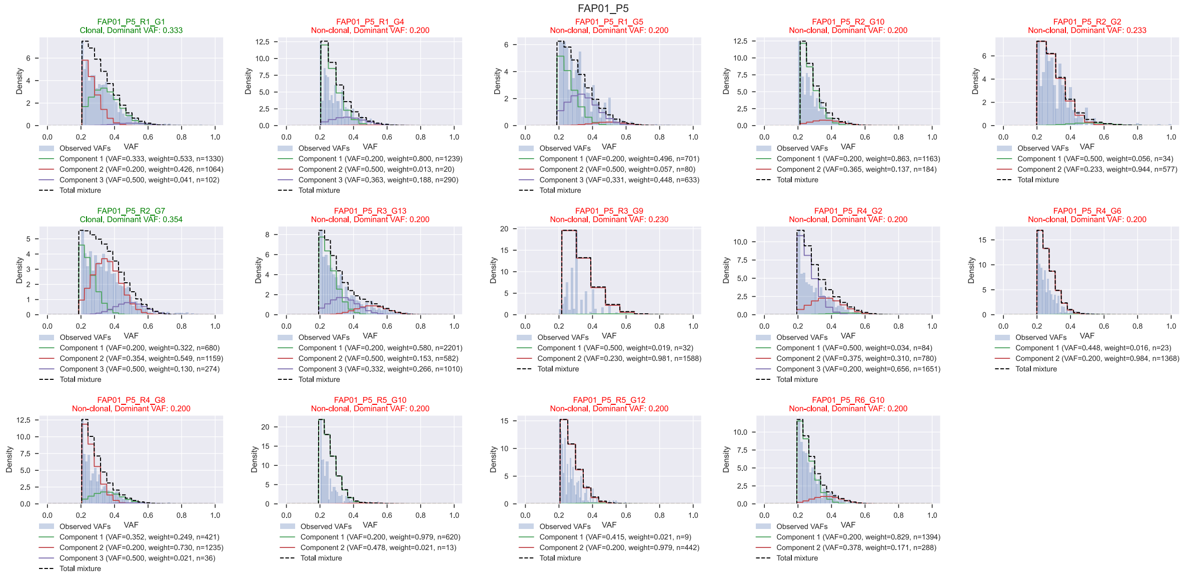

Figure SN3.15: Filtered variant allele frequency (VAF) histograms for lesion FAP01_P5. Also shown are the results of the truncated binomial model fit to the VAFs.

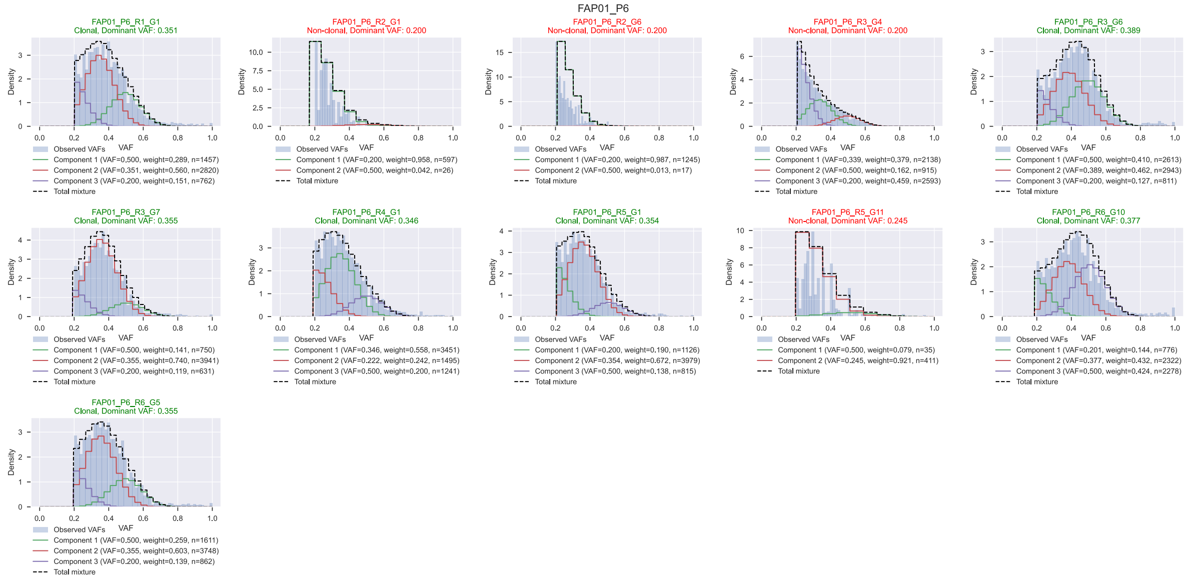

Figure SN3.16: Filtered variant allele frequency (VAF) histograms for lesion FAP01_P6. Also shown are the results of the truncated binomial model fit to the VAFs.

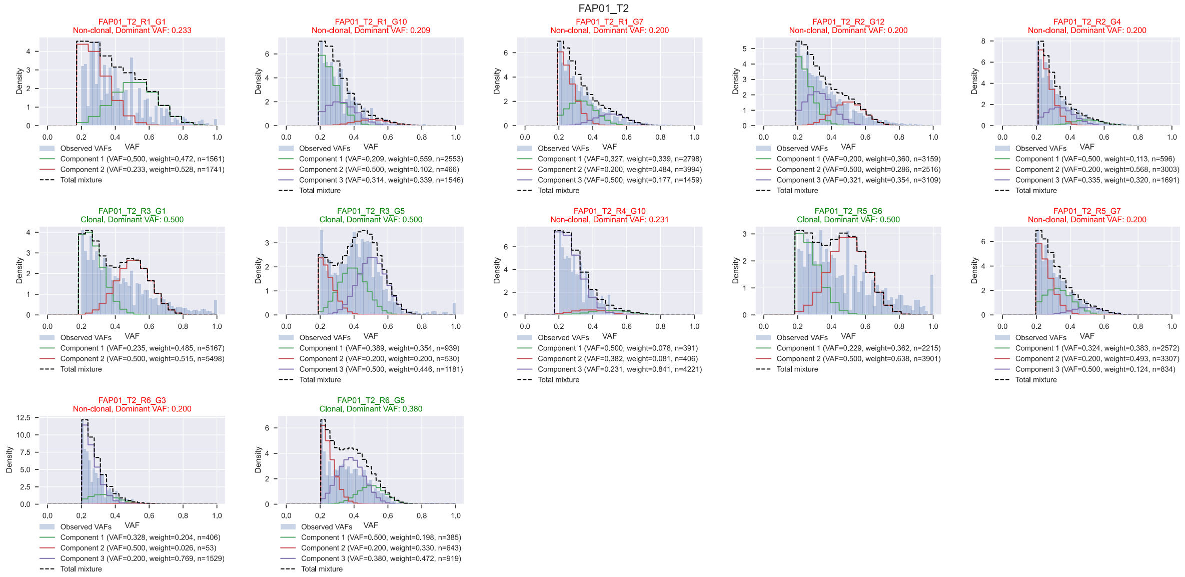

Figure SN3.17: Filtered variant allele frequency (VAF) histograms for lesion FAP01_T2. Also shown are the results of the truncated binomial model fit to the VAFs.

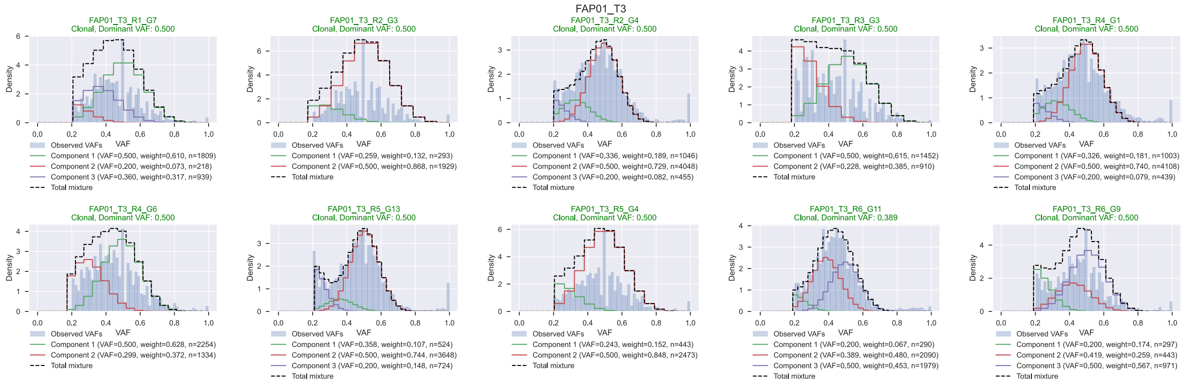

Figure SN3.18: Filtered variant allele frequency (VAF) histograms for lesion FAP01_T3. Also shown are the results of the truncated binomial model fit to the VAFs.

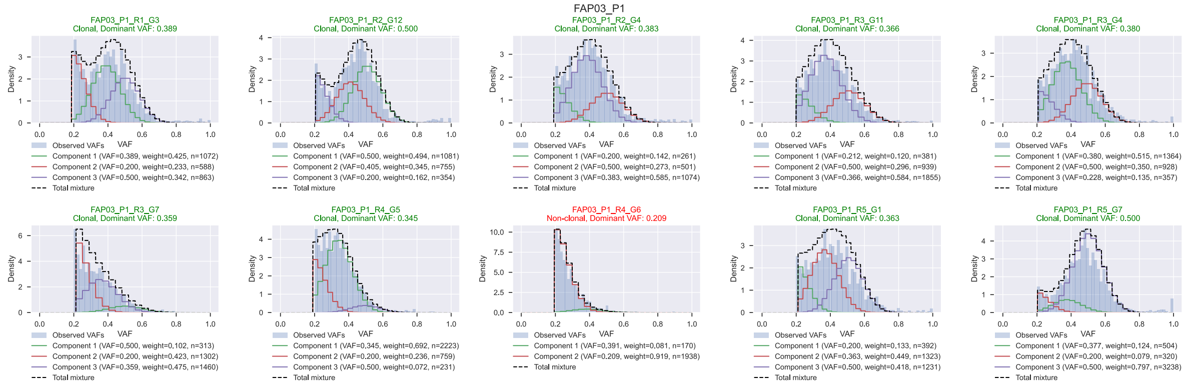

Figure SN3.19: Filtered variant allele frequency (VAF) histograms for lesion FAP03_P1. Also shown are the results of the truncated binomial model fit to the VAFs.

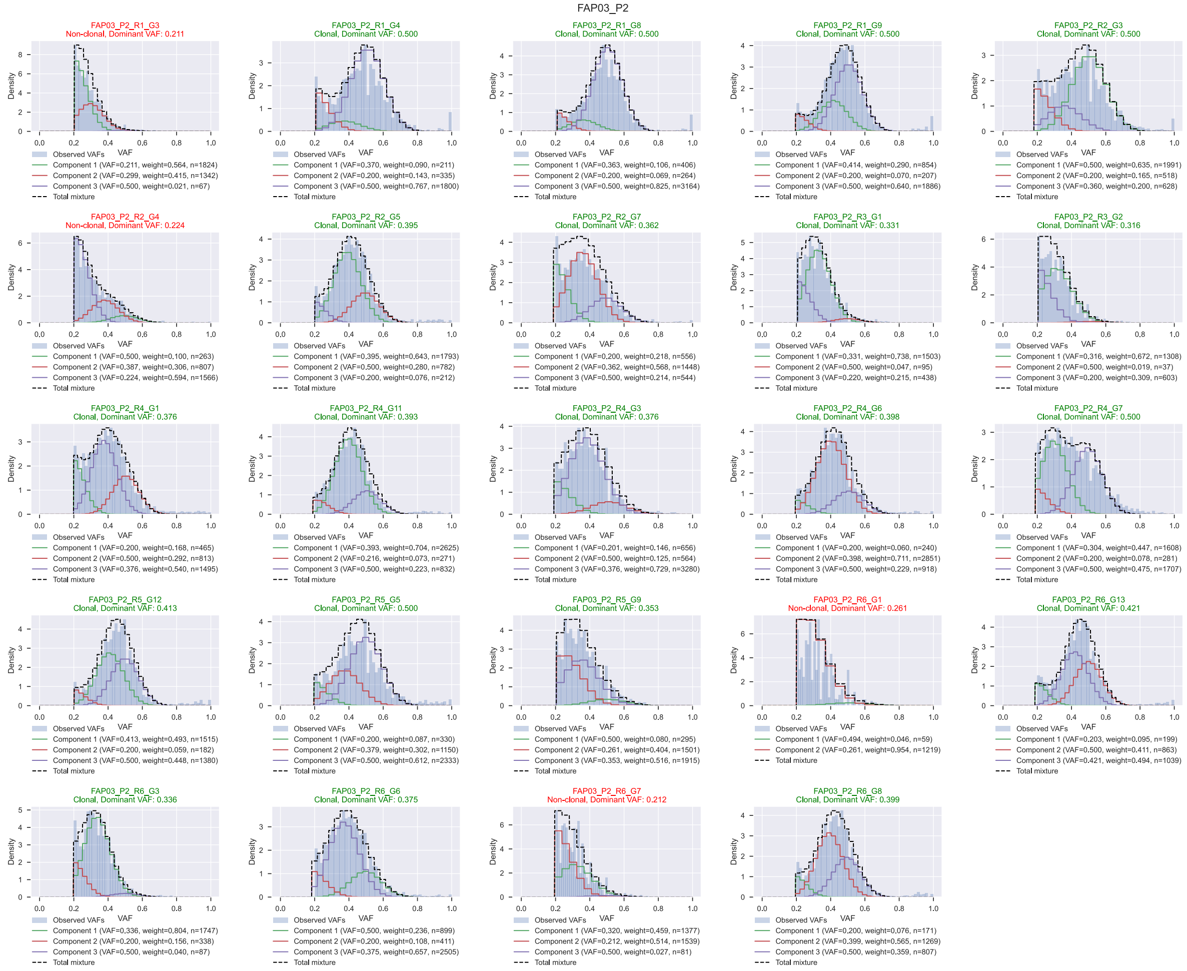

Figure SN3.20: Filtered variant allele frequency (VAF) histograms for lesion FAP03_P2. Also shown are the results of the truncated binomial model fit to the VAFs.

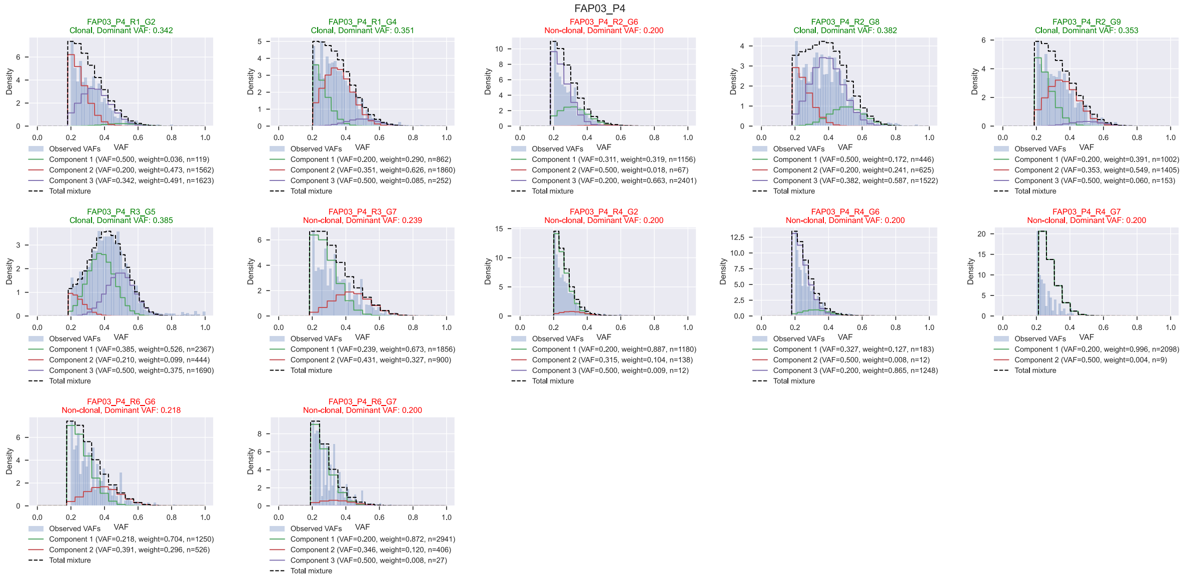

Figure SN3.21: Filtered variant allele frequency (VAF) histograms for lesion FAP03_P4. Also shown are the results of the truncated binomial model fit to the VAFs.

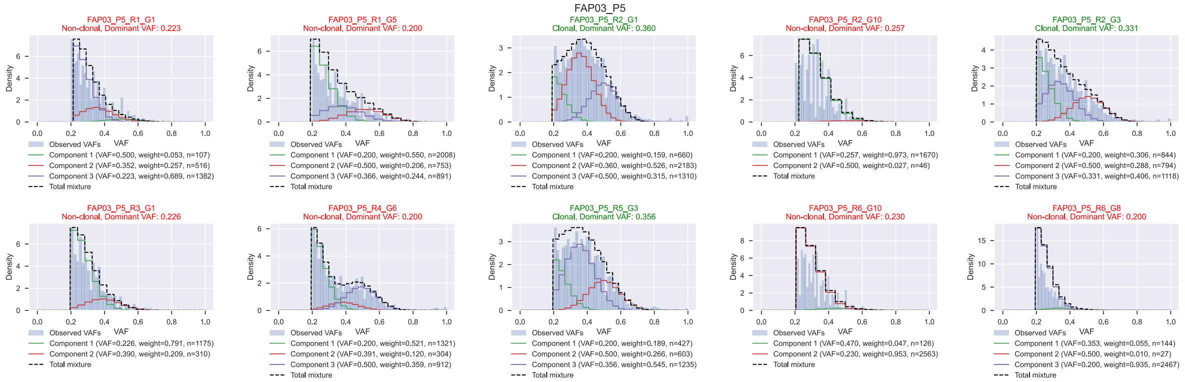

Figure SN3.22: Filtered variant allele frequency (VAF) histograms for lesion FAP03_P5. Also shown are the results of the truncated binomial model fit to the VAFs.

### 3 Raw Coverage Histograms

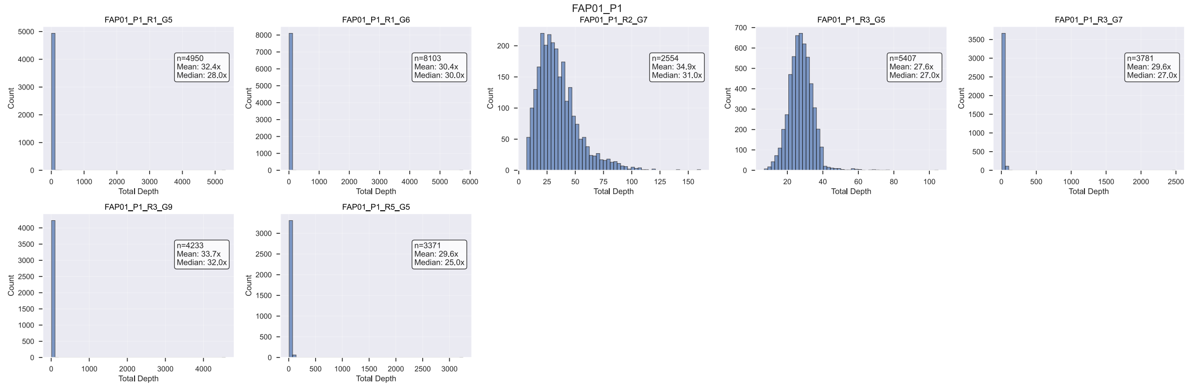

Figure SN3.23: Raw coverage histograms for lesion FAP01_P1.

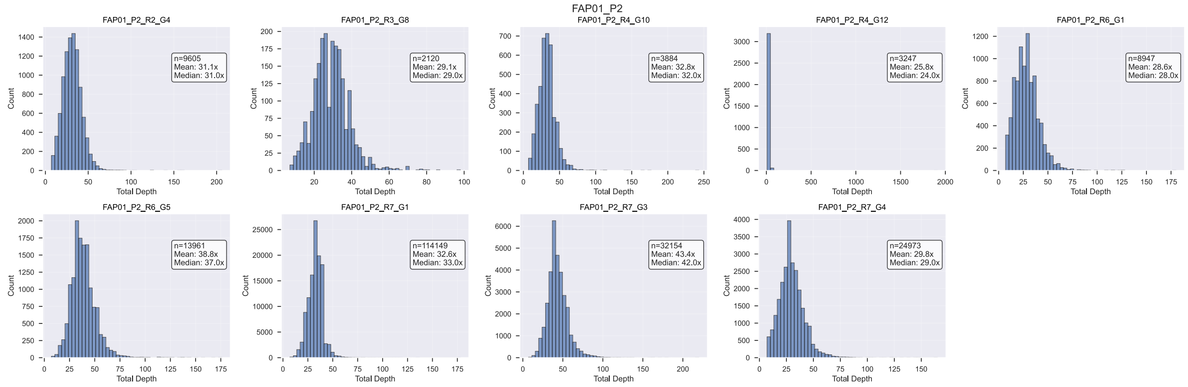

Figure SN3.24: Raw coverage histograms for lesion FAP01_P2.

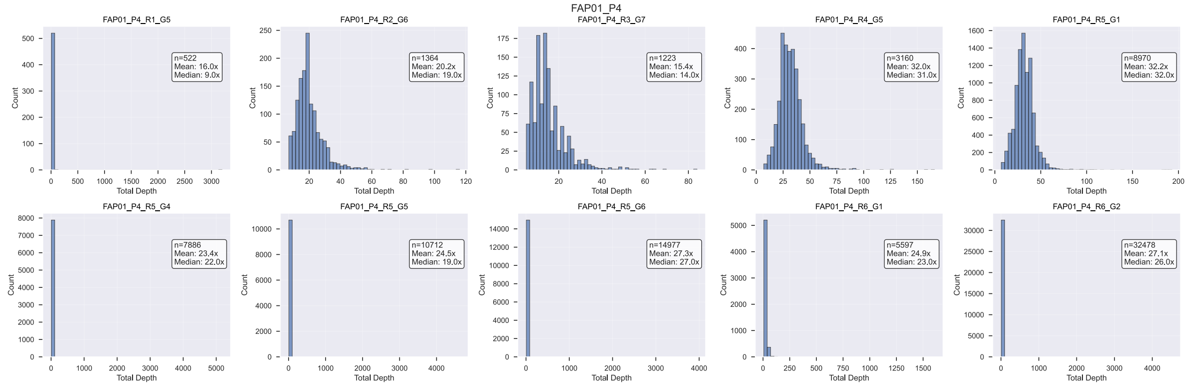

Figure SN3.25: Raw coverage histograms for lesion FAP01_P4.

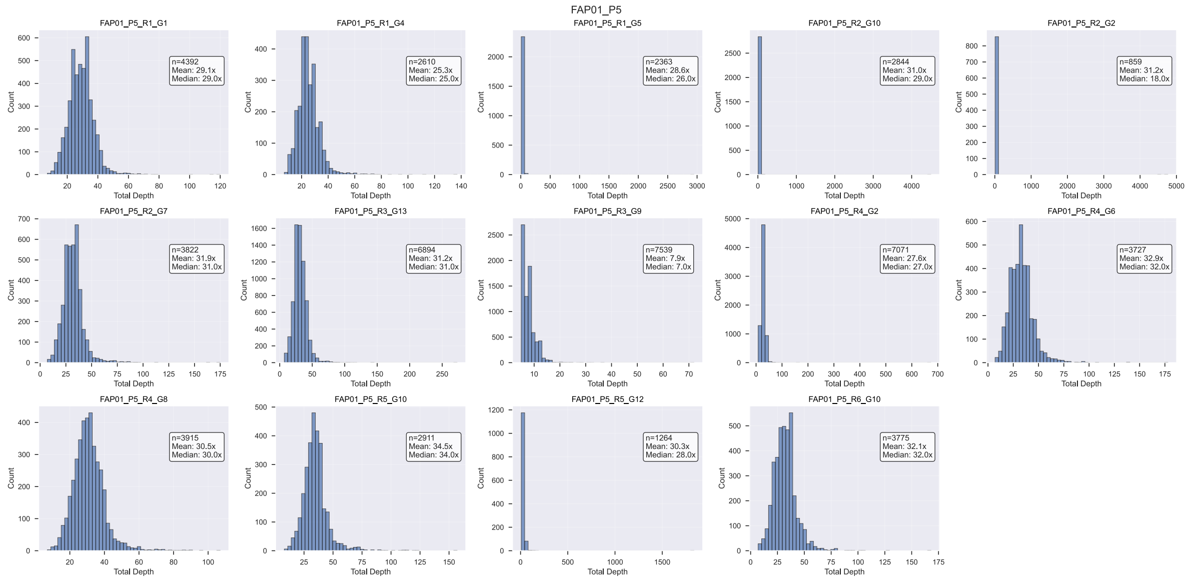

Figure SN3.26: Raw coverage histograms for lesion FAP01_P5.

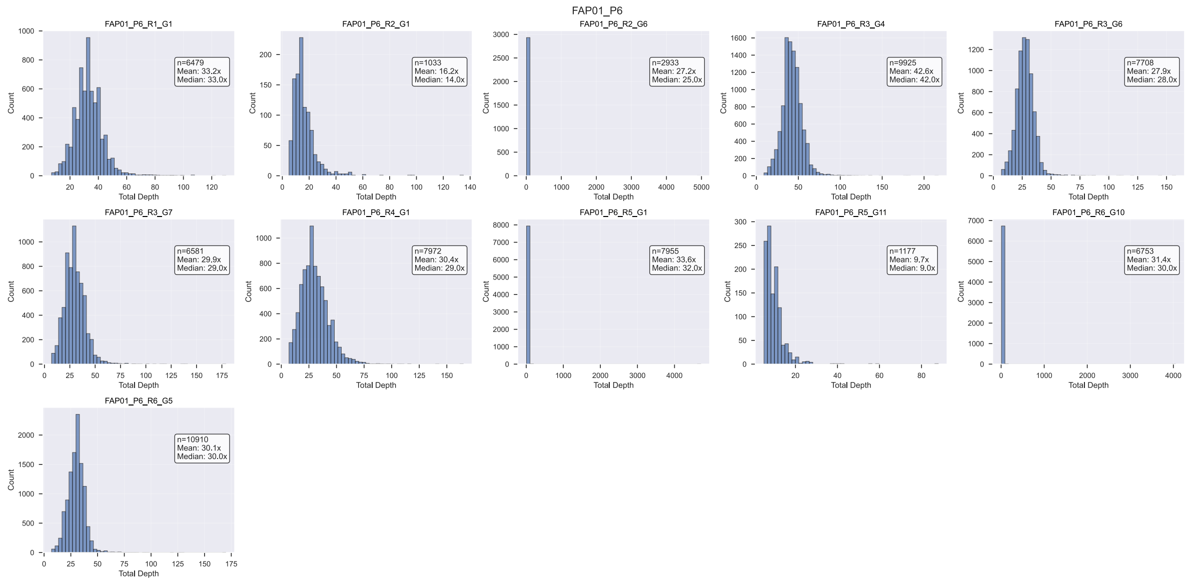

Figure SN3.27: Raw coverage histograms for lesion FAP01_P6.

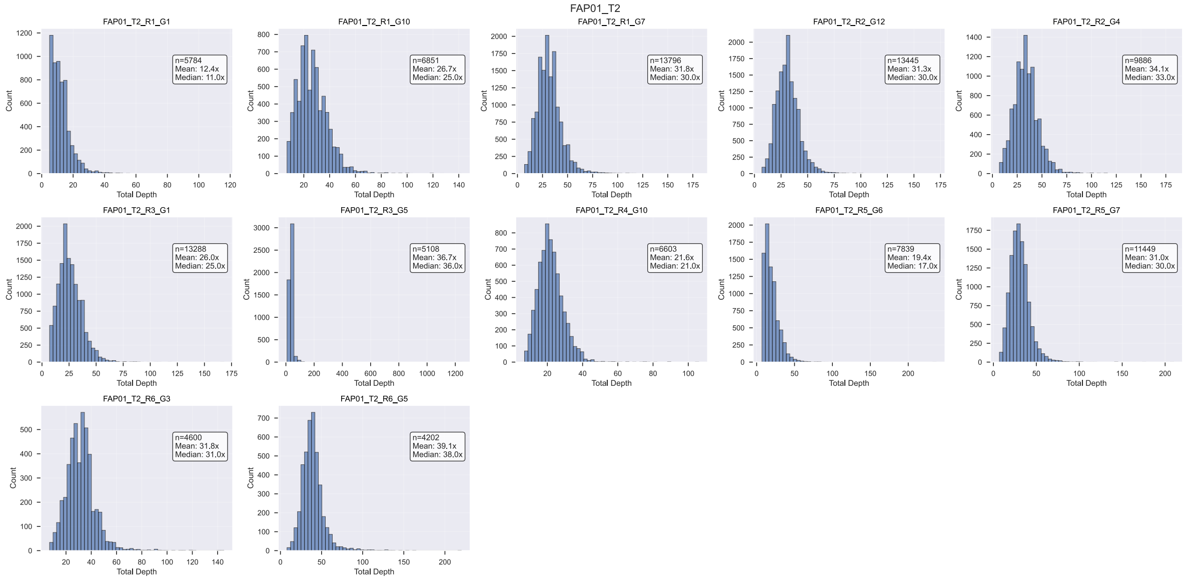

Figure SN3.28: Raw coverage histograms for lesion FAP01_T2.

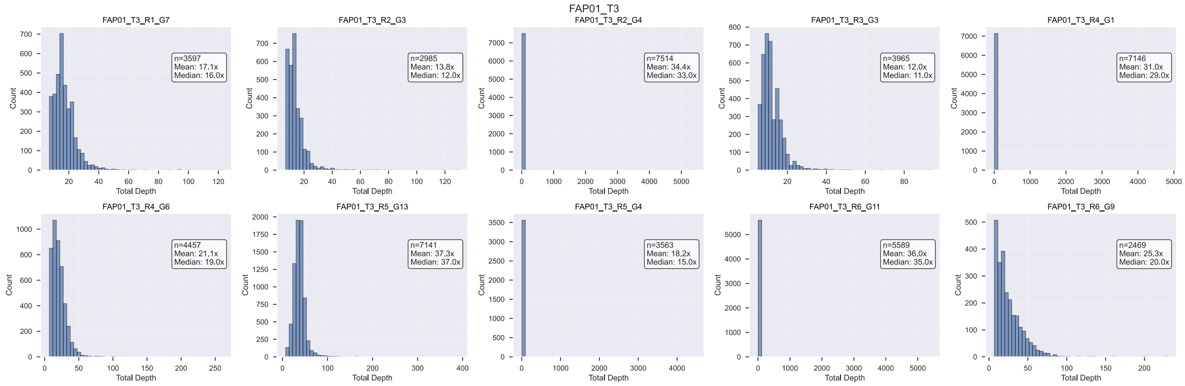

Figure SN3.29: Raw coverage histograms for lesion FAP01_T3.

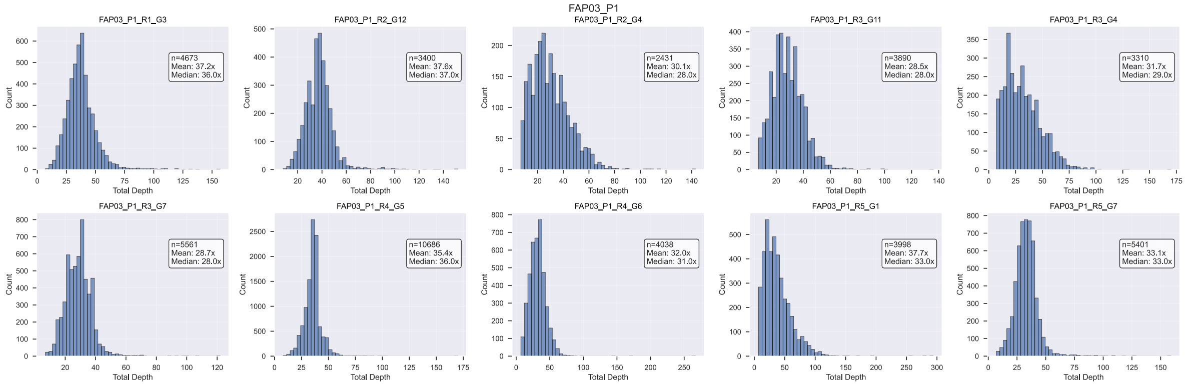

Figure SN3.30: Raw coverage histograms for lesion FAP03_P1.

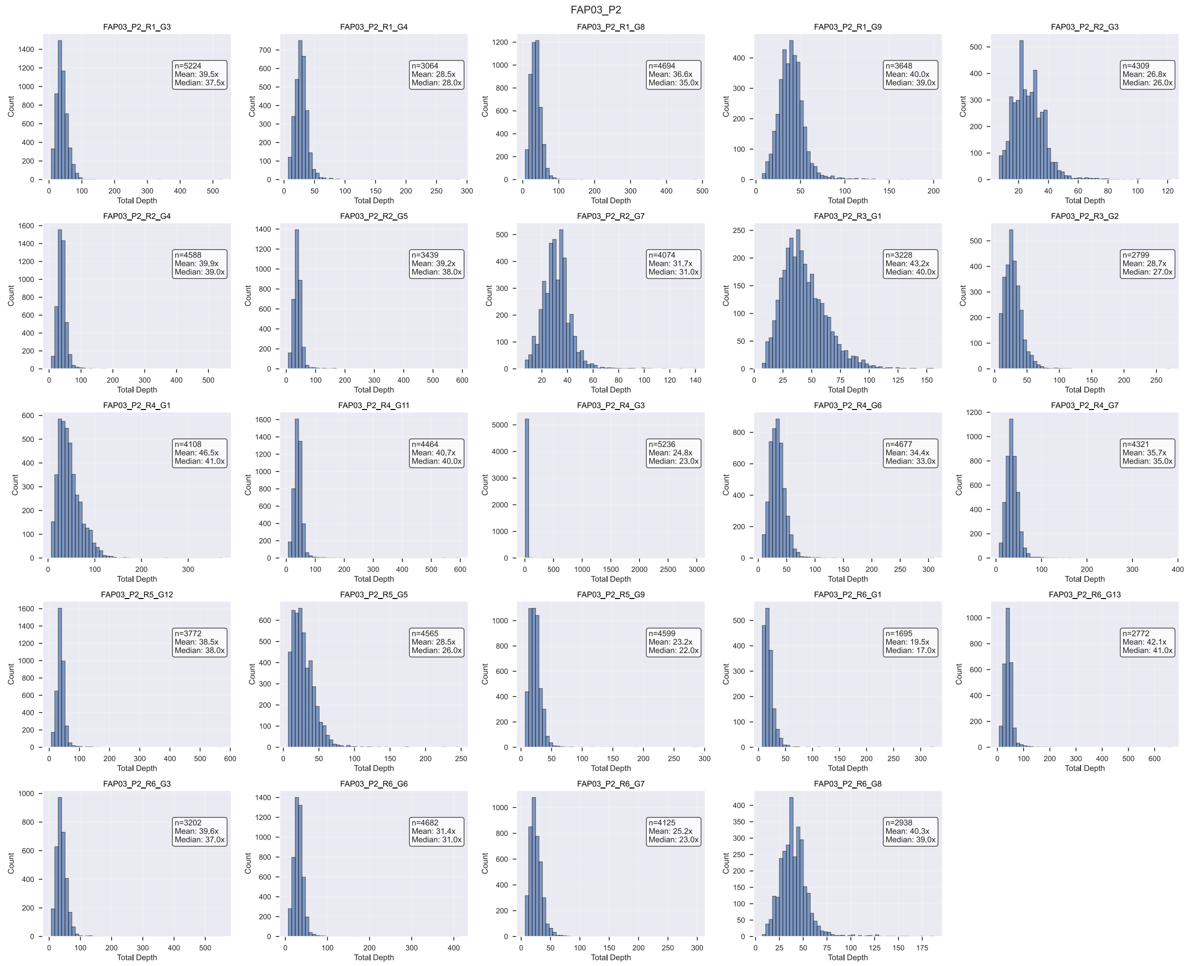

Figure SN3.31: Raw coverage histograms for lesion FAP03_P2.

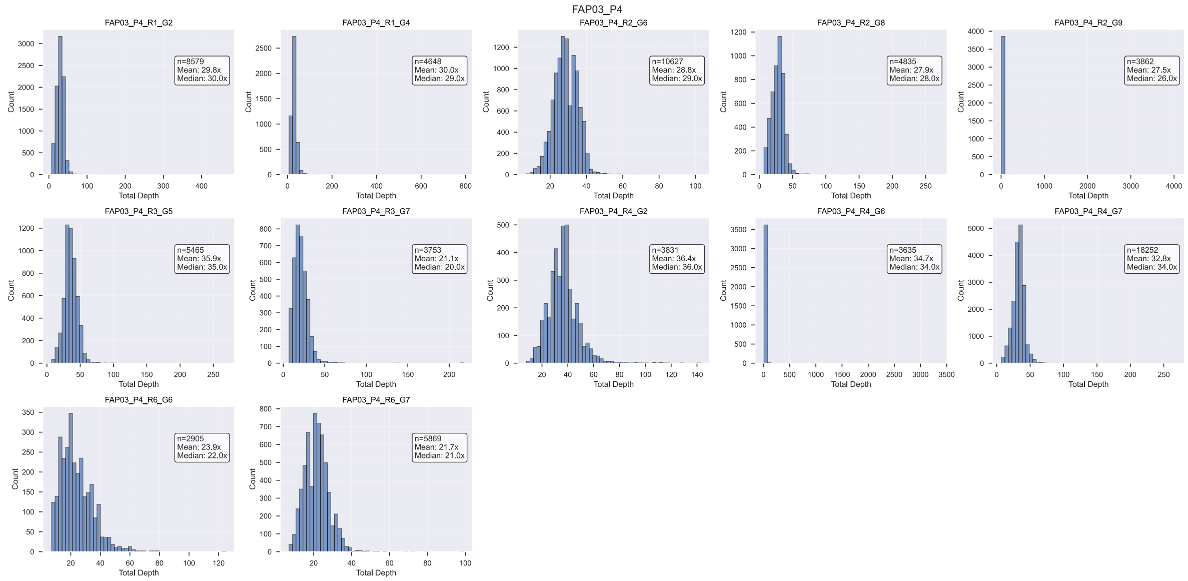

Figure SN3.32: Raw coverage histograms for lesion FAP03_P4.

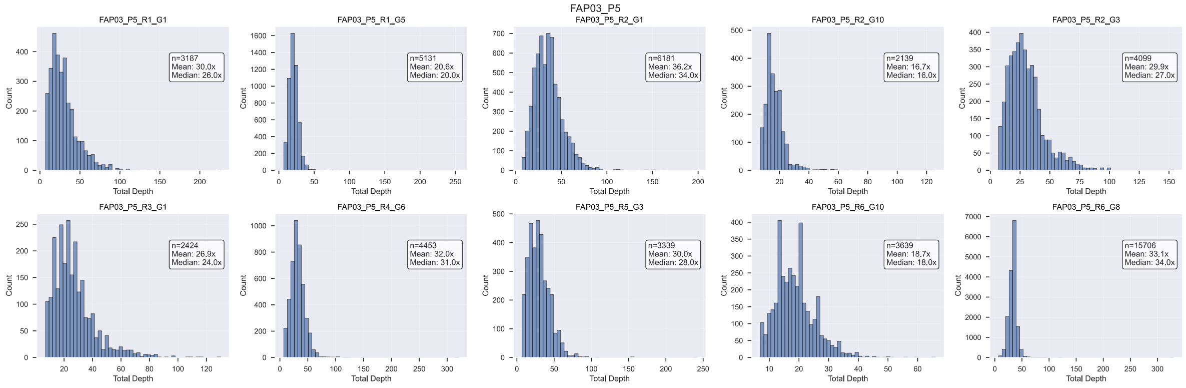

Figure SN3.33: Raw coverage histograms for lesion FAP03_P5.

### 4 Filtered Coverage Histograms

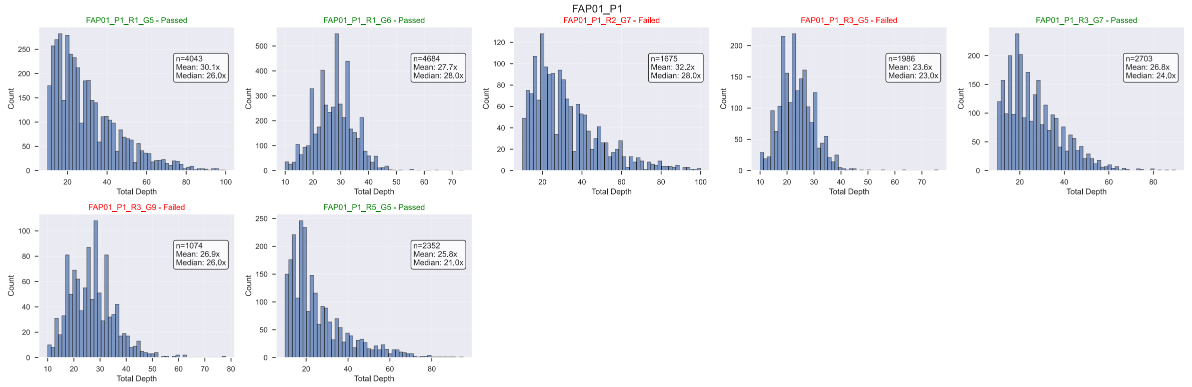

Figure SN3.34: Filtered coverage histograms for lesion FAP01_P1. Also shown in red/green are the results of the lesion-level filtering.

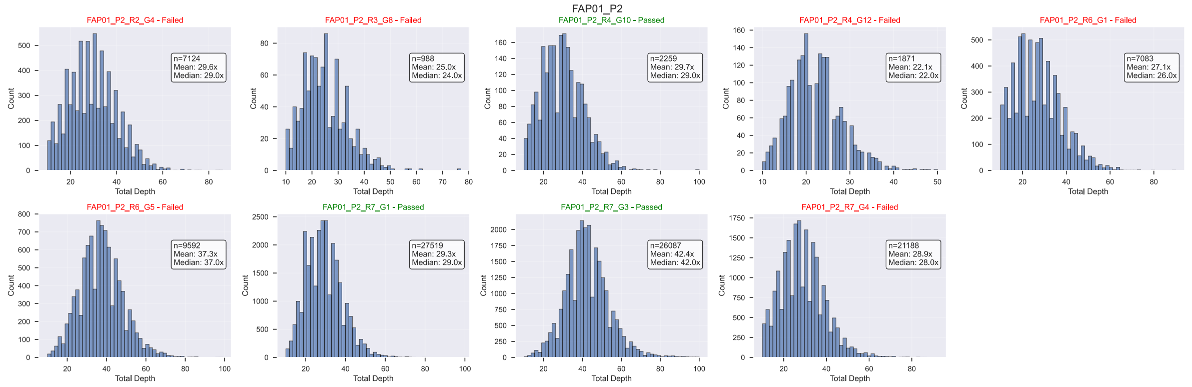

Figure SN3.35: Filtered coverage histograms for lesion FAP01_P2. Also shown in red/green are the results of the lesion-level filtering.

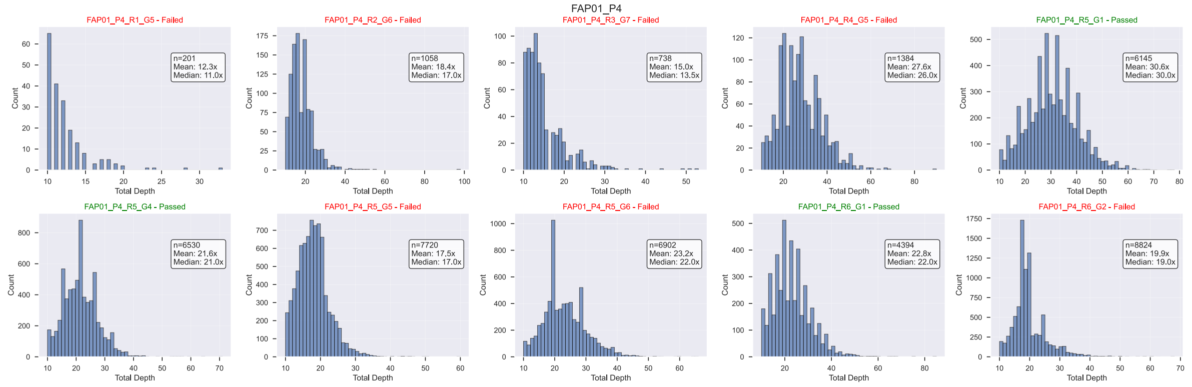

Figure SN3.36: Filtered coverage histograms for lesion FAP01_P4. Also shown in red/green are the results of the lesion-level filtering.

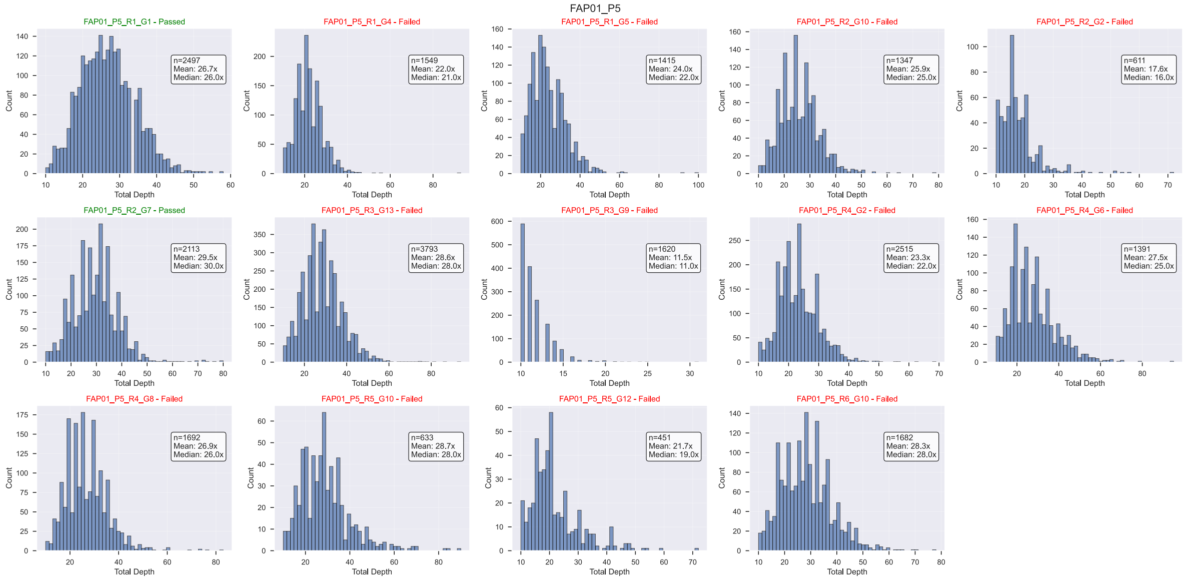

Figure SN3.37: Filtered coverage histograms for lesion FAP01_P5. Also shown in red/green are the results of the lesion-level filtering.

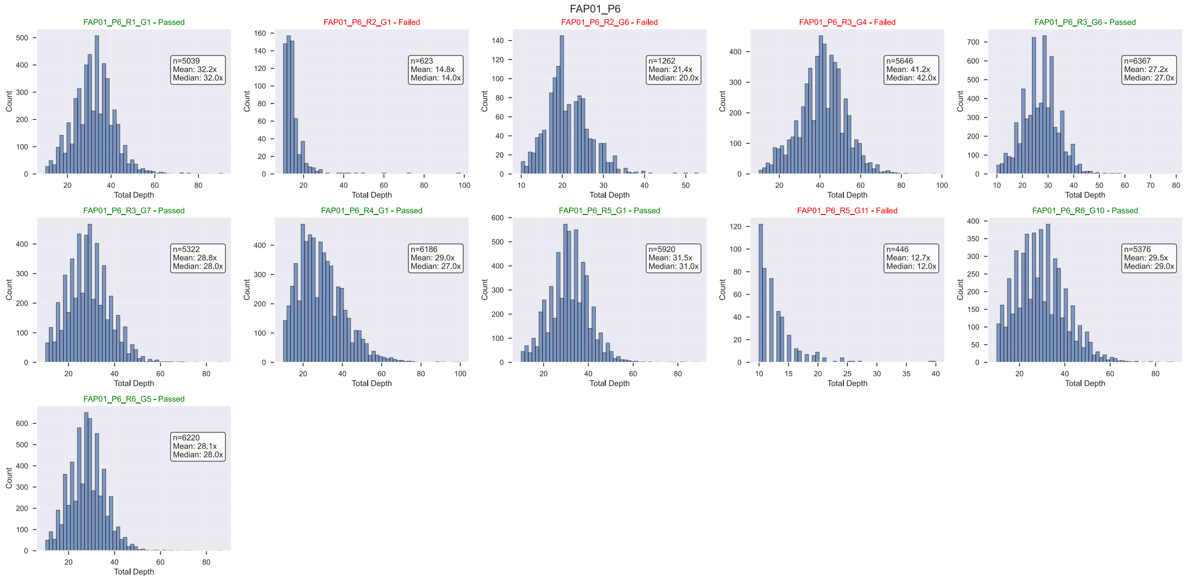

Figure SN3.38: Filtered coverage histograms for lesion FAP01_P6. Also shown in red/green are the results of the lesion-level filtering.

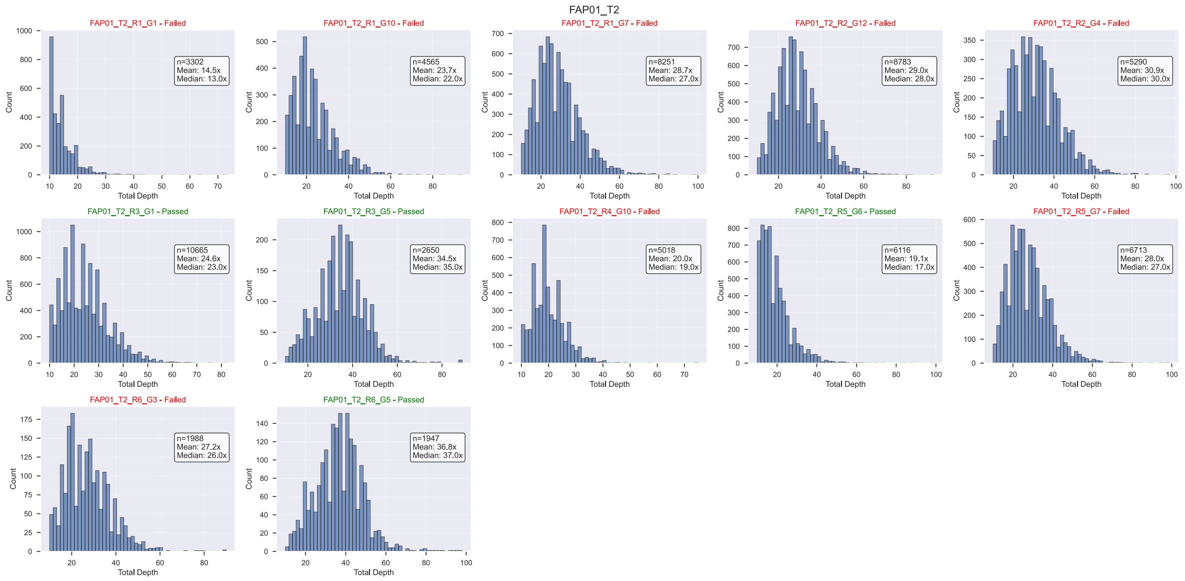

Figure SN3.39: Filtered coverage histograms for lesion FAP01_T2. Also shown in red/green are the results of the lesion-level filtering.

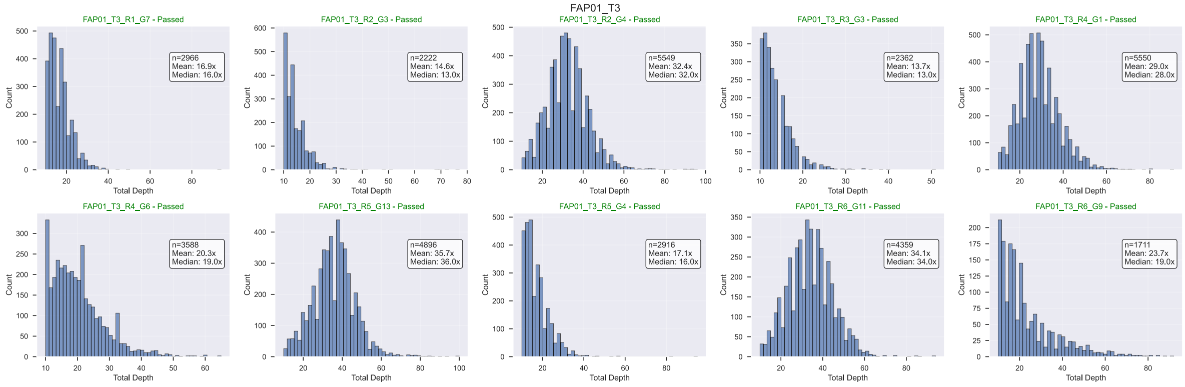

Figure SN3.40: Filtered coverage histograms for lesion FAP01_T3. Also shown in red/green are the results of the lesion-level filtering.

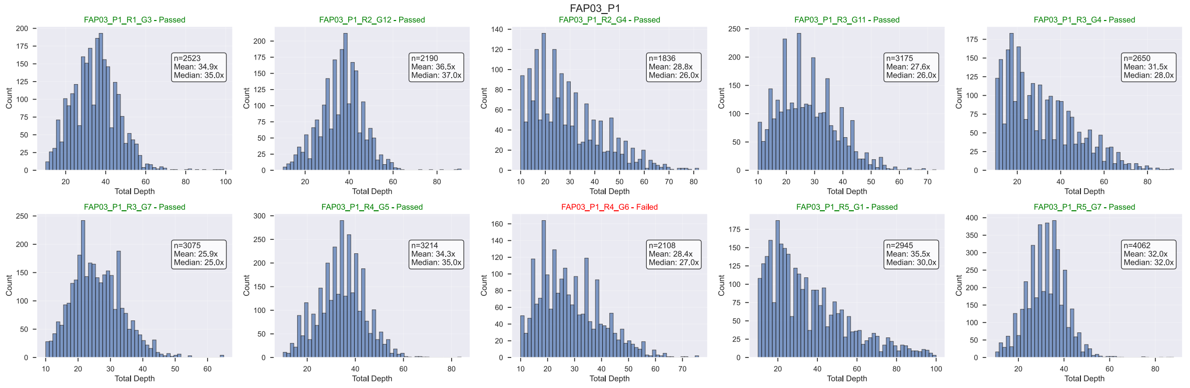

Figure SN3.41: Filtered coverage histograms for lesion FAP03_P1. Also shown in red/green are the results of the lesion-level filtering.

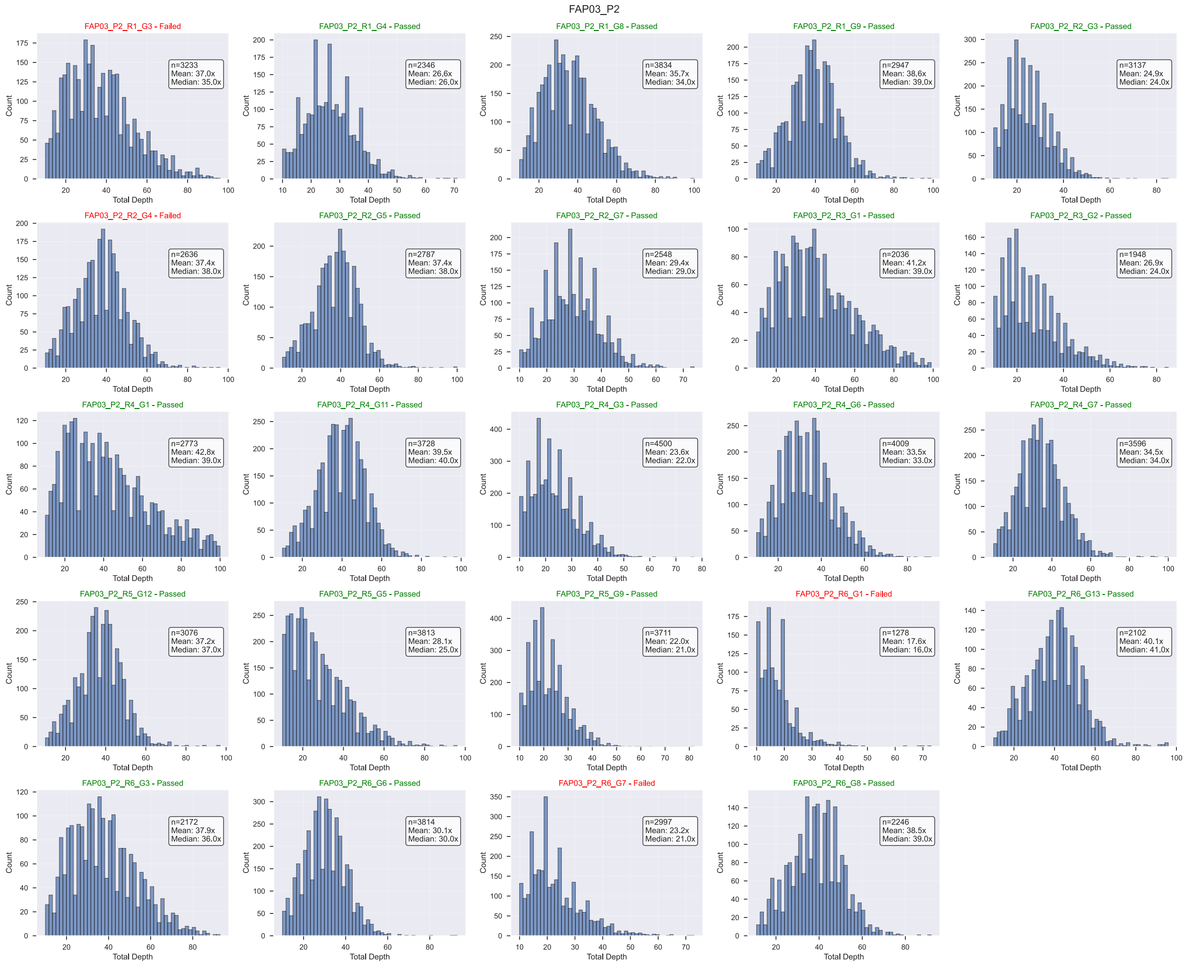

Figure SN3.42: Filtered coverage histograms for lesion FAP03_P2. Also shown in red/green are the results of the lesion-level filtering.

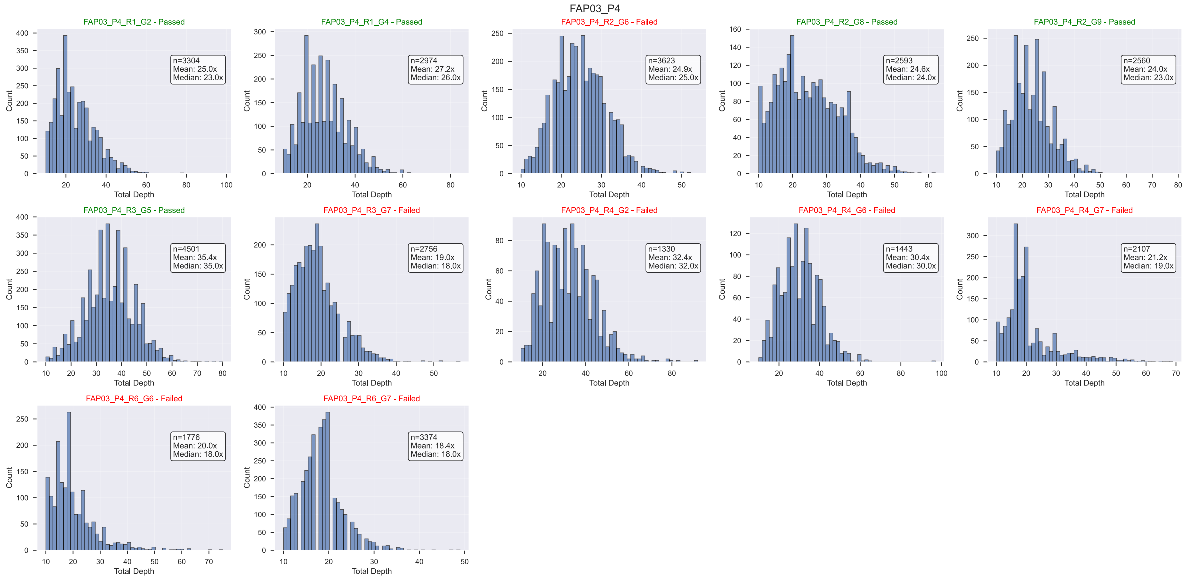

Figure SN3.43: Filtered coverage histograms for lesion FAP03_P4. Also shown in red/green are the results of the lesion-level filtering.

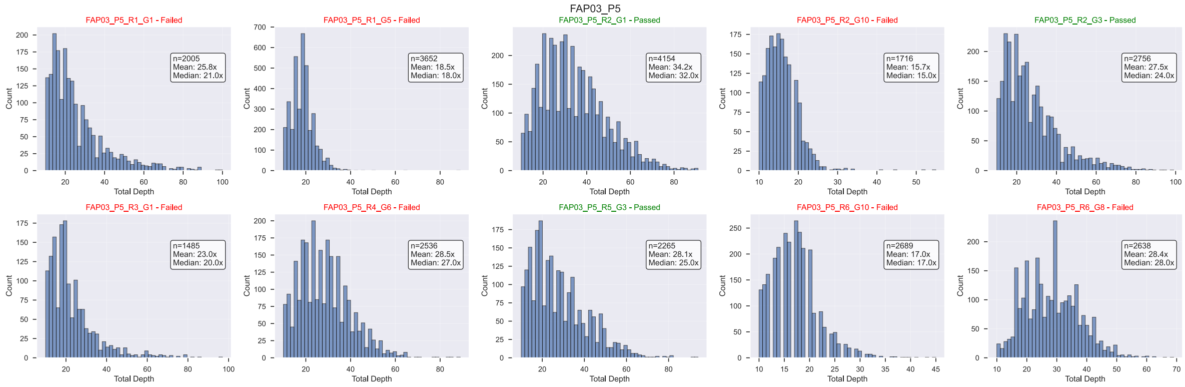

Figure SN3.44: Filtered coverage histograms for lesion FAP03_P5. Also shown in red/green are the results of the lesion-level filtering.

### 5 Low-Coverage Driver Recovery Analysis

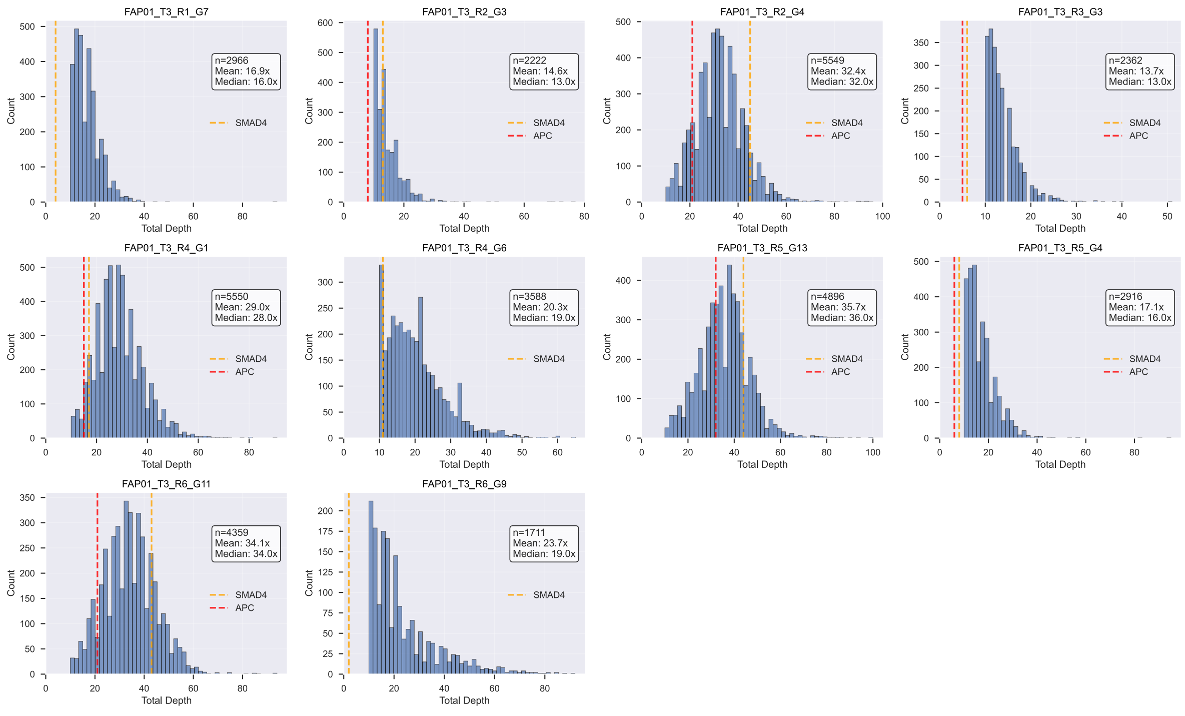

Figure SN3.45: Filtered coverage histograms for lesion FAP01_T3. Red dashed lines indicate coverage for APC mutations and orange dashed lines indicate coverage for SMAD4 mutations, as found either in the filtered dataset or recovered from the data initially filtered out.

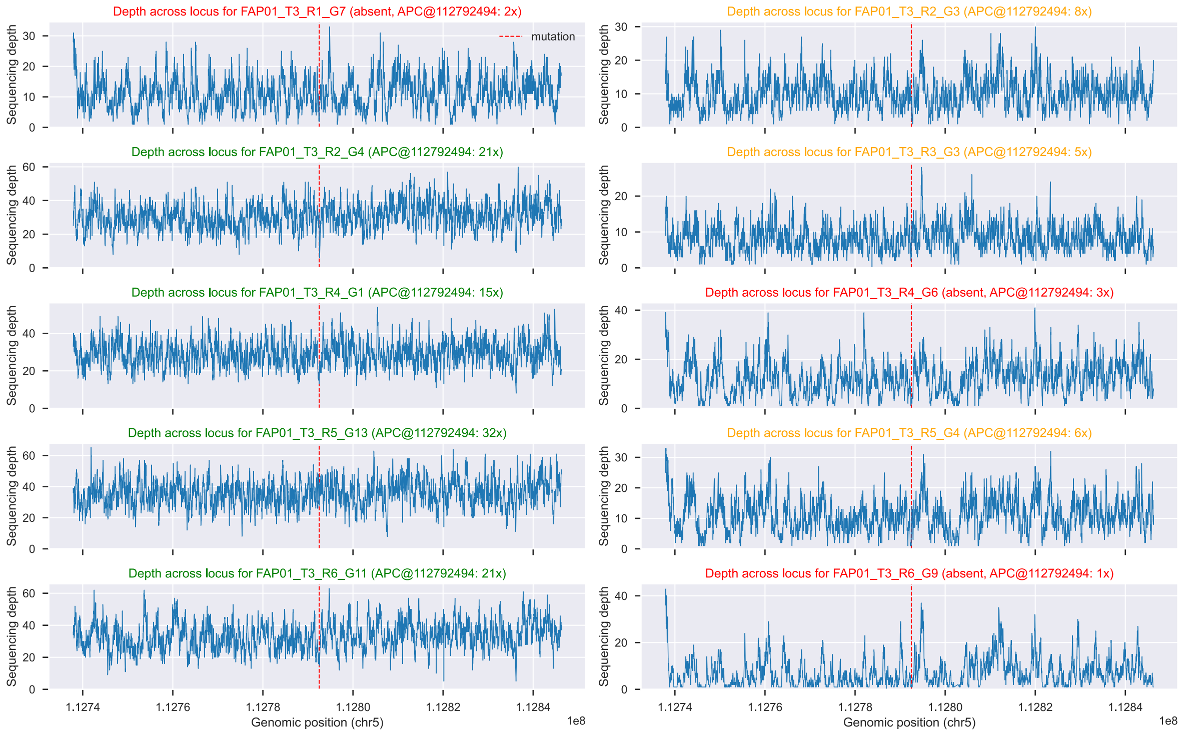

Figure SN3.46: Raw sequencing depth at the APC locus for samples from lesion FAP01_T3, with the APC mutation location highlighted. Green titles indicate samples in which the APC mutations passed our filters, orange titles indicate samples in which the APC mutations were initially filtered out, and red titles indicate samples in which the APC mutations were never detected, most likely due to low coverage.

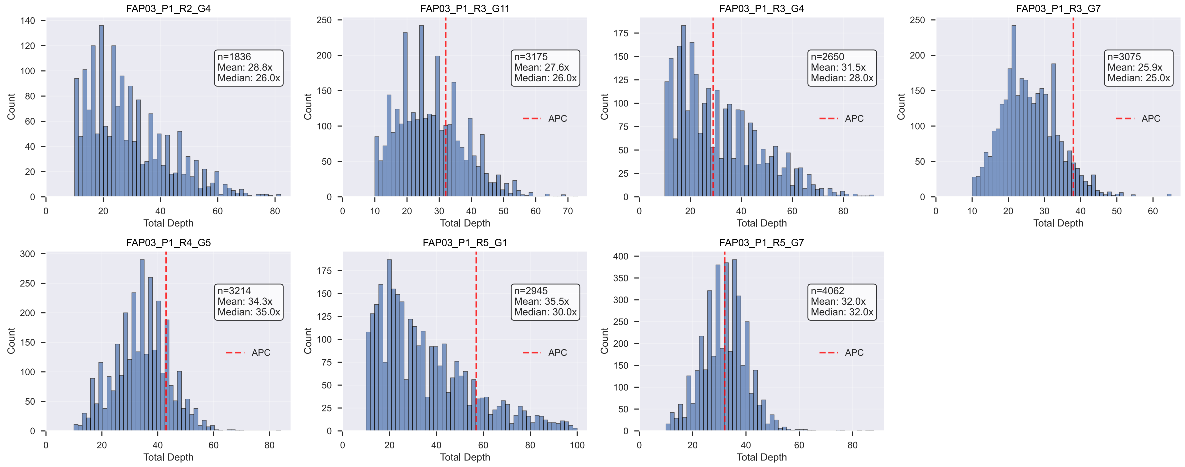

Figure SN3.47: Filtered coverage histograms for lesion FAP03_P1. Red dashed lines indicate coverage for APC mutations.

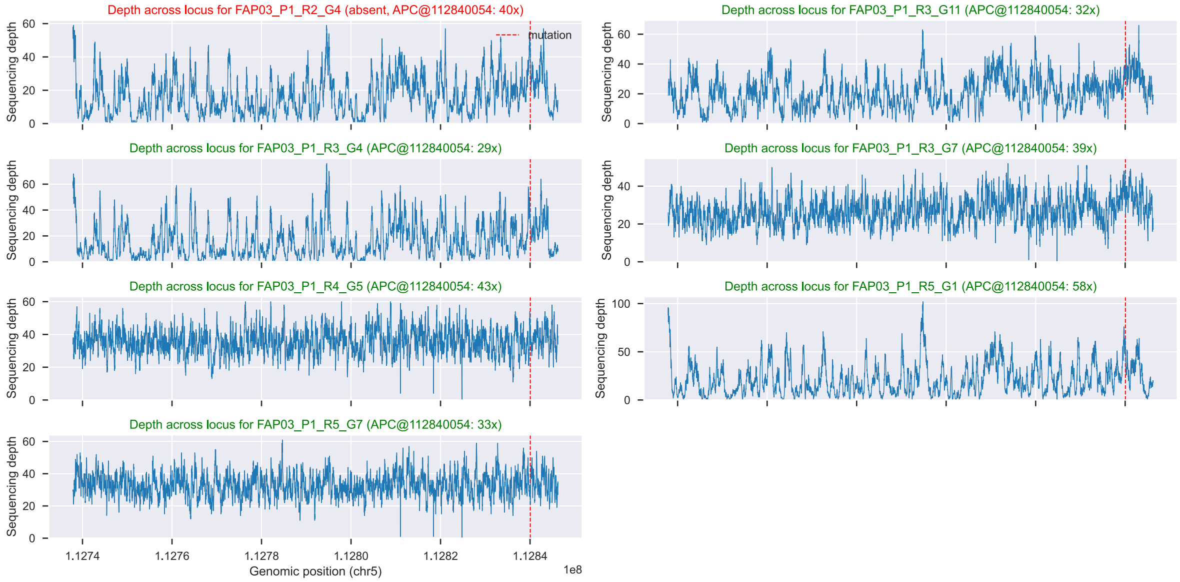

Figure SN3.48: Raw sequencing depth at the APC locus for samples from lesion FAP03_P1, with the APC mutation location highlighted. Green titles indicate samples in which the APC mutations passed our filters, and red titles indicate samples in which the APC mutation was never detected. In contrast to lesion FAP01_T3, the sample missing the APC mutation has high coverage at the mutation locus, making it unlikely that the mutation was missed due to low coverage.

### 6 Tree Inference Method Comparisons

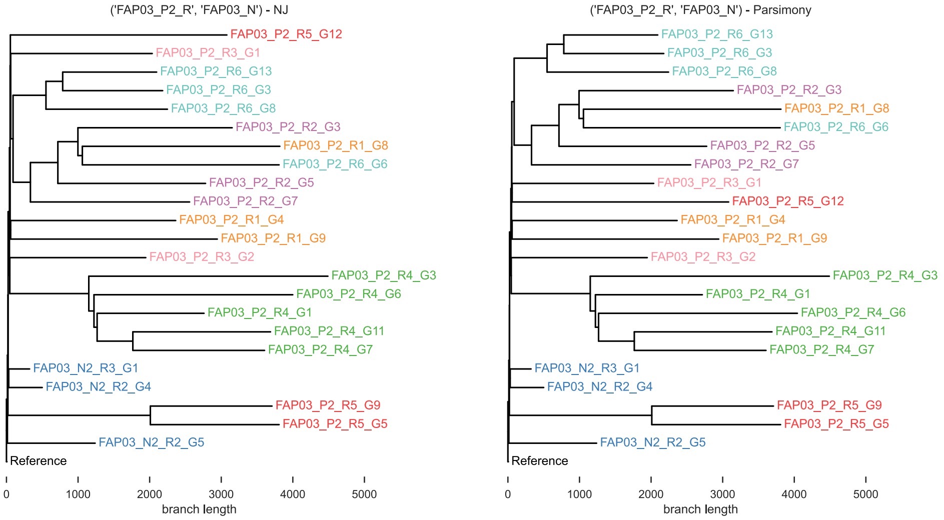

Figure SN3.49: Comparison of neighbor joining vs. maximum parsimony tree inference for lesion FAP03_P2.

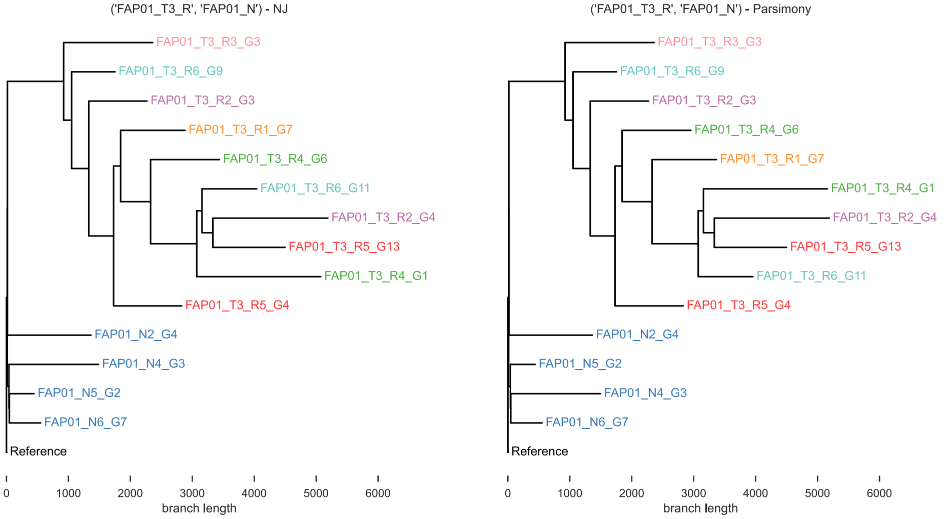

Figure SN3.50: Comparison of neighbor joining vs. maximum parsimony tree inference for lesion FAP01_T3.

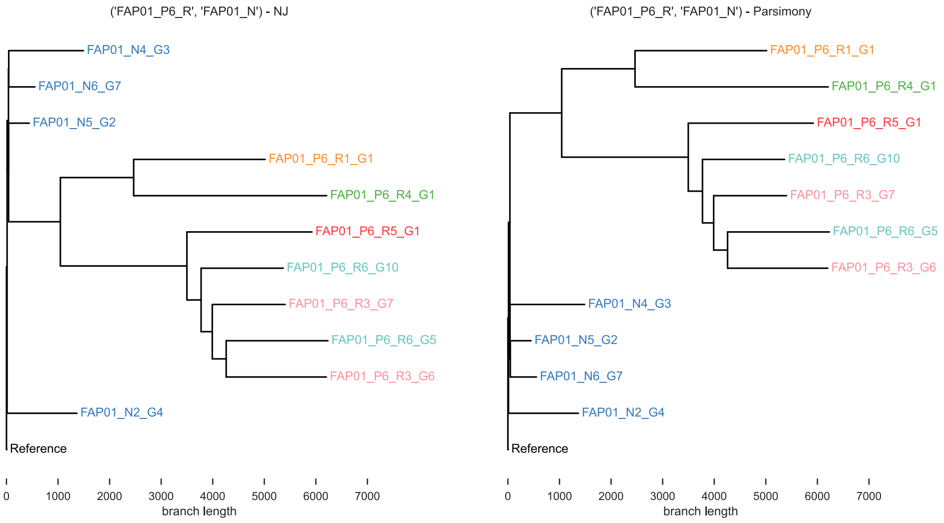

Figure SN3.51: Comparison of neighbor joining vs. maximum parsimony tree inference for lesion FAP01_P6.

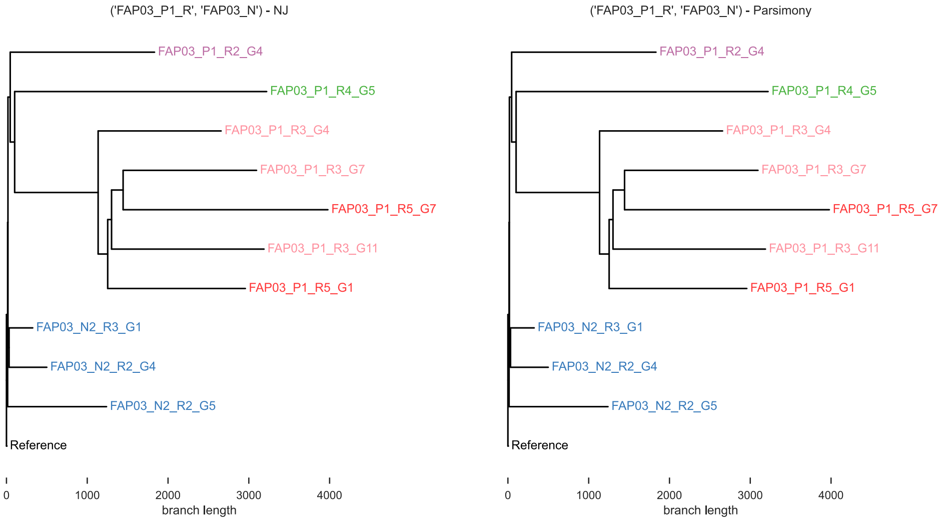

Figure SN3.52: Comparison of neighbor joining vs. maximum parsimony tree inference for lesion FAP03_P1.
